# Massively multiplex multimodal chemical screens at single-cell resolution

**DOI:** 10.64898/2026.05.23.727396

**Authors:** Kelvin Y. Chen, Romain Lopez, Basak Eraslan, Mayu Hata, Yusuke Takeshima, Katsuhiro Makino, Tatsuya Kibayashi, Kenji Ichiyama, Anne Biton, Jan-Christian Huetter, Anshul Kundaje, Jonathan K. Pritchard, Aviv Regev, Shimon Sakaguchi

**Author notes:** These authors contributed equally to this work (Kelvin Y. Chen, Romain Lopez, Basak Eraslan). Department of Computer Science and Biology, New York University, New York, NY, USA.

## Abstract

Recent applications of scRNA-seq for massively multiplexed chemical screens have enabled comprehensive profiling of drug responses at unprecedented scale and resolution. However, current assays remain limited to RNA readouts, lacking information on other phenotypic and mechanistic layers such as chromatin accessibility, protein abundance and post-translational modifications. Here, we introduce a scalable framework for multimodal chemical screens, combining parallel small-molecule perturbations with multimodal readouts. We extend existing experimental platforms into icCITE-plex and DOGMA-plex, enabling joint profiling of RNA, protein, and epigenomic responses to thousands of chemical perturbations in parallel. To systematically decode the regulatory circuitry underlying these responses, we develop MoCAVI, a contrastive analysis framework that disentangles the effect of small molecules from control variation in multimodal measurements, and PERCISTRA, a pipeline that infers causal links between chromatin accessibility and gene expression. Applied across ~410,000 primary T cells under ~2,800 conditions, our approach resolves compound-specific mechanisms, highlights off-target effects, and links chromatin accessibility changes to transcription factor networks in primary T cells. Our results establish a generalizable platform for profiling and analyzing cellular responses to chemical perturbations across multiple modalities.

## Introduction

High-throughput screens (HTSs) are fundamental to biological and therapeutic discovery^1^, yet most large-scale screening approaches continue to employ bulk measurements that ignore cell to cell state heterogeneity and may not be able to recover mechanisms of action (MoA) in cells. Advances in single-cell RNA seq (scRNA-seq) have enabled massively multiplexed small molecule (SM) drug screens, capturing thousands of drug-induced gene expression responses across diverse biological systems and contexts^2–4^. However, most studies remain limited to RNA profiling, lacking other modalities, such as protein and chromatin accessibility, which collectively determine cell state and are needed to decipher a molecule’s MoA.

Systematically understanding how specific compounds impact cellular phenotype requires monitoring their impact on diverse molecular species. Single cell multi-omics methods such as CITE-seq^5^, icCITE-seq^6^ and DOGMA-seq/TEA-seq^7,8^ couple scRNA-seq with measurement of protein or chromatin accessibility in the same single cell, but their application in chemical screens has been limited by challenges in scaling, multiplexing, data analysis and interpretation. First, scaling to thousands of perturbations across multiple modalities is constrained by both technical limitations and reagent costs, because many single cell technologies are based on droplets, but SM screens are executed in separate wells. Second, existing multiplexing strategies that affix oligonucleotide barcodes to cells or nuclei^2^ can compromise RNA quality, whereas other approaches^4,9,10^ are not optimized for large-scale screens, creating challenges in harmonizing data across plates or batches. Finally, analysis and interpretation remain challenging, as perturbations often induce subtle shifts in cellular state or gene programs that can be obscured by intrinsic heterogeneity and require disentangling the sources of variation, particularly when integrating multiple molecular modalities. Moreover, integration methods for joint chromatin accessibility and gene expression are conceived with observational data in mind. In contrast, perturbation datasets require tailored approaches that can identify transcription factors mediating drug effects by linking drug-induced changes in regulatory element accessibility to downstream gene expression responses.

T cells are a relevant case in point, as their activation is orchestrated by multiple signaling cascades that regulate post-translational modifications (PTMs), chromatin accessibility, transcription, and protein expression. This process is dysregulated in many human diseases, including autoimmunity and cancer^11–14^. Therefore, screening SMs for their ability to modulate T cell activation or effector function, as well as for unintended off-target effects, is highly desirable.

Here, we developed a generalizable multimodal chemical screening platform for interrogating drug-induced phenotypic responses across RNA, proteins, PTMs and chromatin accessibility in single cells. Our platform extends existing single-cell multi-omics protocols into icCITE-plex and DOGMA-plex, where thousands of independent perturbations are multiplexed in a single experiment through combinatorial barcoding with defined combinations of viral barcodes paired with hashing antibody-oligonucleotides. To decipher the regulatory effects of individual perturbations, we developed MoCAVI, a contrastive variational autoencoder that separates perturbation-induced changes from background cellular heterogeneity by generating distinct latent spaces for control variation and drug-specific responses. We further introduce PERCISTRA, to infer the causal mechanisms that drive phenotypic changes across chromatin accessibility and gene expression. We apply these approaches to primary T cells across ~410,000 single cells profiled under ~2,800 conditions, resolving compound-specific mechanisms of action and identifying dose-dependent on- and off-target effects and MoAs, demonstrating the broad applicability for high-content drug screening and mechanistic dissection across multiple regulatory and phenotypic layers.

## Results

To enable prospective multiplexing of thousands of independent samples in one experiment, we developed icCITE-plex, a tractable cellular barcoding system for high-content single-cell profiling experiments. We were inspired by previous studies utilizing viral-barcoded cells for lineage tracing^15–19^ and reasoned that a similar approach^9^ could be repurposed towards cellular hashing of multiple independent samples with a virally-encoded molecular readout. In icCITE-plex, chemical perturbation identity is encoded through defined combinations of well-specific viral barcodes and hashtag antibody-based plate-barcodes, followed by the icCITE-seq protocol^6^ (Fig. 1A and fig. S1A). This two-step barcoding approach provides scalable profiling from a single pool of cells and does not affect the data quality of the captured modalities.

**Fig. 1.**
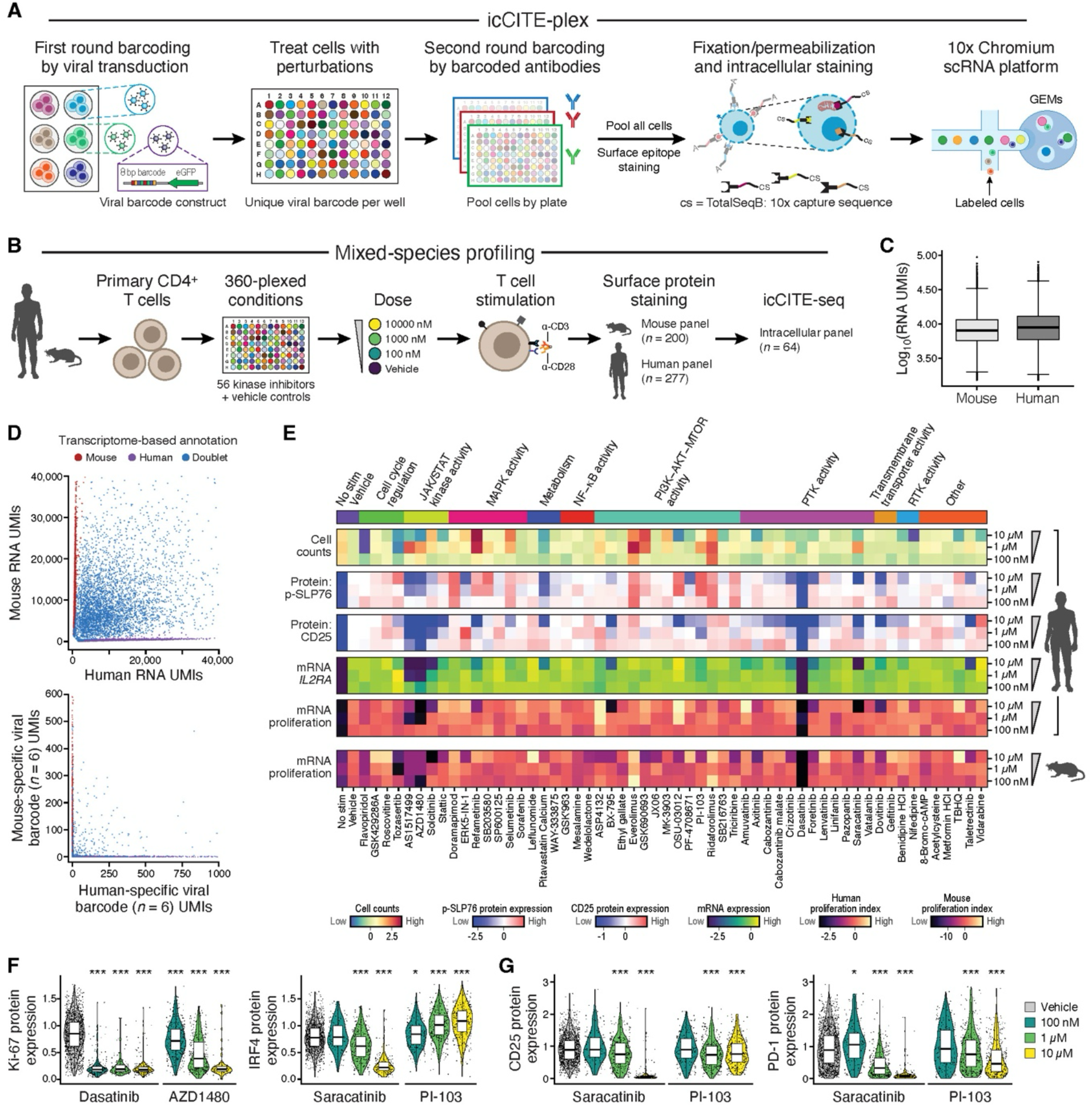
icCITE-plex enables multimodal chemical perturbation screens at single-cell resolution. (**A**) Experimental workflow of icCITE-plex. From left: Cells are barcoded with well-specific viral barcodes (left most), then subjected to SM treatment (second from left). Treatments are pooled by plate and labeled with plate-specific antibody-based hashtag oligonucleotides (center) and profiled by icCITE-seq (right panels). (**B-D**) Species mixing experiment and QC. (B) Experimental scheme. (C) Number of mRNA UMIs recovered per cell (*y* axis) for mouse and human cells (x axis). (D) Number of human (x axis) and mouse (y axis) RNA UMIs (top) and species-specific viral barcode UMIs (*n* = 6 per species, bottom) associated with each cell barcode (dots), colored by transcriptome assignment as human (purple), mouse (red), or doublet (purple). (**E**-**G**) Multimodal phenotypic readouts captured by icCITE-plex. (E) From top: Human cell count (viability), select protein RNA levels, and proliferation estimates, as well as mouse cell proliferation estimates (bottom) across SM treatments (columns) in decreasing doses (rows, from top). Top annotation bar: targeted pathway. (F,G) Distribution of selected intracellular (left) and surface (right) protein levels (y axis) across cells following indicated SM treatments (x axis), colored by dose (color bar) (two-tailed Mann-Whitney U-test, \**P* < 5×10^−2^, \*\*\**P* < 10^−3^). Boxplots (C,F and G) denote three quartiles, distribution with whiskers, and outliers as dots. Values plotted represent cells from a single replicate.

We first demonstrated the feasibility of our method in a species-mixing experiment with primary human and mouse T cells undergoing T cell activation and treated with each of 56 different kinase inhibitors in three different doses (100 nM, 1 µM and 10 µM). We added the inhibitors at the time of T cell activation and selected the doses to span a range of kinase inhibitor potencies^20^ (Methods). After 72 hours, we collected the cells and profiled them by icCITE-seq for RNA, 477 surface proteins, and 64 intracellular epitopes (Fig. 1B and fig. S1B). In total, we measured 360 drug-dose responses across 94,790 cells (table S1). For human and mouse single cells, we detected a median of 7,989 and 7,088 RNA unique molecular identifiers (UMIs); and 2,750 and 2,577 genes per cell, respectively (Fig. 1C and fig. S1C). Moreover, direct capture of polyadenylated viral barcode transcripts yielded 214 and 80 median viral barcode UMIs per cell, enabling robust assignment (fig. S1D and E). Species assignment by either the transcriptome, viral barcodes or hashtag antibodies yielded consistent results, validating the specificity of the method (Fig. 1D and fig. S1F,G).

Examination of changes across multiple single-cell modalities helped contextualize dose-dependent phenotypic effects (Fig. 1E). For example, profiles from cells treated with the tyrosine kinase inhibitor dasatinib largely resembled those of non-stimulated control cells, consistent with dasatinib’s well known negative regulatory effects on T cell activation^21^. These cells had lower RNA expression of proliferation marker genes, reduced levels of the Ki-67 protein, and lower abundance of activation markers such as CD25 and phospho-SLP-76 (Fig. 1E,F). Cells treated with the Src inhibitor saracatinib had a comparable profile, particularly at higher doses (Fig. 1E-G).

In contrast, treatment with the phosphoinositide 3-kinase (PI3K) inhibitor PI-103 caused a dose-dependent increase in phospho-SLP76 and in levels of the transcription factor (TF) IRF4 protein (Fig. 1E,F), indicating an increase in intracellular signaling. Interestingly, intracellular protein profiling revealed an unexpected increased percentage of FOXP3^+^ cells (the canonical Treg TF) in cells treated with the Aurora kinase inhibitor tozasertib (fig. S1H), as well as dose-dependent reductions in the levels of phosphorylated ribosomal protein S6 (Phospho-RPS6) in cells treated with PI3K inhibitors (fig. S1I,J).

Multiple SMs impacted cell abundance, estimated by total cell counts per drug-dose as a proxy readout for survival and proliferation. For example, there was a strong increase in the number of cells treated with the MEK1/2 inhibitor refametinib or the pan-AKT inhibitor GSK690693 compared to vehicle controls (Fig. 1E). This is surprising given the role of the MAPK and AKT pathways in cell growth, and may be due to the fact that their inhibition weakens T cell activation (see below), and thus may protect from activation-induced cell death. Conversely, the pan-CDK inhibitor flavopiridol strongly decreased cell numbers at all doses, perhaps reflecting compound-induced toxicity (Fig. 1E). Overall, 16.1% of compounds at 100 nM, 46.4% at 1 μM, and 62.5% at 10 μM impacted cell recovery (fig. S1K), highlighting the broad impact on cell viability and proliferation during T cell activation, especially at high doses which can involve non-specific effects.

Notably, treatment-induced changes in surface protein expression were overall well-correlated with concomitant changes in mRNA expression levels (fig. S1L). For example, changes in protein expression of CD25, Ki-67 and PD-1 were well correlated with their mRNA counterparts (*r* = 0.95; 0.86 and 0.91, respectively). However, we observed discordance between CD25 protein and mRNA (*IL2RA*) expression in cells treated with a high dose (10 µM) of the PDK1 inhibitor OSU-03012 (Fig. 1E and fig. S1L), perhaps indicative of negative regulation at the post-translational level due to off-target effects. Taken together, these findings establish the efficacy of icCITE-plex for the profiling of SM-induced phenotypes at single-cell resolution and underscore the utility of additional modalities beyond RNA profiles.

### Building multimodal perturbation maps via contrastive analysis

Analysis of icCITE-plex data across all cells, genes and proteins requires advanced computational methods^22,23^. We initially applied scVI^24^ and TotalVI^25^, established methods for analyzing scRNA-seq and single cell multimodal data, respectively. While these methods captured, at least partially, drug targets (Fig. 2A) and dosage (fig. S2A) effects, they also reflected additional cellular phenotypic heterogeneity. Specifically, both scVI and TotalVI separated cells by their cell cycle phase (Fig. 2A) even when examining cells exposed to the same drug (refametinib and GSK690693, fig. S2A). This separation may not be, however, directly relevant to drug effects, since activated T cells naturally proliferate, including in control conditions. This previously documented challenge^26–28^ compromises both interpretability and statistical power^27^, as perturbation effects on gene expression are often smaller than baseline phenotypic variation in the cell population^29^.

**Fig. 2:**
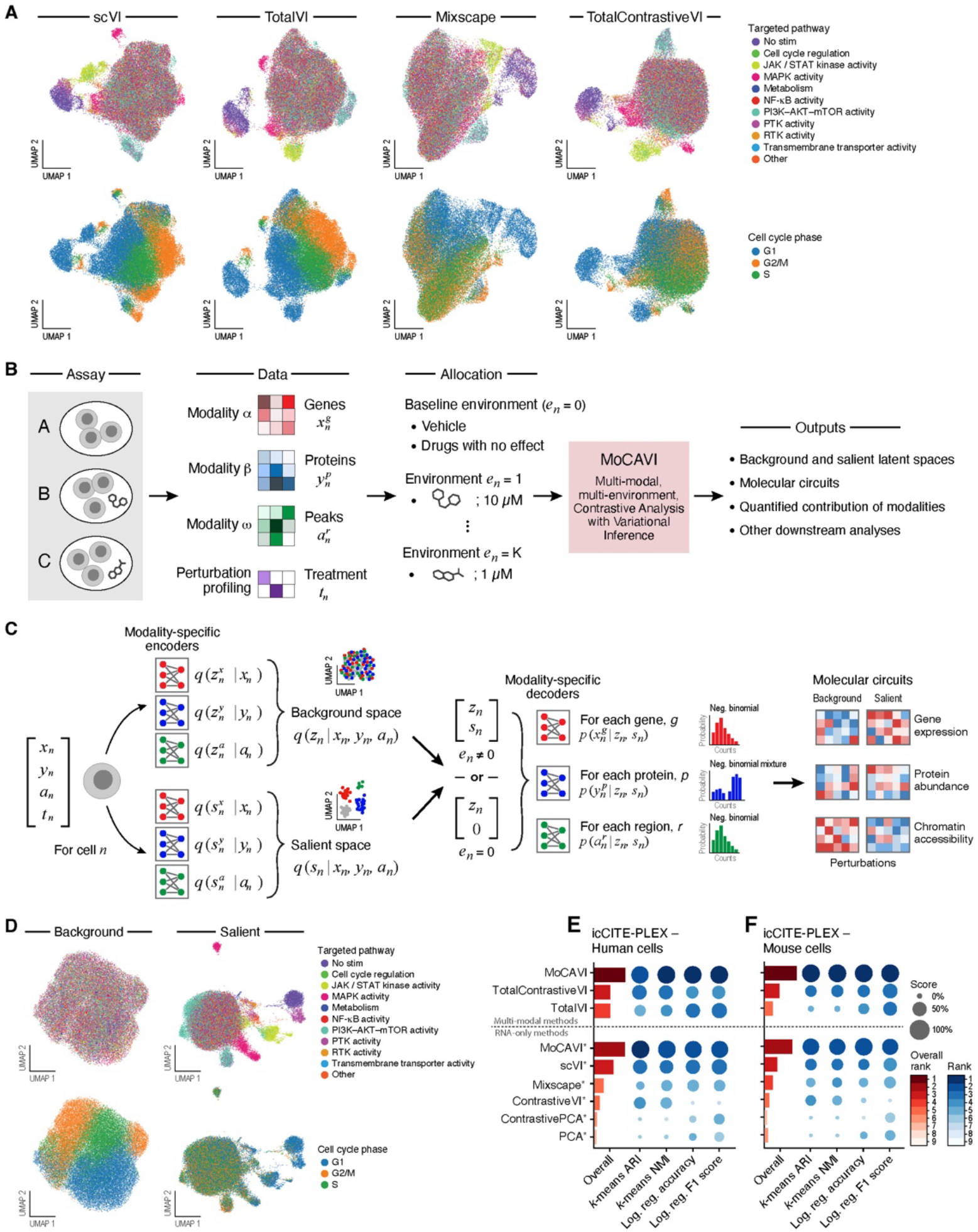
MoCAVI framework for multimodal, multi-environment contrastive analysis of chemical perturbations. (**A**) Existing single-cell perturbation analysis methods combine cell cycle and mechanism of action effects. Uniform Manifold Approximation and Projection (UMAP) representation of the latent space obtained via different embedding techniques (from left to right), with cells (dots) colored by the drug-targeted pathway (top) or by cell cycle phase (bottom). (**B-C**) MoCAVI. (B) Conceptual overview. From left: Chemical screens with single-cell readout (left most) produce multi-omics measurements (second from left) for each cell n that may include (any subset of) the expression level *x*_*n*_^g^ of gene *g*, the abundance *y*_n_^p^ of protein *p*, and the accessibility *a*_n_^r^ of chromatin region *r*, along with the treatment label *t*_*n*_ (i.e., a drug-dosage combination). MoCAVI takes as input these data and a partition of the treatment labels into environments (middle), with the baseline environment defined as the union of the control conditions (unperturbed) and treatments that have no estimated discernible effects (**Methods**). Each remaining treatment is assigned to its own environment. MoCAVI (second from right, pink box) infers the parameters of the generative models, and latent variables. These outputs (right) may be used for downstream analyses. (**C**) Generative model and inference scheme. From left: For each cell (left), data from each modality is passed through two separate sets of modality-specific encoder networks (second from left), one (top) encoding the parameters of the approximate posterior for the background latent variable *z*_*n*_ (capturing biological variation in the control population) and another (bottom) encoding the parameters of the approximate posterior for the salient latent variable *s*_*n*_. Those latent variables encode the biological variation that is most affected by the perturbations. Next, (second from right) modality-specific decoder networks map the latent variables [*z*_*n*_, *s*_*n*_] to the parameter of a count distribution for every feature captured by the modality (e.g., gene (top), protein (middle), or region (bottom)). For any cell in the baseline environment (*t*_*n*_ *= 0)*, the salient latent variable *s*_*n*_ is deterministically set to *s*_*n*_ *= 0* before feeding into the decoder networks. After training (right), the decoder networks may be used to characterize the effect of perturbations in both spaces, and across all modalities (**Methods**). (**D-F**) MoCAVI partitions cell cycle and mechanism of action variation. (D) UMAP representation of cell multimodal profiles (dots) in the background (left) and salient (right) latent spaces obtained via MoCAVI analysis, colored by drug-targeted pathway (top) or cell cycle phase (bottom). (E,F) Adjusted Rand Index (ARI), Normalized Mutual Information (NMI) after k-means clustering, accuracy and F1 score after logistic regression (columns) for assessing recovery of each drug’s mechanism of action based on the latent space obtained via each method (rows) in human (E) or mouse (F) data. The first three methods (top rows) use both protein and RNA measurements, all other methods (bottom, star) only rely on RNA measurements. Aggregate score (left column) was computed by averaging across all metrics, after scaling them to lie between 0 and 1. Higher values for all metrics indicate better performance.

To address these limitations, Mixscape^26^, as well as contrastive analysis methods (CA)^27,30–34^, such as (Total)ContrastiveVI^27^, were previously developed to isolate perturbation effects, while controlling for heterogeneity in the control population. When applied to icCITE-plex data, these methods increased mixing of cell cycle phases (Fig. 2A, Fig. S2A), but with two key limitations. First, they are limited in their ability to capture subtle differences between perturbations, because they only model all perturbed cells against control cells, but are agnostic to the identity of the specific perturbation applied to a cell. Second, they have limited multimodal capabilities (Mixscape operates on a single modality, while (Total)ContrastiveVI works only with gene expression or CITE-seq data).

We thus developed MoCAVI, a more sensitive approach that can handle any combination of modalities, including gene expression, protein measurements, and chromatin accessibility (Fig. 2B-C, and below). MoCAVI takes as input multimodal single-cell perturbation data, incorporating measurements across modalities and perturbation assignments (Fig. 2B). The input data are partitioned into environments: a baseline environment (control cells plus treated cells with non-significant gene expression changes, fig. S2B, Methods) and separate environments for each combination of drug and dosage. MoCAVI generates two distinct latent spaces and molecular maps: (1) a background latent space capturing control population heterogeneity, useful for identifying biological processes varying in control cells (e.g., cell cycle phases), as well as assessing how perturbations bias cells between these existing control states; and (2) a salient latent space that captures environment-specific variation, while controlling for background variation, enabling accurate perturbation classification based on mechanism of action (Fig. 2B, fig. S2C). The salient latent space also facilitates calculation of background-corrected log fold-changes (salient-specific LFCs) between perturbed and control cells (fig. S2D, Methods)—valuable when cell cycle or other heterogeneity dominates control population gene expression variation.

To model the multimodal data (Fig. 2C), the latent variable model posits probability distributions for counts from each modality following established approaches^24,25,35^: negative binomial for RNA expression and fragment counts of chromatin accessibility, as well as mixture of negative binomial for protein measurements. MoCAVI estimates each distribution’s parameters with neural networks, taking as input either both sets of latent variables for each non-baseline environment, or only the background latent variables for the baseline environment. Environment-specific conditional priors, inspired by causal representation learning^36^, enhance model sensitivity (demonstrated through ablation studies discussed below). We fit MoCAVI with amortized variational inference^37^ using dedicated encoder networks for each modality and latent space^38^.

Visualizing MoCAVI’s cell representation of the 56 kinase icCITE-plex screen (Fig. 2D and fig. S2E) shows that the background latent space successfully captured cell cycle phase variation, while the salient latent space effectively removed this effect and better highlighted drug impacts, including dose responses. MoCAVI outperformed competing methods on a quantitative benchmark, which measured how accurately salient embeddings reflected known drug mechanisms of action (information not provided to the model) (Methods; Fig. 2E,F).

To understand which aspects of the model were most critical for its performance, we systematically evaluated the contribution of each design component through ablation studies. We tested multiple model variants that differed in how they handled background cells, incorporated perturbation-specific information, and balanced the objective function of each environment (Methods; fig. S3). The best-performing configurations, including MoCAVI, combined rebalancing of non-affected cells with perturbation-specific modeling, indicating that these elements were key to accurately capturing perturbation effects.

### MoCAVI’s background space reveals T cell activation continuum

MoCAVI’s background representations of SM-treated cells showed minimal separation based on the class of SM (fig. S4A, right) or sequencing batch (fig. S4A, left), and a continuum of cell states rather than discrete clusters (fig. S4), consistent with previous profiling of T cells^39,40^.

To better interpret the biological processes underlying the basal heterogeneity in T cell activation, we applied Hotspot^41^ to identify gene programs that define distinct cellular states within the background space, revealing 11 background gene programs (bGPs) that represent key cellular processes in T cell activation (Fig. 3A,B and fig. S4B,C; table S2).

**Fig. 3.**
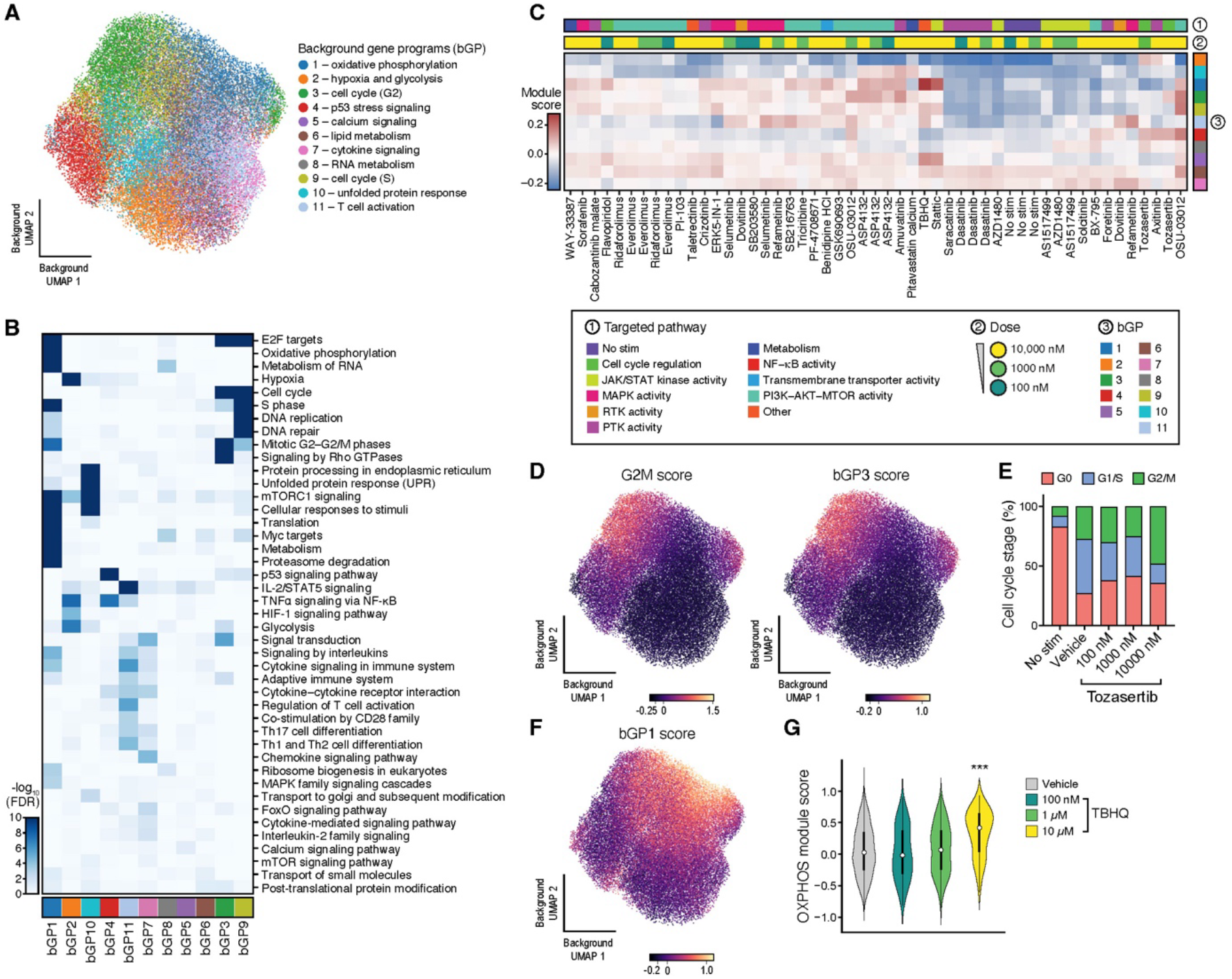
Background gene programs in MoCAVI’s background latent space. (**A-C**) Background gene programs (bGPs). (A) UMAP embedding of icCITE-plex multimodal profiles (dots) from MoCAVI’s background space, colored by the highest-scoring background gene program identified using Hotspot. (B) Pathways (rows) enriched (-log_10_(FDR), Fisher’s exact test, color bar) in each bGP (columns). (C) Scores (intensity color bar) for each bGP (rows, annotation bar 3) in the pseudobulk profiles (columns) for the SM treatments with the strongest impact in the background space, ordered by hierarchical clustering and color coded by drug target pathway (bar 1) and dose (bar 2). (**D-G**) Select functional associations of background gene programs. (D) UMAP embedding of MoCAVI’s background latent space for icCITE-plex multimodal profiles (points), colored by scores (color bar) for G2M cell cycle gene signature (left) or bGP3 (right). (E) Proportion of cells (y axis) assigned to each cell cycle phase inferred from gene expression following each tozasertib treatment (x axis). (F) UMAP embedding (as in D), colored by bGP1 scores (color bar). (G) Distribution of OXPHOS gene signature scores (y axis) for cells following TBHQ treatment (x axis) at each dose (color legend).

For example, bGP1 was enriched for genes involved in mitochondrial oxidative phosphorylation (including *ATP5F1B, SDHB, NDUFS7*, and *COX5A*), mTor signaling and other related pathways, suggesting enhanced activation-dependent cellular metabolism. Similarly, bGP2 was linked to genes associated with hypoxia as well as glycolysis (*SLC2A1, PDK1, HK2*, and *PGK1*), a key metabolic adaptation in T cells driven by hypoxia inducible factor 1 alpha (HIF-1α)^42^. On the other hand, bGP4 and bGP9 included genes involved in cellular stress responses, with bGP4 containing p53 signaling components (*MDM2, CDKN1A* and *GADD45A*) and bGP9 encompassing regulators of the endoplasmic reticulum (ER) stress-related unfolded protein response (*ATF4, HSPA5, DDIT3*, and *HSP90B1*). bGP7 and bGP11 were characterized by intracellular signaling and T cell effector gene expression, with bGP7 defined by genes such as *IL2RA, DGKE, PDE3B, SOS1*, and *GATA3*, whereas bGP11 comprised *PRF1, GZMB, CTLA4, BATF, LAG3* and *TNFRSF18*. As expected, the background latent space also captured variability in cell cycle states, with bGP3 and bGP9 enriched for G2-phase genes and S-phase genes, respectively (Fig. 3B,D).

Scoring drug-treated cells by the expression of each background program, some drugs impacted program usage in ways aligned with their annotated mechanisms of action (MoA) (Fig. 3C, fig. S4D). For instance, cells treated with the highest dose of the Aurora kinase inhibitor tozasertib had increased bGP3 scores, likely indicating a decrease in G1/S cells and an increase of G2-arrested cells, consistent with the role of Aurora kinase in controlling mitotic processes^43^ (Fig. 3C-E and fig. S4D). Conversely, treatment with a high dose of the NRF2 activator TBHQ resulted in the expected increase in a mitochondrial oxidative phosphorylation program (bGP1) and decrease in the glycolysis program (bGP2)^44^ (Fig. 3F,G and fig. S4D). High dose Stattic has a highly similar profile which may suggest some converging MoA. Likewise, the PI3K/AKT/mTOR pathway inhibitors PF-4708671, GSK690693, and triciribine also showed reduced expression of glycolysis programs, aligning with mTOR’s role in regulating the glycolytic metabolism switch in activated T cells^45^ (Fig. 3C and fig. S4D).

Together, these results highlight the diversity of biological pathways underlying T cell activation and illustrate how pharmacological manipulation may skew cells towards usage of specific gene programs and cellular states.

### MoCAVI’s salient representations uncover drug effects and molecular mechanisms

Quantifying drug effect sizes in the background and salient latent spaces showed that overall, drug effects in the salient space were detectable across a broader dose range, whereas effects in the background space were more skewed toward higher doses (Fig. 4A and fig. S5A-B). Drugs such as tozasertib, TBHQ and triciribine exhibited large effects in the background space (at high doses) but minimal ones in the salient space, reflecting an effect on the variations in shared cell states captured within the background space rather than leading to new phenotypic states that were not present in control cells. Higher doses of ERK5-in-1 and ASP4132 showed the opposite pattern, with large effects in the salient latent space but minor background effects, suggesting that these compounds predominantly drive new phenotypic states absent in vehicle control cells. Meanwhile, high dose treatments of drugs such as AZD1480 and dovitinib had substantial effects in both spaces, indicating impact on pre-existing states and the ability to elicit new phenotypic transitions. Thus, drug-induced phenotypes can be classified based on whether they primarily affect shared cellular processes captured in the background space, induce unique phenotypes represented in the salient latent space, or simultaneously influence both aspects.

**Fig. 4.**
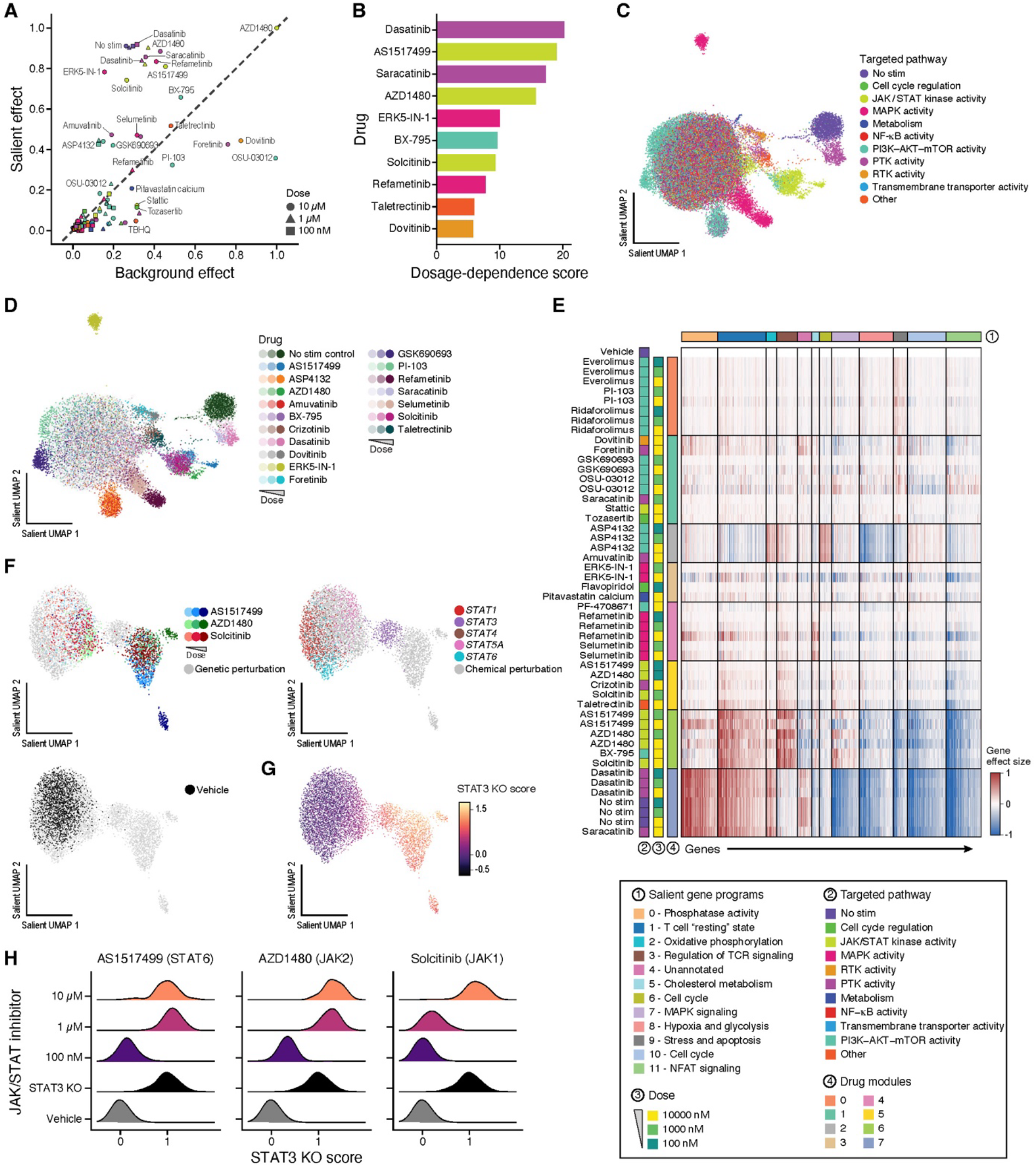
Salient gene programs in MoCAVI’s salient latent space. (**A**) Stronger salient than background effects for most SM perturbations. MoCAVI background (x axis) and salient (y axis) effect scores for each SM treatment (points; shape: dose; color: targeted drug pathway (color legend, C)). (**B**) Impactful treatments. Dosage-dependence scores (x axis) for each of top ten inhibitors (y axis, rank ordered). (**C**,**D**) Salient space distinguished perturbations by pathway and dose. UMAP embedding of MoCAVI’s salient representation of cell profiles (dots) for SM treatments, colored by targeted pathway (C) or SM identity and dose (color-intensity combination) (D). (**E**) Co-functional drug modules and co-regulated gene programs in the salient space. Regulatory effect sizes (color bar) for each drug (colored by pathway, bar 2) - dose (bar 3) combination (rows), grouped by module (bar 4) on the expression of each of the 2,500 most impacted genes (columns), grouped by salient gene programs (sGP, color bar 1). (**F-H**) Low dose JAK/STAT inhibitors phenotypically resemble STAT3 KO. UMAP embedding of MoCAVI’s representation of the joint salient latent space for select STAT family inhibitors and knockouts (dots), colored by drug treatment (F, top left, color bar), genetic perturbation (F, right, color bar) vehicle control cells (F, bottom left, black), or STAT3 knockout module scores (G, intensity color bar). (H) Distributions of STAT3 knockout module scores (x axis) in profiles of individual cells treated with increasing doses (rows) of AS1517499 (left), AZD1480 (middle) and solcitinib (right) or in cells with vehicle controls or STAT3 knockout.

To identify SMs with potent salient effects even at low doses, we calculated an average displacement score across dose increases, capturing the magnitude of embedding shifts as concentration increases, while accounting for cases where higher doses result in cell death (Methods). Dasatinib, AS1517499, saracatinib and AZD1480 ranked at the top drugs using this score, suggesting that these compounds produce substantial phenotypic changes even at lower concentrations (Fig. 4B).

Next, focusing on MoCAVI’s salient representation for small molecules (Methods), we observed that molecules targeting the same pathway tended to group together, indicating similar effects on molecular profiles (Fig. 4C and fig. S5C). For example, the MAPK pathway inhibitors selumetinib and refametinib had comparable dose-dependent phenotypes (Fig. 4C,D), and cells treated with JAK inhibitors solcitinib and AZD1480 co-localized in the salient latent space, suggesting a common MOA (Fig. 4D). In contrast, although cells treated with high doses of GSK690693 and PI-103 were relatively proximal in the overall salient latent space, they formed separate, discrete clusters (Fig. 4D). Since these inhibitors target different nodes of the PI3K-Akt-mTOR signaling pathway (AKT and PI3K/mTOR, respectively), these results potentially highlight how targeting different points within the same pathway may lead to divergent molecular phenotypes.

To better understand the molecular features driving the drug-induced phenotypic changes observed in the salient space, we applied MoCAVI to estimate the effects of small-molecule treatments on individual genes (Methods). The resulting regulatory model associated 49 small-molecule perturbations (molecule-dose combination) to 2,500 downstream genes, which could be further clustered into eight co-functional drug modules (Drug modules 0-7; table S3) and twelve coregulated gene programs (salient gene programs (sGPs) 0-11; Fig. 4E and fig. S5D,E; table S3). The twelve gene programs, each capturing a set of genes that responded similarly across the perturbations, were enriched for different immune and cellular processes. The co-functional drug modules consisted of sets of treatments (compound-dose combination) with similar effects on gene expression and were generally concordant with their annotated targeted pathways. Notable exceptions included crizotinib (targeting ALK, HGFR, c-MET, and ROS1), taletrectinib (ROS1 and NTRK), and BX-795 (PDK1), which exhibited phenotypic similarities to JAK/STAT pathway inhibitors solcitinib (JAK1) and AZD1480 (JAK2; Fig. 4E, modules 5 and 6). Both crizotinib and BX-795 have been previously shown to exhibit off-target activity against JAK/STAT *in vitro*^*46,47*^ and in cell-free systems^48^, but no prior literature evidence exists for taletrectinib (to the best of our knowledge), making this a potentially novel off-target annotation at high doses (10 µM). In another example, the multi-targeted inhibitor amuvatinib (c-Kit, PDGFRα, FLT3 and Met) and the AMPK agonist ASP4132 also exhibited similar effects. Also notable were the similarities between dovitinib (targeting FLT3, c-Kit, FGFR, and VEGFR) and foretinib (targeting HGFR, VEGFR, and Met), both of which strongly impacted RNA profiles despite the absence of expression for their annotated target proteins in T cells.

A similar analysis using only protein data revealed relationships between perturbations that were consistent with those inferred from RNA profiles (fig. S6A; Mantel distance = 0.95, *p* < 10^−5^). This suggests that protein data alone could be sufficient for the model to learn the global relationships between treatments. To obtain a more refined understanding of which modality contributes most to a cell’s representation we calculated modality-specific weights, a feature unique to MoCAVI (Methods). The distribution of weights (fig. S6B,C) suggested that MoCAVI primarily relies on RNA expression, except for a specific part of the latent space containing non-stimulated control and non-stimulated-”like” cells (fig. S6D), which is explained by naïve T cell protein markers CD62L and IL7R, as well as the level of activation markers such as CD69, GITR, CD25, and PD-1 (fig. S6E). We hypothesize that those could be clearer markers than RNA levels for identifying T cell activation states, aligning with their well-established use. Note, however, that while protein measurements robustly capture major phenotypic differences such as naïve versus activated T cell states, they cover far fewer features than RNA, which limits the granularity of the regulatory programs affected by SM treatment. Therefore, while protein-based representations are useful for identifying and understanding phenotypic relationships, fine-scale mechanism-level effects are better captured by global profiles, such as those available from RNA data.

The parsing of individual salient gene programs contextualized the impact of compounds on specific cellular processes, such as naive/resting T cell states (sGP1), cell cycle (sGP6 and sGP10; see Discussion), regulation of T cell signaling (sGP0, sGP3, sGP7 and sGP11), metabolism (sGP2, sGP5 and sGP8) and regulation of cellular stress/apoptosis (sGP9) (Fig. 4E). For example, perturbation by compounds in Drug Module 6, which primarily consists of JAK/STAT inhibitors, led to higher expression of naive T cell programs (sGP1; *BACH2, LEF1, FOXP1, TCF7*, and *SELL*) and a decrease in cell cycle progression (sGP10; *MCM2, E2F1, PCNA*, and *CDC6*) at lower doses. Concurrently, the treatments show some expression of T cell activation related programs, at levels lower than vehicle but higher than unstimulated cells, including elevated expression of downstream T cell signaling effectors (sGP3; *NFATC1, REL* and *NR4A2*) and enhanced expression of effector gene programs (sGP7; *TNFRSF18, TNFRSF4, CTLA4, MYB* and *NFKBIZ*) (Fig. 4E). These findings suggest that while JAK/STAT inhibition promotes a more ‘naïve-like’ cell state, it does not completely suppress activation. Changes in protein levels supported this conclusion, with increased levels of CD62L, CD69, GITR, 4-1BB, CD39 and CD45RO, and reduction in CD71, CD25, and PD-1 levels compared to vehicle control cells (fig. S6E). This effect was enhanced in higher concentrations of AS1517499 and AZD1480, with higher levels of CD3, CD45RA and IL7RA, and further reduced levels of activation markers CD69 and CD109 (fig. S6E). Along with the abrogation of phospho-SLP76 (Fig. 1E) and the decreased expression of T cell signaling effector sGP7 genes (Fig. 4E and fig. S7A), these observations suggest a complete shutdown of active TCR signaling at sufficiently high doses. Drug Module 4, which includes the MEK inhibitors refametinib and selumetinib, exhibited a similar, albeit much weaker, phenotype at the protein level (fig. S6E), particularly at higher concentrations. However, its RNA profile differed (except for the highest dose refametinib), showing mostly strong induction of genes in cholesterol biogenesis and lipid uptake (sGP5; *FDFT1, DHCR7, SREBF1, INSIG1 and OSBPL3*) (Fig. 4E and fig. S7A), along with increased phospho-SLP76 (Fig. 1E), suggesting stronger T cell activation. Indeed, these characteristics, including increased CD62L protein, decreased CD44 protein, increased expression of lipid uptake (sGP5) and ‘naive-like’ (sGP1) programs, and reduced expression of glycolysis programs (sGP8), were consistent with the T_SCM_-like phenotype reported for MEKi in CD8 T cells^49^.

Compounds in Drug Module 0, consisting exclusively of PI3K/mTOR/Akt inhibitors, induced the expression of genes involved in regulation of apoptosis (sGP9) (Fig. 4E and fig. S7B), including anti-apoptotic genes such as *BCL2, MYC, MDM2*, and *MDM4*, which may reflect enhanced T cell survival and resistance to activation-induced cell death. This interpretation is further supported by increased total recovered cell counts, particularly for higher doses of ridaforolimus and everolimus (Fig. 1E). Similarly, the PDPK1 inhibitor OSU-03012 in Drug Module 1 upregulated anti-apoptotic genes *BCL2* and *MDM2* but also strongly induced the expression of pro-apoptotic genes (also in sGP9), *BAX and FAS*, as well as Fas protein expression (fig. S7B,C and fig. S6E), suggesting these cells were more prone to cell death. Concomitantly, increasing OSU-03012 concentrations led to lower numbers of recovered cells (Fig. 1E), highlighting the divergent effects of targeting different nodes within the PI3K pathway on T cell survival. This demonstrates how integrating multiple data layers facilitates interpretation of phenotypic outcomes following SM perturbation.

Interestingly, cells treated with the STAT6 inhibitor AS1517499 exhibited a phenotype similar to those in cells treated with the JAK family inhibitors AZD1480 and solcitinib (Fig. 4E, drug modules 5 and 6). While previous studies reported that AS1517499 specifically inhibits STAT6 phosphorylation via an unknown mechanism at 100 nM^50^, even at this dose AS1517499 aligned with the effects of lower concentrations of AZD1480 and solcitinib within drug module 5, suggesting that these compounds may share a common MoA, rather than reflecting a high-dose non-specific effect (Fig. 4E and fig. S6E). To investigate this hypothesis further, we compared JAK/STAT inhibitor-treated cells with targeted knockouts of STAT family transcription factors (Methods). Co-embedding RNA profiles of genetic and chemical perturbations into a single latent space revealed that while STAT3 knockout induced a strong shift in expression profile, other STAT knockouts had minimal effects (Fig. 4F). Inhibitor treatment drove a dose-dependent shift toward STAT3 KO cells and beyond, with low doses approaching the STAT3 KO state and higher doses driving the cells to an even more extreme expression state (Fig. 4F-H and fig. S7D). These findings suggest that JAK/STAT inhibitors exert their effects in this context, at least in part, by suppressing STAT3 activation and that AS1517499 may also exploit this mechanism.

### Uncovering drug mechanisms through integration with genetic perturbation data

Inspired by the potential utility of integrating genetic and chemical perturbation datasets to uncover MoA^51,52^, we next used MoCAVI to combine our kinase inhibitor screen dataset with a Perturb-icCITE-Seq dataset following CRISPR/Cas9 KO of each of 100 genes encoding proteins in the same biological system (Methods; Fig. 5A)^6^. A co-embedding of the chemical and genetic perturbation datasets revealed both discrete clusters representing genetic- and chemical-specific phenotypes, and regions of convergence that suggested shared mechanisms (top 50 strongest perturbations across chemical and genetic data appear in Fig. 5B and fig. S8A,B). As a positive control, the knockout of *CD3E*, a major mediator of T cell signaling transduction correctly co-localized with non-stimulated control and dasatinib-treated cells (Fig. 5B “1”). Similarly, MEK1/2 inhibitors refametinib and selumetinib are positioned closely with the knockout of their downstream effector *MAPK1/3* (ERK1/2; Fig. 5B “2” and fig. S1I), indicating successful integration.

**Fig. 5.**
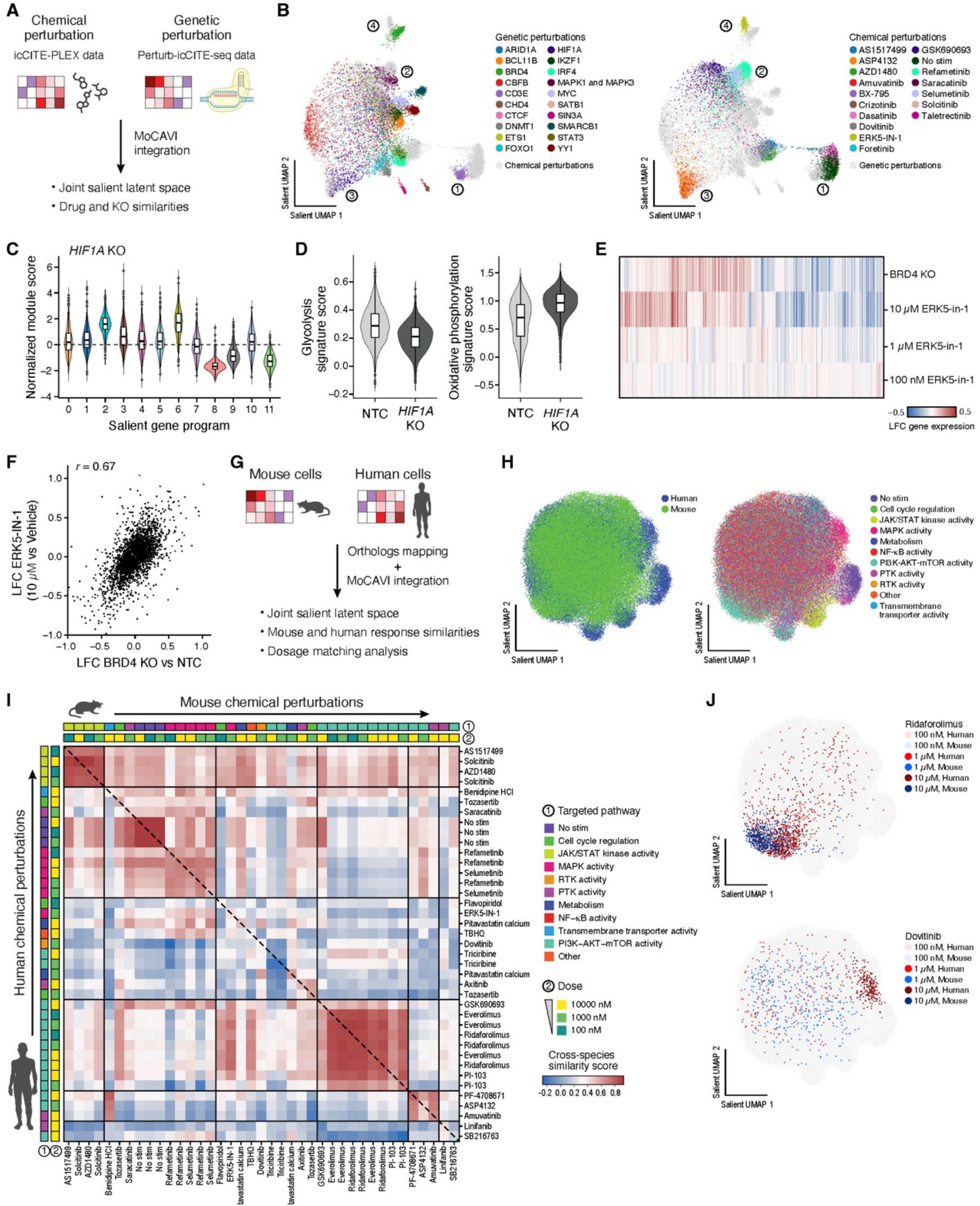
MoCAVI enables joint modeling of genetic and chemical perturbations and cross-species analyses. (**A**,**B**) Linking chemical and genetic perturbations. (A) Schematic workflow. (B) UMAP embedding of MoCAVI’s representation of the joint salient latent space for each multimodal profile (dots), colored by genetic (left, color bar) or chemical (right, color bar) perturbation identity. Numbers indicate points of interest. (**C**,**D**) Salient gene program activity in HIF1A-KO cells. (C) Distribution of normalized module scores (y axis) for each salient gene program (x axis) in *HIF1A* KO cells. (D) Distribution of gene signature scores for glycolysis (left, y axis) and oxidative phosphorylation (right, y axis) pathways in non-targeting control (NTC) and HIF1A knockout cells (x axis). (**E**,**F**) Similar RNA expression responses to BRD4 KO and ERK5-IN-1 inhibitor. (E) Log fold-change (LFC, color) in the expression of each gene (columns) under each perturbation (rows) relative to control (Vehicle or NTC). (F) Log fold change (y and x axes) in expression of each gene (dot) in BRD4 knockout vs NTC (x axis) and ERK5-IN-1 vs Vehicle (y axis). (**G-J**) Comparison of drug effects between human and mouse. (G) Integration approach using MoCAVI. (H) UMAP embedding of icCITE-plex multimodal profiles (dots) in the cross-species salient latent space, colored by species (left) or drug target pathway (right). (I) Cosine similarity (color) between pseudobulk profiles of each SM treatment (rows, columns, annotated by pathway (bar 1) and dose (bar 2)) computed from the MoCAVI joint salient space for human (lower triangle) and mouse (upper triangle) cells. (J) UMAP embedding as in (H) highlighting cells treated by ridaforolimus (top) and dovitinib (bottom) in each species and dose (color legend).

Cells treated with the AMPK agonist ASP4132 or amuvatinib were similar to those with *HIF1A* knockout (Fig. 5B “3” and fig. S8B), with a decrease in the expression of a glycolysis program (sGP8) and an increase in an oxidative phosphorylation program (sGP2) (Fig. 5C,D and fig. S8C). This metabolic rewiring is consistent with the known role of HIF1α in promoting glycolysis while suppressing fatty acid oxidation (FAO)^53^, and the established function of AMPK agonism in enhancing FAO metabolism in activated T cells^54^. These results therefore suggest that both pharmacological activation of AMPK and genetic ablation of HIF1α converge on similar metabolic pathways, potentially offering multiple strategies to modulate T cell metabolism. Note that the multi-target inhibitor amuvatinib (c-Kit, PDGFRα, FLT3 and Met) has a nearly identical profile but we cannot determine if this is through an indirect or an off-target effect, given its low specificity.

Lastly, there is a striking juxtaposition of cells treated with ERK5-in-1 and those with *BRD4* knockout (Fig. 5B “4” and fig. S8B). An independent analysis revealed a dose-dependent similarity at the RNA level, suggesting that BRD4 may be a target of ERK5-in-1 treatment, particularly at higher concentrations (Fig. 5E,F and fig. S8D,E). Prior studies have reported off-target binding *in vitro* between ERK5-in-1 and bromodomain-containing proteins, including BRD4, based on a comparison of BET and ERK5 inhibitors^55,56^, but our analysis provides the first evidence of functional equivalence between ERK5-in-1 treatment and *BRD4* genetic ablation. These results illustrate the utility of our integrative framework in uncovering and characterizing off-target effects, which could be broadly applicable to other chemical inhibitors for more precise elucidation of drug mechanisms and systematic probing of off-targeting activity.

### Cross-species analysis reveals conserved and divergent drug effects

Leveraging the cross-species nature of our dataset, we compared compound-induced phenotypes between mice and humans. To this end, we co-embedded mouse and human primary T cell RNA profiles into a shared latent space with MoCAVI (Methods, Fig. 5G). While drug responses were largely conserved across species, certain treatments led to notable differences (Fig. 5H,I and fig. S9A). These differences were not restricted to drugs targeting only one pathway, such that human/mouse differences were observed for multiple drug classes. For example, non-stimulated control cells and those treated with varying doses of ridaforolimus exhibited consistent RNA profiles across species compared to respective vehicle-treated controls (Fig. 5I,J top, and fig. S9B,C). Conversely, human and mouse T cells exposed to dovitinib (Fig. 5J bottom and fig. S9D) or triciribine (fig. S9E) displayed distinct RNA profiles, suggesting that these drugs may affect human and mouse cells through different mechanisms or sensitivities. In some cases, mouse cells had reduced viability, especially at higher drug doses, compared to human cells (fig. S9F). This discrepancy may reflect differences in drug effective doses or in off-target effects between species, especially with molecules more optimized to human protein targets. Consistently, mouse cell profiles at a given dose were frequently most similar to the human cell profiles at a lower dose of the same drug (*p* < 0.05; permutation test, Methods, fig. S9G). Consequently, doses that remain tolerable in human cells may be toxic in mice, potentially due to differences in drug metabolism, uptake, downstream signaling, or cell size. These results suggest that intrinsic differences between human and mouse cells may need to be accounted for when interpreting comparative drug response data.

### RNA, chromatin, and protein profiling in drug-treated cells with DOGMA-plex

Single-cell multi-omic profiling of RNA and chromatin accessibility can help better understand gene regulation^57–59^, and, for chemical perturbation screens, help better decipher how compounds impact gene expression. We therefore extended our chemical screening platform to DOGMA-plex, by co-profiling RNA, chromatin accessibility and select proteins using DOGMA-seq^7^.

We applied DOGMA-plex to a panel of 290 epigenetic inhibitors measured at four doses with two independent biological replicates in primary human CD4^+^ T cells undergoing activation (Fig. 6A,B), to a total of 2,400 multiplexed conditions, in 314,329 high-quality cell profiles (277 cells per molecule-dose condition on average, Fig. 6C; fig. S10A, Methods). Several compounds, specifically dBET57, dacinostat, JIB-04, CUDC-907, BI-2536, chaetocin, panobinostat, 2’-deoxy-5-fluorocytidine, and dBET6 substantially reduced cell viability at higher concentrations, leading to fewer than 50 detectable cells in at least 2 drug-dose conditions (fig. S10A). On a per cell basis, there was a median of 9,228 RNA UMIs, 3,344 detected genes and 11,768 ATAC fragments mapping to the nuclear genome (fig. S10B-D). Across compounds, the extent of impact was relatively consistent across modalities (fig. S10E). Indeed, inhibitors that induced widespread RNA changes also tended to broadly alter chromatin accessibility and protein expression, suggesting coordinated regulation across multiple layers (fig. S10E).

**Fig. 6.**
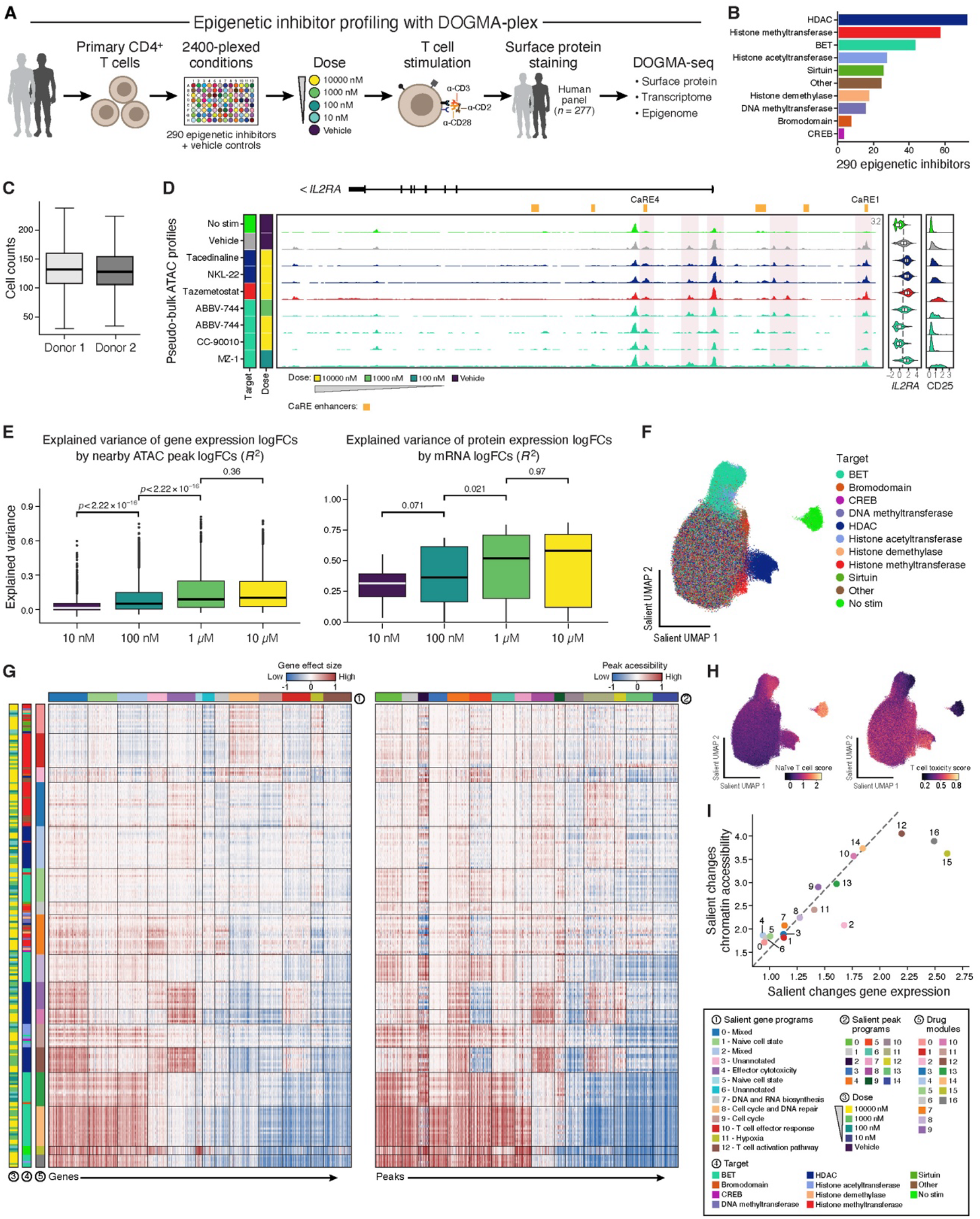
Joint gene expression, chromatin accessibility and protein profiling of chemical screens with DOGMA-plex. (**A**,**B**) DOGMA-plex experiment. (A) Overview. (B) Drug target distribution of the 290 epigenetic inhibitors in the DOGMA-plex experiment. (**C**) DOGMA-plex quality metric. Number of cells recovered per drug-dose condition (y axis) from each donor (x axis). Boxplots denote three quartiles, distribution with whiskers, and outliers as dots. (**D**) Chromatin regions differentially accessible between drug-treated and vehicle cells at the *IL2RA* (CD25) locus. Pseudo-bulk ATAC signal genomic tracks in each drug-dose combination (rows) at the *IL2RA* locus along with corresponding distribution of normalized *IL2RA* RNA and CD25 protein counts (right panels, x axis). CaRE enhancers annotated. Drug target (as in (B), left) and dose (color bar, left) are indicated. (**E**) Detection of propagated effects from chromatin to RNA to protein increases with dose. Distributions (three quartiles, distribution with whiskers, and outliers as dots) of the variance explained (y axis) in gene expression log fold-changes by accessibility log fold-changes of nearby peaks (left), and in surface protein log fold-changes by the corresponding gene’s mRNA log fold-changes (right), across drug treatments in each dose (x axis). Group differences were assessed using one-sided paired T-tests \**P* < 5×10^−2^, \*\**P* < 10^−2^, \*\*\**P* < 10^−3^, p ≤ 0.1, ns = not significant. (**F**,**G**) MoCAVI representation of the salient latent space for select drug perturbations. (F) UMAP of cell profiles (dots) in MoCAVI’s salient latent space, colored by drug target. (G) Regulatory effect sizes (color) for each drug (bar 4) - dose (bar 3) treatment (rows), grouped by modules (bar 5) on the expression of the 2,500 most impacted genes (left, columns), grouped by salient gene programs (bar 1) or 47,345 impacted peaks (right, columns), grouped by salient peak programs (bar 2). (**H**) BET and HDAC inhibitors impact distinct T cell programs. UMAP representation (as in G) colored by gene signature scores (color bar) of T cell “naiveness” (left) or cytotoxicity (right). (**I**) Some drug modules are particularly impacting the RNA vs. chromatin level. Changes in gene expression (x axis) and chromatin accessibility (y axis) effect scores (**Methods**) for each drug module (points), colored by the drug target (color bar, G).

We analyzed the impact of SM treatment on RNA expression within the context of the associated chromatin states. For example, cells treated with high doses of BET inhibitors ABBV-744 and CC-90010 had a strong reduction in chromatin accessibility at select regions in the *IL2RA* locus (by pseudobulk) with a concomitant decrease in both expression of *IL2RA* mRNA and CD25 protein (Fig. 6D). These included changes in accessibility in the previously characterized *IL2RA* CRISPRa-responsive element 1 (CaRE1) and in activation-dependent enhancers immediately upstream and downstream the *IL2RA* transcriptional start site (TSS)^60^. Notably, there were no significant changes in the TCR stimulation-responsive enhancer CaRE4, suggesting that the failure to upregulate *IL2RA* was not solely a consequence of impaired signal transduction following TCR stimulation^7,60^ (Fig. 6D). In contrast, cells treated with HDAC inhibitors tacedinaline and NKL-22 had increased *IL2RA* and CD25 expression, with largely unchanged chromatin accessibility. This suggests that an increase in local histone acetylation may suffice to promote transcriptional activation.

We systematically assessed how much of the variability at each regulatory layer can be attributed to variation at a preceding upstream step, by analyzing relative changes in chromatin accessibility, mRNA expression, and protein abundance across a range of drug-dosage conditions. Overall, the extent to which variance in mRNA log fold-changes is explained by chromatin accessibility at nearby genomic loci increased steadily with higher drug concentrations (Fig. 6E, left; median R^2^ values of 0.01, 0.05, 0.09, and 0.10 for 10 nM, 100 nM, 1 µM, and 10 µM, respectively). However, this relationship varied widely across individual genes. For some genes—such as *IL1RAP, AMN1, SYTL3, INTS7, STAT1, IFNG*, and *DLEU2*—more than 75% of the variance in mRNA levels was explained by chromatin accessibility changes at proximal loci, while for other genes, this proportion approached zero. For high doses, this explained variance only weakly correlated with the number of perturbations that affected each gene (Pearson’s *ρ* = 0.11 (*P* < 8.9×10^−7^) and 0.20 (p < 2.2 × 10^−16^) for 1 µM and 10 µM, respectively; fig. S10F), suggesting that this variance is not solely explained by the overall magnitude of the perturbation effect but instead by regulatory rules. Next, for genes with both RNA and protein measurements, mRNA LFCs explained a substantial proportion of the variance in the corresponding protein’s LFC, and increasingly at higher doses (Fig. 6E, right, median *R*^2^ = 0.31, 0.36, 0.51, and 0.58 for 10 nM, 100 nM, 1 µM, and 10 µM, respectively). Thus, as the number of significant effects rises with increased dosage, it sharpens the regulatory relationship from chromatin accessibility to mRNA levels to protein expression.

### Distinct RNA and epigenetic programs underlie BET and HDAC inhibitor responses

Integrating the impacted responses across chromatin accessibility, RNA features, and protein levels, many compounds are grouped by their drug families in MoCAVI’s salient space, including BET, HDAC and HMT inhibitors, highlighting distinct patterns of perturbation characterizing each family (Fig. 6F and fig. S11A-C). We estimated the effects of SM treatment on individual genes and chromatin peaks, associating 220 strong-effect drug-dose combinations (fig. S12A,B, Methods) to 3,000 and 47,345 significantly changed genes and peaks, respectively, and clustering those into seventeen co-functional drug modules (Drug modules 0-16), thirteen co-regulated gene programs (salient gene programs 0-12) and fifteen co-regulated chromatin peak programs (salient peak programs 0-14) (Fig. 6G and fig. S12C).

There was strong concordance between drug modules and their annotated inhibitor class. For example, five drug modules (#5, 8, 13, 14, and 16) are near-exclusively comprised of BET inhibitors, suggesting that while there are intra-family differences in the impact of individual compounds at specific doses, they still share similarities that distinguish them from other drug classes (Fig. 6G). In particular, BET inhibitors associated with drug modules 13, 14 and 16 epigenetically and transcriptomically resembled a ‘naive-like’ phenotype (sGP1 and sGP5), with upregulated expression of naïve-like genes such as *FOXP1, LEF1*, and *TCF7*, along with associated chromatin changes (Fig. 6G,H and fig. S12D top). Notably, drug modules 9, 10 and 12 comprise exclusively of HDAC inhibitors and were associated with induction of the T cell cytotoxicity gene programs (sGP4 and sGP10), with upregulation of RNA and corresponding chromatin accessibility of genes such as *IFNG, GZMB, PRF1 and HAVCR2* (Fig. 6G and fig. S12D bottom). This phenotype is reminiscent of highly activated effector cells with upregulation of a cytotoxicity program (Fig. 6H, right). While previous reports identified HDAC3 as a negative epigenetic regulator of a RUNX3-driven CD8^+^ T cell cytotoxic program^61^, our findings suggest that this regulatory role may extend beyond HDAC3 and include additional HDAC isoforms, at least in CD4^+^ T cells.

The distribution of modality-specific weights across the dataset suggests that MoCAVI predominantly leverages gene expression and chromatin accessibility for constructing cell representations, possibly because the feature space of these modalities is more complex than that of the select measured proteins (fig. S13A). In four drug modules (#2 (mixed), 12 (high dose HDACi), 15 (no stimulation) and 16 (high dose BETi)) MoCAVI relied on RNA levels more heavily than the average (Fig. 6I). Drug modules 12, 15 and 16 have large effects in each modality (Fig. 6G,I), simultaneously affecting many peaks and genes. (Note that modules 15 and 16 are small modules.) Drug module 2, however, showed a moderate effect on the transcriptome and a weak effect on chromatin accessibility, with the upregulation of cell cycle and DNA replication genes in sGP8. This is consistent with the well-established fact that chromatin accessibility is less sensitive to cell cycle changes^58^. Four drug modules (#0 (mixed), 6 (mixed), 4 (low dose HDACi), and 5 (lose dose BETi)) were more explained by chromatin accessibility than RNA levels (Fig. 6I). We hypothesized that at lower doses, chromatin accessibility may better describe the effect of those BET and HDAC inhibitors than gene expression.

To corroborate this, we compared total log-fold-changes in RNA and ATAC for each drug-dose combination and trained linear discriminant analysis classifiers in each modality using as input the multi-modal perturbation classes (weak-versus strong-effect; fig. S12B, Methods). This analysis projects the global perturbation classes into modality-specific spaces. Using this framework, BET inhibitors such as OTX015 and CPI-203 from drug module 5 showed weak effects on gene expression at lower doses, but strong effects on chromatin accessibility, and became strong in both modalities at higher doses (fig. S13B). This chromatin-first pattern was significantly enriched in drug module 5 compared with other clusters (Fisher’s exact test, odds ratio = 22, P < 10-3).

We also reasoned that BET inhibitors disrupt bromodomain-mediated reading of acetylated histones^62^, and thus it is possible that at low doses they preferentially affect chromatin accessibility prior to downstream RNA changes. Supporting this hypothesis, the proportion of peaks whose nearest gene was DE only at higher dosage (out of differentially accessible peaks at the lower dose) was three-fold higher in BET than HMT inhibitors (fig. S13C, Kolmogorov-Smirnov test, *P* < 10^−3^, Methods).

### A multi-step framework for decoding perturbation-induced regulatory networks

To gain further mechanistic insights into how epigenetic inhibitors impact the observed gene programs, we developed PERCISTRA, an analysis pipeline that infers TFs that mediate perturbation effects by regulating target genes through accessible binding sites (Fig. 7A). While many established single cell multiomics data analyses approaches rely only on cell-to-cell variability in observational data^63^, we leveraged our perturbation data to uncover putative causal relationships between TFs whose binding activities are impacted by the perturbations and their target gene expression, both across and within each drug-dosage condition. This duality enabled the reconstruction of regulatory networks responsive to perturbations.

**Fig. 7.**
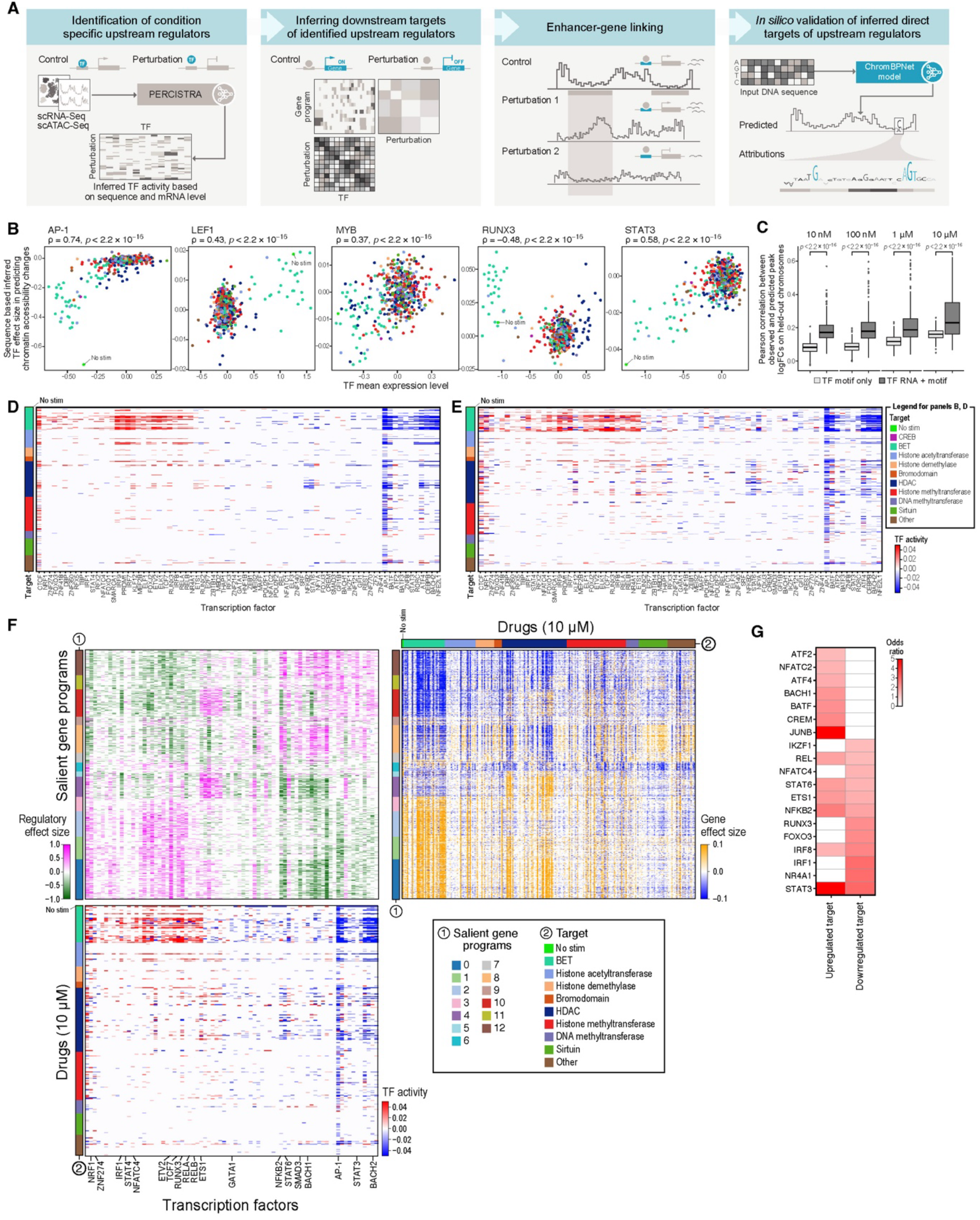
PERCISTRA framework for dissecting gene regulatory networks. (**A)** PERCISTRA overview. (**B**) Significant association between differential TF activity by TF motif accessibility and TF RNA level. Association between TF motif and locus accessibility (vs. vehicle) (y axis) and mean TF mRNA expression (x axis) in each drug-dose combination (dots, color coded in **D**). (**C-E**) Benefits of integrated TF motifs, chromatin accessibility and TF mRNA for TF activity prediction. (C) Distribution of Pearson correlation coefficients (y axis) between predicted and observed peak log-fold-change values on held-out chromosomes (2, 8, and 11) for TF activity models relying only on motif-chromatin accessibility association (light grey) or on jointly modeling TF mRNA expression and its motif accessibility (dark grey) across dose levels (x axis). P-value: one-tailed Wilcoxon matched-pairs signed-rank test. (D,E) Estimated activity scores (color bars) for each TF (columns) in each drug treatments (rows, ordered by drug target, color code) at the maximal (10 μM) dose based on motif-accessibility relation only (D) or by also incorporating TF mRNA expression (E). (**F**) Association of TF activity to gene programs. Regulatory effect size (activator (pink) / repressor (green) color bar, top left matrix; regularized regression; **Methods**) of each TF (top left matrix, columns) on each gene (top left, rows, ordered by MoCAVI gene program, bar 1), based on each TF’s activity (red/blue color bar, bottom matrix) in each 10 μM drug treatment (bottom rows, ordered by drug target (bar 2)) and on the RNA abundance change (top right, orange/blue color bar, **Methods**) of each gene (top right, rows, ordered by gene program (bar 1)) to each 10 μM drug treatment compared to STIM (vehicle) (top right, columns, ordered by drug target (bar 2)). **G)** Correspondence of inferred TF activity to genetic TF KO effects. Transcription factors (rows) whose inferred down- or upregulated targets (columns) are enriched (color bar, odds ratio, Fisher’s Exact Test, FDR<0.1) in targets identified by an independent KO of the same TF in the same system ^6^. White: non-significant.

PERCISTRA consists of four sequential steps. (1) Differential TF activity inference. PERCISTRA infers which TFs have altered activity in response to specific perturbations based on coordinated changes in TF mRNA expression (*trans* factors)^64^ and in chromatin accessibility at their corresponding *cis*-binding elements across the genome. By comparing perturbed and control conditions, PERCISTRA infers TFs that are likely to be differentially active or repressed under specific conditions. (2) TF-target genes inference. PERCISTRA infers the downstream target genes regulated (directly *or* indirectly) by each TF by associating gene expression changes to inferred TF activities (from (1)) across drug-dosage conditions. (3) Enhancer-target gene inference. PERCISTRA associates enhancer regions to their putative target genes by correlating changes in chromatin accessibility of enhancers with changes in RNA levels of nearby genes across drug-dosage conditions. This step helps build locus–gene regulatory maps grounded in interventions, which enhance interpretability and biological relevance. (4) *In silico* validation and refinement of inferred regulatory interactions. PERCISTRA provides orthogonal evidence supporting the TF–target gene connections (from (2)) by fitting ChromBPNet models^65^ to drug-dosage conditions in order to identify (*de novo*) TF footprints in the linked regulatory loci and confirm the presence of expected TF sequence motifs at regulatory regions distally or proximally to direct TF targets.

### TFs with altered activity in drug-dosage conditions

TF activity is often inferred from changes in the TF’s mRNA levels and their correlation to the expression of putative target genes^66–68^, but TF expression can be noisy (fig. S14A), and not reflect TF activity^69^. To better infer causal and directional regulatory signals of global TF activity, we related the mechanistic signal of the relation between a given TF’s motif presence and locus accessibility to TF expression. First, we organized motifs for 401 core human TFs from the HOCOMOCO motif database (57) into 41 motif-based TF groups, each capturing multiple factors with highly similar binding motifs, along with 275 additional TFs analyzed individually (Methods; Table S7). We assessed differential TF activity by quantifying the effect sizes in a regression model using the presence of putative TF binding sites for predicting chromatin accessibility changes (Methods) and evaluating whether these estimated effects correlated with changes in the mRNA levels of the corresponding TFs. Across all tested drug–dosage conditions, there was a strong global correlation between the differential TF activity scores from the multivariate regression models (Methods) of motif presence in predicting differential chromatin accessibility in a locus (501bp) and the corresponding TFs’ RNA expression levels (Fig. 7B, Table S8). In contrast, this relationship disappeared when the analysis was repeated using permuted motif assignments (fig. S14B, Table S9). Motivated by this concordance, we inferred differential TF activity in response to drug treatment by integrating chromatin accessibility profiles in binding motifs with TF mRNA levels (Methods). Incorporating both motif presence and TF expression markedly improved differential chromatin-accessibility predictions on held-out chromosomes (Methods), compared with using motif information alone (Fig. 7C). By analyzing the interaction terms between TF motif presence and TF mRNA expression levels in the model, we identified 77 TFs that had significant dosage-specific regulatory activity (FDR < 0.1, two-sided t-test) in at least 10 drug conditions (Fig. 7D,E fig. S14C-E). Most of the TFs that were inferred as active have well-established roles in activation and differentiation of human CD4^+^ T-cell, including the AP-1 family TFs (e.g., FOS, FOSB, JUN, JUNB), STAT family members (STAT3, STAT6, STAT4), NFKB family members (REL, RELA, RELB) and regulators of the naive cell state (TCF7, BACH2). Notably, RNA expression levels of CTCF, NRF1, and ZNF274 were significantly associated with genome-wide chromatin accessibility changes in several drug treatments, independent of motif presence (fig. S14J). CTCF has a broad role in organizing genome architecture^70–72^ and its expression may capture broader regulatory dynamics beyond its direct binding sites, possibly through its indirect targets. Together, these findings support that our integrative motif-expression framework faithfully reconstructs the core transcriptional network governing human CD4^+^ T-cell activation and its dynamic response to small-molecule perturbations.

### Downstream target identification for differentially active TFs

To link inferred differential TF activities to downstream expression responses across treatments, we modeled the log-fold changes of each gene in MoCAVI’s gene programs as a function of differential TF activities across 1,085 drug–dose conditions, while conditioning on the presence of corresponding TF motifs in regulatory regions of a gene to enforce regulatory directionality (Fig. 7A, step 2, Fig. 7F, Methods). By defining (in step (1)) a TF’s activity based on the ability of its cognate motifs to predict chromatin accessibility changes, rather than relying solely on TF mRNA levels, we aim to reduce spurious TF–target associations arising from broad co-regulation. Consistent with this, significant associations derived from our composite TF activity model showed an 85% signed overlap with those obtained using a model relying on TF motifs (but not TF mRNA)–only model (fig. S15A, top left, Methods), but only 65% signed overlap a model relying on TF mRNA (but not motifs) (fig. S15A, top right, Methods).

The predicted regulatory functions of multiple TFs by this analysis aligned well with established biological functions in T cell activation and effector programs, supporting the model’s inferences (Table S10) with significant enrichment of target genes in certain gene programs (fig. S15B,C and Table S11). For example, RELB, ETS1 and RUNX family TFs positively regulated the cytotoxic effector gene program sGP4, a gene program enriched for cytotoxicity effector genes such as *IFNG, GZMB*, and *IL2RA*, and upregulated in response to HDACi, whereas TCF7 suppressed it, consistent with its known role in repressing effector gene expression in naïve T cells^73^. Conversely, TCF7 activity was positively associated with the naive T cell program sGP1, which was upregulated in response to BETi and non-stimulated cells, and was negatively associated with effector-associated AP-1 family TFs (JUNB) and by BATF.

We corroborated the inferred TF targets with a published dataset generated by profiling CD4^+^ T cells with TF knockouts under the same stimulus^6^. The genes identified as regulated (up or down) by the TFs in our model were significantly enriched (FDR < 0.1, one-sided Fisher’s exact test) among the experimental (genetic) targets for those TFs for 19 of 28 evaluable TFs (those present both in the model and the published dataset; Fig. 7G), supporting our TF target inference approach.

### Mapping the regulatory circuit at the locus level

We next investigated the genomic loci involved in the charted regulatory responses and mapped *genomic loci - binding TF - target gene* relationships based on co-accessibility and expression patterns across drug-doses.

The genomic loci most differentially accessible across all drug-dose conditions were predominantly linked to CD4^+^ T cell activation and had differential accessibility between vehicle-treated (stimulated) and non-stimulated cells (28% of our ~115K peak set) (Fig. 8A-C). These differentially accessible regions fell in distal or intronic regions, while accessibility of promoter regions remained relatively unchanged, suggesting TF-driven enhancer remodeling rather than basal promoter regulation (fig. S16A-C). This is consistent with previous findings that accessibility in distal enhancers accounts for most of chromatin remodeling in regulating stimulation-induced gene expression changes in T cells^74–76^. Conversely, distinct sets of peaks showed strong differential accessibility across different drug families (Fig. 8D,E), with a subset of impacted peaks noticeably proximal to TSS and spanning promoters, introns and exons (compare Fig. 8D,E to fig. S16A,B).

**Fig. 8.**
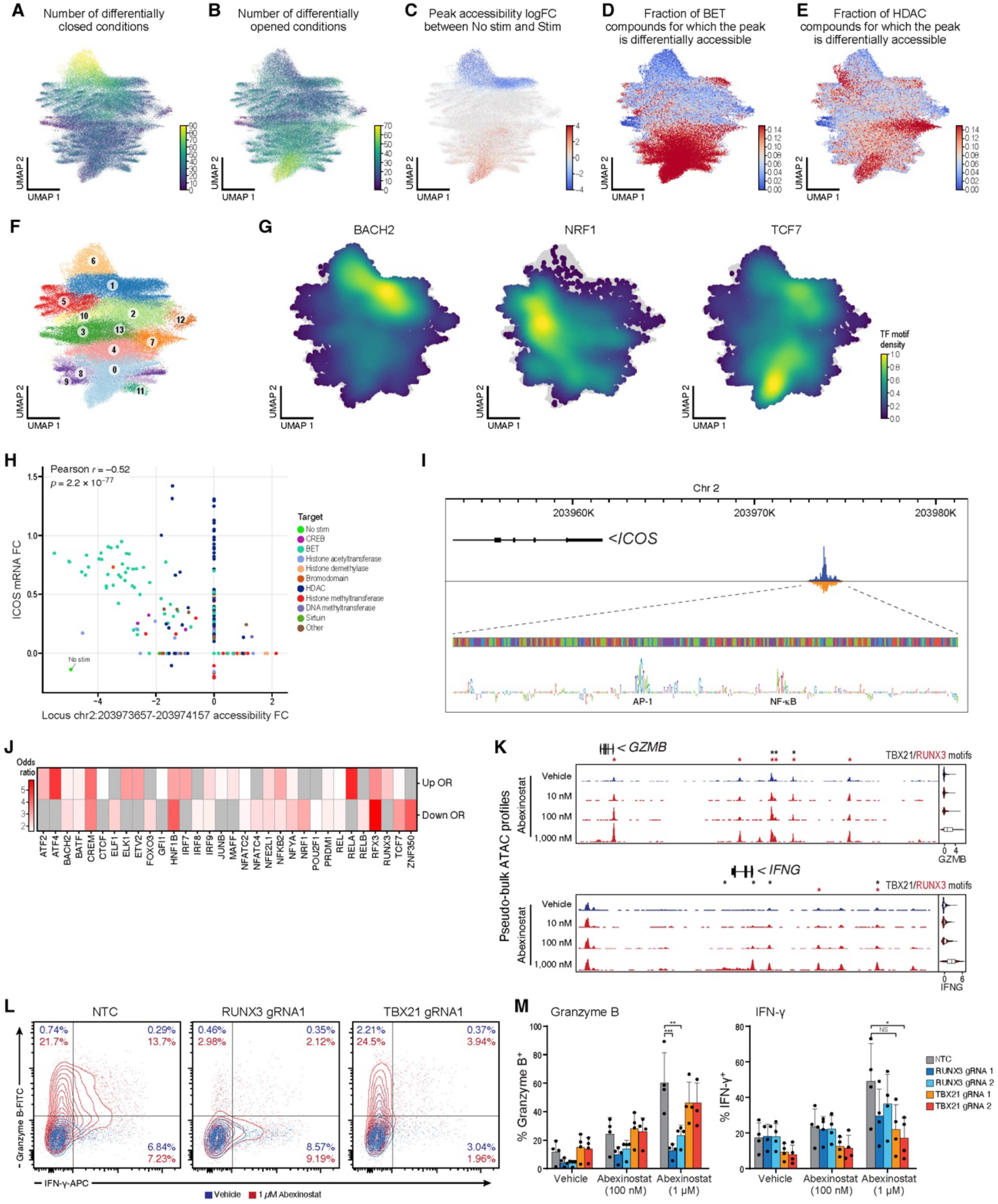
Transcription factor targets inferred from drug-responsive chromatin landscapes. (**A-G**) Measured drug and inferred TF impact on chromatin accessibility. UMAP representation of ATAC-seq peak profiles in each region (dot) across drug-dose treatments relative to vehicle control colored by the number of conditions in which each peak was significantly less (A) or more (B) accessible (vs. vehicle; log-fold change of peak accessibility in unstimulated conditions (vs. stimulated (vehicle)) (C); fraction of BETi (D) or HDACi (E) treatments in which each peak was significantly more accessible (vs. vehicle); peak cluster assignment (F), and density of peaks harboring BACH2 (left), TCF7 (middle) or NRF1 (right) motifs. (**H**,**I**) Enhancer-target gene map at the ICOS locus. (H) Chromatin accessibility fold-change at the distal peak *chr2_203973657:203974157* (x axis) and RNA only estimated abundance change (**Methods**) of its closest gene, ICOS (y axis), in each drug-dose condition (dots) compared to STIM (vehicle) condition. Accessibility fold changes and RNA abundance changes with FDR > 0.1 are set to zero. **(I)** *ICOS* locus (top), pseudobulk ATAC-seq profile for vehicle cells at the adjacent peak *203973657:203974157* (middle), and ChromBPNet attributions at the zoomed-in peak site (bottom). (**J**) ChromBPnet predicted target binding sites are enriched for inferred TF-target gene relationships. Enrichment (odds ratio, Fisher’s Exact Test, FDR < 0.1) of inferred up- or down-regulated targets (rows) of TFs (columns) for corresponding ChromBPnet attributions at nearby peaks. Grey: non-significant values (FDR >= 0.1). (**K**) Chromatin regions differentially accessible between abexinostat-treated and vehicle cells at the *GZMB* and *IFNG* loci. Genomic tracks (rows, left) of pseudo-bulk ATAC reads in each drug-dose combination at the *GZMB* (top) and *IFNG* (bottom) loci. Right: distribution of normalized RNA counts (x axis). * TBX21 (black) and RUNX3 (red) sequence motif matches. (**L**,**M**) Proinflammatory cytokine expression in HDAC inhibitor-treated T cells is dependent on TBX21 and RUNX3. (L) Flow cytometry plots for levels of of Granzyme B (y axis) and IFNγ (x axis) in vehicle (blue) or abexinostat-treated (red) non-targeting control (left), RUNX3 KO (middle) and TBX21 KO (right) CD4^+^ T cells after 72 h of *in vitro* stimulation. (M) Percent (y axis, mean ± s.e.m.) of vehicle or abexinostat-treated cells (x axis) expressing Granzyme B (left) or IFNγ (right) as assessed by flow cytometry in non-targeting control (gray), RUNX3 KO (blue) and TBX21 KO (red) CD4^+^ T cells (*n* = 4 independent donors). \**P* < 5×10^−2^, \*\**P* < 10^−2^, \*\*\**P* < 10^−3^, NS, not significant (One-way ANOVA with Dunnett’s multiple-comparisons test).

To identify those epigenetic inhibitor effects that are independent of T cell activation, we also embedded the treatments using only the chromatin accessibility changes at 67,962 regions that were invariant between stimulated and unstimulated conditions (in the absence of drug). Compounds from the same family often grouped together, especially at higher doses, as seen for BETi, HDACi, and HMTi, consistent with shared mechanisms of action and convergent remodeling of chromatin architecture (fig. S16D,E). The tighter grouping at higher doses may reflect broader inhibition within drug families, where higher concentrations engage additional family members and drive more coordinated chromatin responses.

Next, we grouped all ~115K peaks into 13 clusters (C0-C12) by the co-variation of their accessibility across 1,085 drug-dose conditions (Fig. 8F and fig. S16F). Distinct peak clusters were enriched for different sets of TF motif sites (Fig. 8G and fig. S16G), and the genes most proximal to peaks in each cluster were enriched in particular salient gene programs (Fig. S16H), suggesting that the clusters reflect for groups of co-regulated genomic regions that impact corresponding gene programs. For example, RUNX3 TF activity was associated with the cytotoxic program sGP4 (fig. S15B), and the RUNX3 motif was enriched in peak clusters C1 and C7 (fig. S16G), which, in turn, were enriched for proximal sGP4 genes (fig. S16H). Cluster C1 loci had increased accessibility in vehicle-treated stimulated *vs*. non-stimulated cells, whereas C7 accessibility was independent of T cell activation but impacted by HDACi-treatment (Fig. 8E and fig. S16F). This supports a model where the impact of HDAC inhibition on sGP4 is mediated through the RUNX3 TF acting in accessible sites. In another example, TCF7 activity is associated with the naive T cell gene program sGP1, and both the TCF7 motif and sGP1 genes were significantly enriched in peak cluster C0 (fig. S15B and S16G,H), which was more accessible in BETi-treated and unstimulated cells than in vehicle-treated stimulated cells (Fig. 8D). This supports a model where the impact of BETi in maintaining a naive state is mediated through TCF7 action through its binding site in relevant regulatory regions upstream of naive state genes, which become more accessible by the inhibitor. Finally, the activation-associated effector program sGP10 was linked to peak clusters C1 and C6 (fig. S16H) representing peaks that open upon T cell activation (Fig. 8C,F and fig. S16F) and were enriched for AP-1 family TFs (fig. S16G), consistent with the known role of AP-1 activity in regulating stimulation-induced chromatin opening^76^.

Next, we generated enhancer-target gene maps, based on enhancer regions whose chromatin accessibility changes predict the expression of genes within the salient gene programs (sGPs). The differential accessibility of 4,426 of 18,581 enhancer regions whose nearest gene was in the 13 sGPs was significantly associated (FDR < 0.1, two-sided T-test, Table S11) with the differential expression of their proximal genes across 1,085 drug–dosage conditions (e.g., see Fig. 8H for *ICOS*). To distinguish direct TF targets and further support these TF-target gene inferences, we trained ChromBPNet models^65^ for each drug dosage condition and examined motif attribution scores that are significantly enriched relative to background attributions for each locus linked to these genes at the highest dose with at least 100 cell profiles (e.g., Fig. 8I for the *ICOS* locus, Methods). Notably, for 32 TFs out of 77 tested, these predicted target binding sites were significantly enriched (FDR < 0.1, one-sided Fisher’s Exact Test) among our inferred TF-target gene relationships (Fig. 8J, Methods).

Finally, to illustrate the functional relevance of these inferred regulatory networks, we focused on the ‘naive-like’ and cytotoxic phenotypes induced in BETi- and HDACi-treated cells, respectively (using the motif only (not TF mRNA expression) model, Methods). The strength of TF activity (positive or negative) under BET inhibitor treatments closely aligned with the magnitude of the ‘naive-like’ salient gene program 1 under those treatments (sGP1) (fig. S17A). Across BETi conditions, AP-1 family TF activity was downregulated, whereas CTCF activity was broadly increased, suggesting that AP-1- and CTCF-associated chromatin regulation is particularly susceptible to BETi. The increased activity of naive-associated TF TCF7, as well as ELF1, ELK1 and ETV2 was less susceptible, as it was upregulated mostly in treatments with strong sGP1 induction at highest doses (fig. S17A). These results support a model in which BET inhibition prevents establishment of the T cell effector-associated chromatin landscape primarily through suppression of AP-1-associated regions. At lower doses, this suppression is incomplete, allowing partial engagement of activation-associated TF circuitry. At higher effective doses, activation-dependent chromatin architecture is strongly impaired, and the cells adopt a more ‘naive-like’ transcriptional and epigenetic state.

A similar pattern was observed with HDAC inhibitor treatments, where AP-1, BATF, and NF-κB TF activity was decreased and the activity of RUNX2, RUNX3 and TBX21 was increased in a manner correlated with the strength of expression of the sGP4 program (fig. S17B). Notably, conditions showing concomitant increases in RUNX family and TBX21 activity, such as in high doses of tacedinaline, givinostat, abexinostat, and scriptaid, also exhibited markedly elevated GZMB and IFNG expression (fig. S17B and fig. S18A-C) and a strong correlation between the RNA levels of TBX21, RUNX3, IFNG and GZMB (fig. S18A), further supporting a potential role for these TFs in regulating effector gene expression. Pseudo-bulk ATAC profiles at the *GZMB* and *IFNG* loci had a strong increase of chromatin accessibility and a concomitant increase in the GZMB and IFNG RNA for cells treated with abexinostat (100 nM and 1 µM), aligning with the dependence of HDACi on the enhanced activity of these putative enhancers (Fig. 8K). These impacted enhancers were strongly enriched for TBX21 (*IFNG* and, to a lesser extent, *GZMB*) and RUNX3 (*GZMB*) TF motifs, suggesting that these TFs may help mediate the increase in enhancer accessibility and gene expression at these loci.

To validate the role of RUNX3 and TBX21 in HDACi-dependent cytokine induction, we tested the impact of the HDACi abexinostat in CD4^+^ T cells that were perturbed using targeted CRISPR gene-editing with two independent single guide RNAs (sgRNAs) (Fig. 8L). As expected, abexinostat treatment of WT cells markedly increased GZMB and IFNG expression in a dose-dependent manner, compared to vehicle (Fig. 8M, grey bars). Consistent with our models’ predictions, RUNX3 perturbation strongly reduced GZMB induction by abexinostat and, to a lesser extent, IFNG, whereas TBX21 loss had only a marginal effect on GZMB but markedly attenuated IFNG expression (Fig. 8M, blue and red bars). Together, these findings highlight distinct yet collaborative roles for RUNX3 and TBX21 in regulating HDACi-driven cytotoxic gene programs. More broadly, our results highlight how incorporation of chromatin accessibility and motif annotations enables the mechanistic dissection of gene regulatory networks, allowing us to move beyond correlative expression analyses and identify TFs mediating drug-induced T cell responses.

## Discussion

Here, we present a scalable integrated platform for multimodal chemical screening and extend existing single-cell multi-omics protocols into icCITE-plex and DOGMA-plex to enable prospective multi-omic profiling of chemical perturbations across RNA, protein and chromatin accessibility. Our two-step barcoding approach, pairing defined combinations of viral barcodes with hashing antibody-oligonucleotides, enables multiplexing of thousands of perturbations within a single experiment and provides a scalable framework for dissecting drug-induced molecular phenotypes at single-cell resolution. Notably, because sample indexing occurs at the level of cell hashing rather than during library preparation, all cells can be pooled and stained together for protein measurement, eliminating the need for separate staining of each sample pool. Moreover, because the platform does not rely on fixatives, it preserves the quality of the parental assay and is, in principle, compatible with any 3′-based RNA capture method as well as additional modalities. Going forward, we envision that stable cell lines carrying defined viral barcodes could be generated, allowing repeated use of the same barcoded populations across independent screens and long-term storage for future experiments.

We also introduced MoCAVI, a contrastive analysis approach that separates drug-specific effects from natural cellular heterogeneity with increased sensitivity compared to existing methods. Compared to the state-of-the-art approaches, typically limited to one or two modalities, MoCAVI simultaneously integrates RNA, protein, and chromatin accessibility data, enabling comprehensive MoA studies that capture coordinated effects across molecular layers and can be readily extended to handle additional emerging modalities such as methylation^77^ or polyadenylation^78^. The contrastive analysis framework also presents opportunities for spatiotemporal extensions. For example, recent time-series single-cell CRISPR screens^79^ demonstrated the feasibility of capturing perturbation dynamics, suggesting that temporal integration could reveal how drug effects propagate through regulatory networks over time. Similarly, emerging spatial perturbation platforms like Perturb-Multimodal^80^, PerturbView^81^, Perturb-FISH^82^, CRISPRmap^83^ and Perturb-map^84^, which combine imaging with sequencing in intact tissues, could extend MoCAVI to understand how perturbation effects manifest across tissue architecture and cellular neighborhoods, moving beyond single-cell analysis to systems-level mechanism discovery.

A key strength of MoCAVI’s framework is its ability to decompose variation into complementary background and salient latent spaces that are not required to be mutually exclusive. The background space captures natural variation present in control populations, while the salient space highlights states in which gene expression density deviates from this baseline and requires additional dimensions to explain perturbation effects. As a result, gene programs such as those associated with cell cycle phases can appear in both spaces: they reflect natural variation across cells in the background, yet specific perturbations can also induce abnormal expression patterns along these same biological axes even after controlling for baseline heterogeneity. This dual representation allows MoCAVI to distinguish perturbations that bias cells toward pre-existing states from those that drive cells into novel phenotypic territories.

MoCAVI’s framework facilitated mechanistic grouping of compounds into drug modules based on similar phenotypic effects, agnostic of their intended targets. Moreover, by examining dose-dependent associations of compounds with different groupings, we identified tendencies that were unexpected based on the current known drug MoA. Such phenotypic shifts, especially at higher doses, suggest potential off-target activities, highlighting the importance of leveraging multiple modalities while assessing phenotypic similarity to uncover unanticipated mechanisms and interpretability of chemical perturbation screens, as well as noting that results should be interpreted with care given the relatively high doses for many phenotypic effects in our study. By aligning drug-induced latent representations with CRISPR-mediated genetic knockouts, we matched unexpected phenotypic shifts to specific genetic perturbations, thereby nominating candidate genes responsible for off-target effects. This integrative analysis establishes a systematic strategy to bridge chemical and genetic perturbations, enabling more precise annotation of compound activity profiles. Moving forward, several methodological advances could enhance MoCAVI’s interpretability. Building on recent advances in causal representation learning^85–87^ and incorporating more sophisticated sparse mechanism shift models^88,89^ would provide more principled avenues for modeling chemical perturbations as combinations of genetic knockouts and identifying minimal genetic perturbation sets that recapitulate drug phenotypes.

MoCAVI also clarifies the distinct contribution of each modality. We found that protein data alone was sufficient to recover global relationships between perturbations and resolve major phenotypic states. This suggests that high-throughput drug screening platforms based on multiplexed protein measurements, such as mass cytometry approaches like CyTOF^90^ or sequencing-based single-cell protein methods such as SCITO-seq^91^, may be adequate for capturing coarse phenotypic similarity across treatments. However, because current antibody-based protein panels necessarily interrogate a limited feature space, their ability to resolve finer-grained regulatory programs is inherently constrained. In contrast, transcriptome profiling captures distributed network-level changes, providing the dimensionality required for mechanistic dissection at higher resolution. Thus, while protein-based representations may enable scalable phenotypic screening, comprehensive mechanistic interpretation will benefit from integration with global RNA profiles.

To help decipher molecular mechanisms that underlie drug effects, we introduced PERCISTRA, a framework for inferring regulatory networks explaining the drug-affected pathways by jointly modeling chromatin accessibility and gene expression changes across drug and doses. By integrating *cis*-regulatory accessibility with TF expression within and across ~1,200 drug-dosage conditions spanning 299 epigenetic inhibitors at four doses, PERCISTRA uncovered “TF - bound locus - target gene” links for gene programs that are up- or down-regulated upon treatment with epigenetic inhibitors in activated human CD4^+^ T cells. We showed that combining TF motif accessibility with mRNA levels improves inference of treatment-specific TF activity, revealing many regulators such as AP-1, REL, RUNX3 and TCF7 with roles consistent with known effector and naïve programs. Treatment-dependent covariation further enabled prediction of TF–target gene links, validated by external knockout datasets. At the locus level, enhancer clusters co-responsive to specific drug families mapped coherently to gene programs and TF activity, exemplified by RUNX3- and TBX21-dependent regulation of cytotoxic enhancers under HDACi, which we validated experimentally. Together, these findings highlighted how integrative modeling of drug treatment screens can reveal both drug-specific and drug group-wide regulatory effects, providing insight into the MoA of epigenetic inhibitors in activated human CD4^+^ T cells.

While PERCISTRA offers several advances, it also has limitations that affect our analysis of the epigenetic inhibitors screen. First, causal inference with PERCISTRA relies on the diversity and resolution of treatment conditions, as it utilizes the covariation patterns across conditions. While ~1,200 drug–dosage conditions provide rich variation, the inference of causal relationships is limited to the specific space of treatment tested. Regulators that are not modulated under these conditions may remain undetected. Second, inference of enhancer–target gene relationships, while supported by motif attribution and validation experiments, may still include confounding factors, highlighting the need for systematic perturbation of putative regulatory elements. Additionally, our current approach only focuses on linking enhancer regions to nearby genes, which may miss distal regulatory interactions mediated by three-dimensional chromatin architecture. Third, although covariation across treatment conditions provides stronger evidence for causal relationships than purely observational multi-omics approaches, some inferred TF–gene links may remain indirect due to extensive co-regulation. To mitigate collinearity and confounding, we defined TF activity based on motif-associated chromatin accessibility changes rather than just TF mRNA expression, jointly modeled all TFs for each target gene using regularized multivariate regression, and conditioned TF effects on the presence of cognate motifs in regulatory ATAC-seq peaks. While these constraints reduce spurious associations, ChromBPNet footprinting further refines predictions but does not fully distinguish direct from indirect regulatory effects. Finally, epigenetic inhibitors often have pleiotropic effects and dose-dependent off-target activities. These broad influences complicated interpretation of TF activity changes and enhancer remodeling, especially at higher doses. Thus, we cannot rule out the possibility that broad effects on chromatin and downstream transcription, such as the dichotomy observed in the response in our study, can lead to some spurious associations even when relying on all available modalities. Future approaches that integrate the entire dataset within a unified framework, instead of performing sequential inference steps, could enhance consistency and transitivity among the intermediate and final results.

Overall, we demonstrated the wide applicability of single-cell high-content multimodal drug screening in uncovering the phenotypic effects of SM drug treatments and inferring their underlying mechanism across multiple regulatory dimensions. Our analysis also revealed that different modalities provide complementary information, protein markers capture broad cellular states such as T cell activation, while RNA and chromatin accessibility offer finer granularity, thus highlighting important considerations for experimental design and resource allocation. As per-cell profiling costs continue to decline and throughput scales up, we envision that these and future approaches will enable systematic, high-resolution cataloging of both on- and off-target drug activity and cellular phenotypes across complex and heterogeneous cellular landscapes.

## Acknowledgements

We thank members of the Sakaguchi lab; the Regev lab; and others, particularly

P. Yadollahpour, S. Melo-Carlos, J. Kamm and S. Nair for fruitful discussions on multimodal data integration and analysis. The authors thank Leslie Gaffney and Anna Hupalowska for their help in figure making.

## Data and code availability

Raw and processed count data will be deposited in the National Center for Biotechnology Information’s Gene Expression Omnibus upon publication of the manuscript. The reference implementation of MoCAVI along with tutorials is publicly available on GitHub (https://github.com/Genentech/MoCAVI). PERCISTRA’s reference implementation and reproducibility code is publicly available on GitHub (https://github.com/EraslanBas/PERCISTRA).

## Methods

### Cell culture

Frozen PBMCs from healthy donors were obtained from Cellular Technology Limited or STEMCELL Technologies Inc. and processed immediately after thawing. For separation of naive CD4^+^ T_conv_ cells from PBMCs, CD4^+^ T cells were first enriched by the EasySep™ Human CD4^+^ T Cell Enrichment Kit (STEMCELL Technologies Inc.). CD4^+^ CD25^−^ CD45RA^+^ naive T_conv_ cells were next sorted on a BD FACSAria III or BD FACSAria Fusion system. T cell culture medium was composed of ImmunoCult™-XF T Cell Expansion Medium (STEMCELL Technologies Inc.), 100 U/ml of penicillin and 100 µg/ml of streptomycin. Cells were seeded at a density of 1×10^6^ cells/mL and stimulated for 72 hours using plate-bound 10 µg anti-CD3e (BioLegend; UCHT1) and 5 µg anti-CD28 (BioLegend; CD28.2) in culture medium supplemented with 50 IU/mL IL-2 (Kyowa Pharmaceutical Industry; cat# 4987-058-69793-1).

For mouse cells, splenic CD4^+^CD62L^+^CD44^−^CD25^−^ T_conv_ cells from C57BL/6J mice (CLEA, Japan, Inc.) were sorted on a BD FACSAria Fusion system and seeded at a density of 1×10^6^ cells/mL and stimulated for 72 hours using plate-bound 10 µg anti-CD3e (BD; Clone 145-2C11) and 5 µg anti-CD28 (BD; Clone 37.51) in culture medium consisting of RPMI-1640 (Nacalai Tesque) supplemented with 10% Fetal Bovine Serum (Thermo Fisher Scientific; cat# 10437028) 50 IU/mL IL-2 (Kyowa Pharmaceutical Industry; cat# 4987-058-69793-1), 100 U/ml of penicillin and 100 µg/ml of streptomycin.

### Mice

Mice were housed at the Animal Resource Center for Infectious Diseases of Osaka University with a 12-h light–dark cycle, and mice were kept at a temperature of 21.5–24.5°C with humidity ranging from 30–60%. All procedures were performed in accordance with the National Institutes of Health Guide for the Care and Use of Laboratory Animals^94^ and approved by the Committee on Animal Research of Osaka University.

### Flow cytometry

For flow cytometric analysis, cells were stained with antibodies for cell surface proteins and Live/Dead Dye. In some cases, cells were fixed and permeabilized using the Foxp3/Transcription Factor Staining Buffer Set (Thermo Fisher; 00-5523-00) followed by intracellular staining. For intracellular cytokine staining, stimulated cells (STEMCELL Technologies; #10990) were treated with brefeldin A (Thermo Fisher; 00-4506-51) at 37°C for 3 hours and then processed using the Cytofix/Cytoperm™ Fixation/Permeabilization Kit (BD; 554714). Stained cells were analyzed or sorted on the BD FACSCanto, BD FACSAria Fusion or FACSAria III systems and collected using FACSDiva software (version 9.1, BD Biosciences). A list of antibodies used is in Table S12 under the “FACS antibodies” tab.

### Viral production and transduction

A modified CellTag expression vector was generated by replacing the original CMV promoter in pSMAL-CellTag-V3 (Addgene # 115645) with a EF-1α promoter gene fragment (Twist Bioscience) using Gibson Assembly^®^ (NEB; E2611S). Individual colonies harboring unique barcodes were picked and propagated separately. All barcode sequences were sequenced and validated using Sanger sequencing on an Applied Biosystems 3500xL Genetic Analyzer. A list of viral barcodes used in this study is in Table S1 and Table S5.

For validation experiments with genetic knockouts, an ‘All-in-One’ Cas9-sgRNA expression vector was cloned by replacing the EF1α core promoter from lentiCRISPR v2-dCas9 (Addgene plasmid #112233) with a MSCV promoter fragment (IDT), and the dCas9-P2A-Puromycin cassette was substituted with gene fragments (IDT) encoding Cas9-T2A-mScarlet3 using Gibson Assembly^®^. sgRNA cloning was performed as previously described^95^ with digestion using the restriction enzyme Esp3I (Thermo Scientific; ER0452). A list of sgRNA sequences used is in Table S12 under the ‘sgRNA’ tab.

Lentivirus was generated by co-transfecting Lenti-X 293T cells (Takara; 632180) with EF1α-modified Celltag plasmids, psPAX2 (Addgene #12260), and pCMV-VSV-G (Addgene #8454) using Lipofectamine 3000 (Thermo Fisher; #L3000015), following the manufacturer’s protocol. Viral supernatants were collected at 48 and 72 hours post-transfection, concentrated (ALSTEM; VC100) and stored at −80°C for later use.

For infection of both human and murine activated T cells, titered amounts of lentivirus were used 24 hours post-activation. Cells were spinfected by centrifugation 1,220 g, 90 minutes, 32°C.

### Drug treatment

Transduced cells were rested and expanded in 100 IU/mL IL-2-(Kyowa Pharmaceutical Industry; cat# 4987-058-69793-1) containing culture medium 72 hours post-stimulation and fresh IL-2-containing medium was added every two days. Cells were expanded for a minimum of 14 days before sorting for use. Successfully transduced cells were sorted to purity based on eGFP (CellTag) or mScarlet3 (CRISPR experiments) expression on a BD FACSAria Fusion or FACSAria III system.

Sorted cells were plated in 96-well flat-bottom plates at a density of 1×10^6^ cells/mL in culture medium containing a pre-diluted solution of the appropriate compound or vehicle. The medium was supplemented with 50 IU/mL IL-2 (Kyowa Pharmaceutical Industry; cat# 4987-058-69793-1) and either 5 µL CD3/CD28 Dynabeads (Thermo Fisher Scientific; #11132D) for the species-mix icCITE-plex experiment, or 5 µL ImmunoCult CD3/CD28/CD2 T Cell Activator (STEMCELL Technologies; #10990) for the DOGMA-plex experiment. Cells were then exposed to the compounds at the specified concentration for 72 hours.

72 hours post-activation, activation beads were removed (for the icCITE-plex experiment) and cells were washed twice with FACS buffer (2% FBS in PBS) before pooling by plate. Cells for each plate were stained with defined combinations of two specific TotalSeq-A hashtag antibodies (BioLegend; Table S1 and Table S5) as per the manufacturer’s recommendations. Hashed cells were washed and live cells were enriched and pooled by cell sorting on a BD FACSAria Fusion or FACSAria III system. Sorted cells were blocked with TruStain FcX (4223020, BioLegend) and then stained with a TotalSeq-A barcoded surface antibody panel (BioLegend; Table S1 and Table S5) for 30 min on ice. Stained cells were washed three times before being processed as per the icCITE-seq protocol described below.

### icCITE-plex library preparation

icCITE-plex libraries were prepared essentially as previously described 6,96. Briefly, approximately 1 million stained cells were resuspended with Cell Staining Buffer (420201, BioLegend) and pelleted by centrifugation for 5 min at 500g. The supernatant was removed, leaving approximately 50 µl residual volume. The cell pellet was thoroughly resuspended in the residual liquid and then fixed and permeabilized by dropwise addition of 1 ml pre-chilled True-Phos Perm Buffer (425401, BioLegend) while vortexing. Fixed cells were incubated overnight at −20°C.

On the next day, fixed cells were thawed on ice and then centrifuged at 4°C at 2,000g for 5 min. Pelleted cells were washed with 2 ml ice-cold Intracellular Wash Buffer (1×) with 2 mM dithiothreitol and 0.2 U µl−1 Protector RNase Inhibitor (3335402001, Roche). Cells were centrifuged again to completely remove the supernatant. Intracellular staining was then performed using Intracellular Wash Buffer (1×; custom part no. 900002577, BioLegend), with the addition of 2 mM dithiothreitol, TruStain FcX (4223020, BioLegend), True Stain Monocyte Blocker (426102, BioLegend) and 1 U µl−1 Protector RNase Inhibitor (3335402001, Roche) in a 100 µl volume, as per the manufacturer’s recommendations. Stained cells were washed three times with 1 ml of Intracellular Wash Buffer (1×), and then resuspended in Intracellular Wash Buffer (1x) with 2 mM dithiothreitol and 0.2 U µl−1 RNase inhibitor. A maximum of 4 µl of the cell suspension was used for processing with the Chromium Next GEM Single Cell 3′ Kit v3.1 (1000268, 10x Genomics) and 3′ Feature Barcode Kit (1000262, 10X Genomics) according to the manufacturer’s recommendations. For two of the libraries, the RT Enzyme C (10x Genomics; PN 2000102) from Step A 1.1a was replaced with 8.7 µl of Maxima H Minus Reverse Transcriptase (Thermo Scientific; EP0742). A list of intracellular antibodies used in this study is in Table S1 under the tab ‘Intracellular staining’.

Celltag libraries were enriched by a two-step dial-out PCR protocol, essentially as previously described 97 with modified PCR primers (Table S12 under the ‘PCR primers’ tab). TotalSeq-A hashtag and surface antibody libraries were prepared as per the manufacturer’s recommendations (BioLegend). TotalSeq-B intracellular antibody libraries were prepared as per the guidelines in the Chromium Single Cell 3′ Reagent Kits User Guide (v3.1 Chemistry) with Feature Barcoding Technology for Cell Surface Protein (CG000206, 10x Genomics, User Guide). Resulting libraries were sequenced on an Illumina NovaSeq6000.

### DOGMA-plex library preparation

DOGMA-plex libraries were prepared essentially as previously described 7. Briefly, stained cells were pelleted by centrifugation for 5 min at 4°C and 500g. Cells were permeabilized with an isotonic lysis buffer (20 mM Tris-HCl pH 7.4, 150 mM NaCl, 3 mM MgCl2, 0.01% Digitonin and 2 U µl−1 of Protector RNase Inhibitor (3335402001, Roche)) for 5 min on ice. Next, 1 mL of chilled wash buffer (20 mM Tris-HCl pH 7.4, 150 mM NaCl, 3 mM MgCl2 and 1 U µl−1 of Protector RNase Inhibitor (3335402001, Roche)) was added and cells were mixed by inversion before centrifugation for 5 min at 4°C and 500g. The supernatant was removed and cells were resuspended in wash buffer and counted. Cells were transposed and processed with the 10x Genomics Epi Multiome ATAC + Gene Expression kit (CG000338 RevD, 10x Genomics) as per the manufacturer’s recommendations with modifications as previously described^7^. Celltag viral barcode libraries were enriched as described above, using the purified cDNA fraction isolated from Step 6.2n as input. Note that it is essential to use only 10 ng of cDNA as input and the minimal number of PCR cycles necessary to reduce PCR artifacts.

### Preprocessing for icCITE-plex/DOGMA-plex data

For gene expression for the icCITE-plex dataset, sequencing reads were separately aligned to the hg38 and mm10 reference genomes using Cell Ranger (v7.0.0). For chromatin accessibility and gene expression data from the DOGMA-plex dataset, sequence reads were aligned to the hg38 reference genome using Cell Ranger - ARC (v2.0.0). Surface protein tag abundances, hashtag antibody barcode abundances and Celltag viral barcode library reads were estimated using the kallisto (v0.44.0), bustools (v0.39.3) and kite (v0.02) frameworks^98,99^. Intracellular protein tag abundances were estimated using Cell Ranger (v7.0.0).

### icCITE-seq data processing

#### Initial quality check

Cells that had less than one UMI barcode in any of the available modalities (Celltag, HTO, ADT, TSB or gene expression) were initially filtered out. Cells with high mitochondrial UMI counts (> 8%), low RNA UMI counts (<1.5k), low protein counts (<10) or abnormally high RNA UMI counts (>100k) were then removed.

#### Species assignment

To assign cells to species, the fractions of UMIs assigned to either human or mouse genes were assessed. Cells were assigned as “human”/ “mouse” if the fraction of UMIs assigned to human/mouse genes was more than 92%. All remaining cells were deemed ambiguous and labeled as “doublets”. Except when mentioned otherwise, the rest of the analysis was performed separately for each species.

#### Perturbation assignment

To assign cells to plates, HTO counts were demultiplexed with demuxEM^100^, using a minimum UMI HTO count of 10. A similar approach was used to assign cells to wells, using the cell tag barcodes, with a minimum UMI CT count of 2.

#### Genomic features filtering

Genes with less than 500 UMIs were filtered out and only the top 5,000 variable genes were kept using scanpy^101^. Surface proteins and intracellular proteins that had low counts (< 10 UMIs) or high collinearity (> 0.5 Pearson correlation followed by manual inspection) with the HTO counts were filtered out, leaving a total of 83 surface proteins and 41 intracellular proteins. Transcripts from mitochondrial genes and from ribosomal protein genes were filtered out to reduce potential confounding effects in the downstream analysis.

### The MoCAVI generative model

For cell *n*, let *t*_*n*_ ∈ {0,1…, *K*} be its assigned environment label as a categorical variable, where *K* denotes the number of distinct perturbations with non-trivial effect, and 0 denotes the baseline environment (i.e., cells assigned to control perturbations or to perturbations with no effect). Let *x*_*n*_ denote the *G*-dimensional vector of RNA counts (*G* genes), *y*_*n*_ the *P*-dimensional vector of protein measurements (*P* proteins), and *a*_*n*_ the *R*-dimensional vector of fragment counts for chromatin accessibility (*R* regions). *b*_*n*_ is also considered, a *B*-dimensional one-hot vector describing the batch index in case the data presents batch effects.

MoCAVI estimates a background generative distribution *p*_*θ*_(*x*_*n*_, *y*_*n*_, *a*_*n*_ ∣ *t*_*n*_ = 0, *b*_*n*_) for cells from the baseline environment (*t*_*n*_ = 0), as well as a target generative distribution *p*_*θ*_(*x*_*n*_, *y*_*n*_, *a*_*n*_ ∣ *t*_*n*_, *b*_*n*_) for cells from a non-baseline environment (*t*_*n*_ ≠ 0). *θ* is used to refer to all the parameters of the generative model (details about the inference procedure appear below). Below, the target generative distribution is specified (i.e., for now, *t*_*n*_ ≠ 0).

#### Priors

The p-dimensional background cell representation

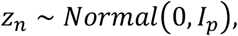

encodes the heterogeneity of phenotypes in cells from the baseline environment (control cells). The q-dimensional salient cell representation

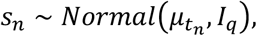

encodes the effect of the drugs. Parameters *µ* ∈ *R*^*K*×*q*^ represent how each perturbation may shift the prior mean for the salient variables *p*_*θ*_(*s*_*n*_ | *t*_*n*_) for every environment *t*_*n*_. The specification of environment-specific priors avoids shrinking the effect of distinct perturbations towards a common distribution, as can happen with ContrastiveVI^27^ and thus significantly improves the quality of the cellular representations (Fig. 2E,F, fig. S3). It is noted that similar modulation of the prior as a function of the perturbation label appears in other recent works about identifiable deep generative models^36,85,102^.

#### Gene expression likelihood

Following^24^, for every gene *g* ∈ {1,…, *G*}, the RNA count *x*_*ng*_ for gene *g* in cell *n* follows the negative binomial distribution

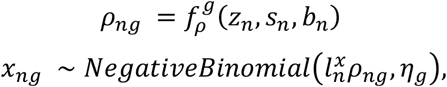

where 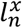 denotes the RNA size factor, and *η* _*g*_ denotes the gene-specific inverse dispersion parameter. *ρ*_*n*_ is the output of a neural network that takes as input *z*_*n*_, *s*_*n*_ and *b*_*n*_, and applies a softmax non-linearity to the output to ensure that the output can be interpreted as a gene expression frequency.

The RNA size factor is estimated from data as 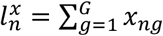, while *η*_*g*_ is treated as a parameter to be inferred via maximum likelihood.

#### Protein abundance likelihood

Following^25^, every protein *p* ∈ {1,…, *P*}, the protein abundance *y*_*np*_ for protein *p* in cell *n* follows a mixture of negative binomial distributions, defined as follows:

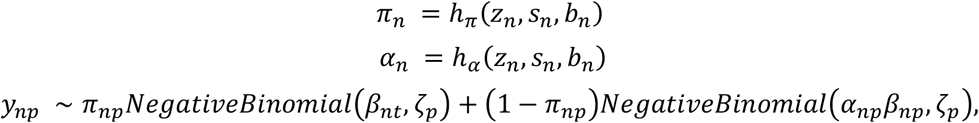

where the mixture weight *π*_*n*_ and the fold-change of the foreground protein abundance *α*_*n*_ are outputs of neural networks *h*_*π*_ and *h*_*α*_, that both takes as input *z*_*n*_, *s*_*n*_ and *b*_*n*_. *ζ*_*t*_ is the inverse-dispersion parameter of the negative binomial distributions. The background protein abundance *β*_*n*_ is learned via (amortized) maximum likelihood. Neural network *h*_*π*_ has a sigmoid nonlinearity for its output layer, *h*_*β*_ and has an exponential nonlinearity for its output layer, and *h*_*α*_ has a ReLU non-linearity with an offset of 1 for its output layer. The offset ensures identifiability of the two components of this mixture (i.e., *α*_*np*_*β*_*np*_ must be the mean of the protein abundance after denoising).

#### Chromatin accessibility likelihood

Following^35,103^, for every region *r* ∈ {1,…, *R*}, the fragment count for chromatin accessibility *a*_*rn*_ for region *r* in cell *n* follows a Poisson distribution, defined as follows:

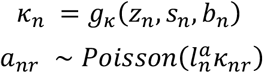

where 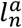 denotes the accessibility size factor, and *κ*_*n*_ is the output of a neural network that takes as put *z*_*n*_, *s*_*n*_ and *b*_*n*_, and applies a softmax non-linearity to the output to ensure that the output can be interpreted as an accessibility frequency. The chromatin size factor is estimated from data as 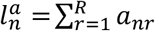.

#### Background generative distribution

Following^27^, the generative process described above applies to any cell *n* from a non-baseline environment *t*_*n*_ ≠ 0. In the case of a cell from a baseline environment *t*_*n*_ = 0, the value of the salient variables *s*_*n*_ is set to zero:

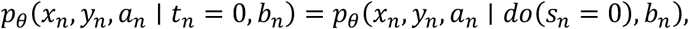

where the *do*(⋅) operator denotes a deterministic intervention.

### Inference for MoCAVI

Neither the model evidence for the baseline environment, *p*_*θ*_(*x*_*n*_, *y*_*n*_, *a*_*n*_ ∣ *t*_*n*_ = ∅, *b*_*n*_) or for the perturbed cells *p*_*θ*_(*x*_*n*_, *y*_*n*_, *a*_*n*_ ∣ *t*_*n*_, *b*_*n*_) can be exactly computed because of the intractable integral over the latent variables. Therefore, variational inference is used to approximate the posterior distributions, as well as learn the model parameters^104^.

#### Variational distribution

For non-baseline environments, the variational distribution admits the following parameterization:

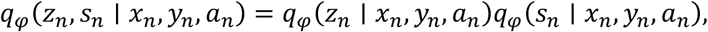

where each distribution is modeled with a product of expert distribution^105^ to incorporate information from each modality, for example:

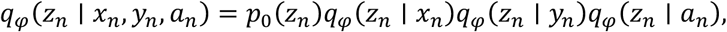

where *p*_0_ is the isotropic Gaussian distribution, and each of the variational distributions *q*_*φ*_(*z*_*n*_ ∣⋅) is Gaussian with diagonal covariance. The product of experts of Gaussian distributions is Gaussian, with mean and covariance matrices available in closed form. The parameters for each of those three distributions are the output of neural networks (encoder network) taking a single modality as input. Similarly, for *s*_*n*_:

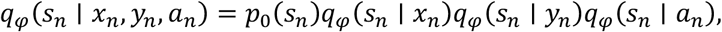

As well as for the baseline environment (*t*_*n*_ = 0), but solely relying on the distribution *q*_*φ*_(*z*_*n*_ ∣ *x*_*n*_, *y*_*n*_, *a*_*n*_), since *s*_*n*_ is deterministically set to zero.

#### Numerical optimization

The composite evidence lower bound (ELBO) is defined as follows:

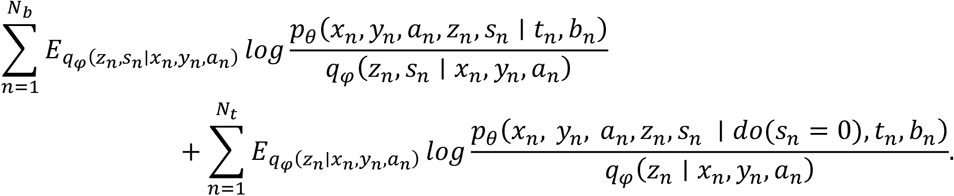

It is optimized with respect to the variational parameters and model parameters using stochastic gradients^106^, evaluated by sampling 128 observations from each dataset, as well as samples from the variational distribution. Model parameters are optimized with the Adam method^107^ with weight decay. Standard multi-layer perceptron architecture is used, with common activation functions (for example, exponential, softmax, sigmoid), to encode the variational and generative distributions.

#### Salient encoder regularization

As highlighted in previous implementations of deep generative models for contrastive analysis^27,32,108,109^, the latent spaces *z*_*n*_ and *s*_*n*_ may contain redundant information when the number of latent variables *p* or *q* becomes large. To avoid this behavior, the encoder networks parameterizing the approximate posterior over the salient latent space are regularized, following Multi-GroupVI^109^ and ContrastiveVI^27^. More precisely, let us denote the approximate posterior over the salient latent space as

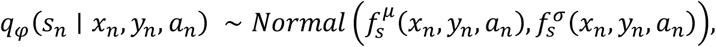

where 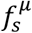 and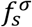 designate respectively the mean and variance of the approximate posterior as a function of the input data. The composite ELBO is altered to include, in addition, the following penalization:

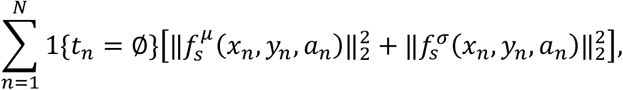

which corresponds to the squared Wasserstein-2 distance between the approximate posterior and the Dirac at zero^110^ (i.e., what the value of *s*_*n*_ should be for the background cells).

#### Environments definition

To apply MoCAVI, each cell *n* must be assigned to an environment *t*_*n*_. Out of all the possible environments, a notable one is the baseline environment (*t*_*n*_ = ∅). For the data presented in this study, the baseline environment was defined as the union of the control cells, as well as all the cells from perturbations assessed to have a non-significant effect. Then, every remaining perturbation was assigned to its own environment.

Any available method can be used to assess whether a perturbation has a significant effect (e.g., the energy distance^111^, or differential expression^112^). In this study, an extension of the TotalVI algorithm was applied, in which the prior *p*(*z*_*n*_) is changed to depend on the perturbation label *t*_*n*_. Then, changes were quantified in latent spaces between each perturbation and the control cells using the energy distance ^113^, and an empirical null distribution was created based on sampling subsets of control cells. Any perturbation that has an empirical p-value of more than 0.05 is considered non-significant.

### Post-hoc interpretation of the MoCAVI model

#### Modality weights

An importance score was attributed to each modality, and each latent space as follows, proceeding separately for each space. For example, the importance of each of the modalities *x*_*n*_, *y*_*n*_ and *a*_*n*_ for latent variable *z*_*n*_ was calculated. For that, let us note as 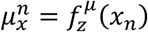 and 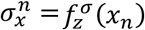 the output of amortization network for the gene expression expert:

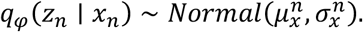

Similarly, 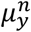, 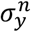 is introduced for the protein expert, as well as 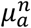, 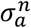 for the chromatin accessibility expert. The importance 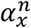 of gene expression for the background latent space is defined as:

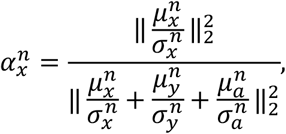

where the expression for 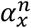 aims at assessing the contribution of the modality *x* compared to other modalities, in terms of the Euclidean norm. The importance of the other modalities, in either the background or the salient latent space was defined similarly.

#### Background molecular map

Interpretation of the background latent space inferred from MoCAVI may be difficult when the heterogeneity in control cells is not composed of discrete states, but rather well-described by continuous variation, with potentially several axes of variation. This is often the case for perturbation screens within a single cell type, and especially while working with immune cells, for which scRNA-seq data often exhibit continuous heterogeneity^40^.

To interpret the background latent variables, Hotspot analysis was performed^41^ on the control cells, as embedded by *z*_*n*_. The Hotspot method identifies clusters of genes that co-vary in the latent space, thus easily enabling a full characterization of the heterogeneity of continuous phenotypic variation. Using only the control cells for this task is important, as it ensures that Hotspot does not use any data related to the perturbation to construct this molecular map.

After clustering of a gene-gene auto-correlation matrix from Hotspot, Metascape was used to annotate the resulting gene programs^114^. Additionally, the score of each of those background gene programs was calculated for every cell in the data set (including perturbed cells), after using a z-score normalization, taking as reference the score in the control cells. This helped quantify how those gene programs were changed upon perturbation.

#### Salient molecular map

Although the salient embeddings are interesting in their own rights and directly interpretable, a post-hoc interpretative procedure was specifically introduced to identify genes (as well as proteins and genomic regions) that are most associated with perturbations, as opposed to those varying in the background.

To this end, the contribution of a gene *g* to a perturbation *t*^*^ was evaluated by calculating a salient-specific log-fold change (ssLFC). Given a cell *n* with measurements *X*_*n*_ = (*x*_*n*_, *y*_*n*_, *a*_*n*_), perturbation *t*_*n*_ = *t*^*^, and a gene *g*, the ssLFC is calculated as:

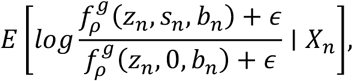

where *ϵ* designates a small offset used for stability, as in^115^. This amounts to comparing the expression of cell *n* to the imputed value of a pseudo-control cell with the same background latent variable, but projected onto the space of control cells. The expectations in the numerator and denominator are computed using the plug-in estimator^116^. Briefly, this means that the (intractable) posterior is replaced by the approximate posterior obtained by the variational inference procedure. The ssLFC may be calculated for each cell, and averaged across all cells *n* such that *t*_*n*_ = *t*^*^.

To assess for significance, the methodology from lvm-DE was followed^115^. First, let us define the following probabilistic event:

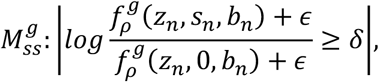

where *δ* is the desired level of effect size. Then, let us calculate the posterior probability of this event 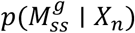 using the plug-in estimator (i.e., replacing the posterior by the approximate posterior).

This calculation departs from the traditional paradigm of differential expression in several ways (fig. S2D, Supplementary Note 1). First, this procedure aims at controlling for the background variation, and should be used if it is expected to yield more interpretable outputs. Second, it compares perturbed cells to their in-silico imputation of what the gene expression would have been had the cell not been perturbed (paired comparison). By comparison, differential expression would compare expression of the perturbed cells to the control cells, in an unpaired fashion. A suite of results assessing the robustness of this novel procedure appears in Supplementary Note 2.

Protein abundance changes are assessed similarly, by replacing *f*_*ρ*_ by *h*_*β*_ ⋅ (1 − *h*_*α*_), as are chromatin accessibility changes, by replacing *f*_*ρ*_ by *g*_*κ*_.

#### Differential abundance

Following the methodology from ContrastiveVI^27^and scVI^24^, the implementation of differential expression was extended to also assess changes in each of the modalities, for example, for comparing the gene expression profiles of two non-baseline environments. For gene *g*, and pair of cells *α, β*, the probability of differential abundance (here, expression) must be computed. First, let us define differential abundance as a probabilistic event:

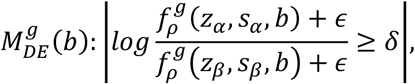

where *ϵ* designates a small offset used for stability, as in lvm-DE ^115^, and *δ* is the desired level of effect size. Let us then marginalize the latent variables, as well as categorical covariates to calculate the posterior probability of differential expression:

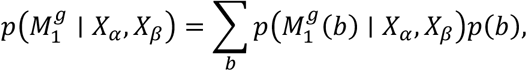

where each individual posterior probability 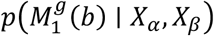 is computed using the plug-in estimator (i.e., replacing the posterior by the approximate posterior).

In order to compare two cell populations *A* = (*α*_1_,…, *α*_*A*_) and *B* = (*β*_1_,…, *β*_*B*_), the posterior probabilities of differential abundance for random pairs of cells in each population are averaged. If one group of cells is part of a baseline environment (*t* = 0), then *s* is replaced by 0 in all of the above calculations.

The calculation of the posterior probabilities allows for either the calculation of a Bayes factor or of a binary label that indicates the genes that are differentially abundant with a controlled posterior expected false discovery rate at a desired level.

Such calculations can be conducted similarity with protein abundance, replacing *f*_*ρ*_ by *h*_*β*_ ⋅ (1 − *h*_*α*_), as well as chromatin accessibility, replacing *f*_*ρ*_ by *g*_*κ*_.

### Ablation studies and comparison between MoCAVI and ContrastiveVI

MoCAVI is most related to the ContrastiveVI model^27^, but differs from it in several ways. First, MoCAVI is extended to handle arbitrary subsets of genomic modalities. Second, MoCAVI’s constitution and treatment of the background and target data sets is distinct, as motivated by early investigations of ContrastiveVI on the icCITE-seq data set. Finally, MoCAVI specifies perturbation-specific prior distributions for the salient variables *p*_*θ*_(*s*_*n*_ ∣ *t*_*n*_). The merits of these modifications, and the impact on downstream performance were evaluated as part of an ablation study.

#### Overfitting the background data set

When ContrastiveVI was applied to the icCITE-seq data set, it stopped training after a few epochs, showing signs of overfitting, and yielding poor latent representations of the perturbations. Because scVI was not overfitting on the same data, and that the number of parameters of the ContrastiveVI model is on the same order of magnitude as scVI, it was hypothesized that the issue would come from splitting the data set into two. Indeed, in cases where the number of cells in the background data set is too small, the part of the model that is specific to the baseline environment could overfit. Several ideas were explored to remedy this problem, as described below.

#### Weighing baseline and non-baseline environments

In those models, each of the target and background dataset are treated differently. A composite objective function taking into account both data sets is therefore implemented. Previous implementations of deep generative models for contrastive analysis use one data loader for each of the data sets *X*^*b*^ and *X*^*r*^. This is equivalent to maximizing the following objective function:

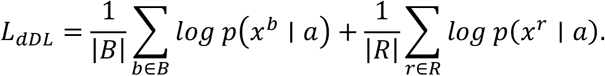

Although this is a sensible approach, its performance heavily depended in practice on the composition (specifically, the number of cells) of the data sets *X*^*b*^ and *X*^*r*^. Traditionally, *X*^*b*^ denotes measurements from the control cells, and *X*^*r*^ from the perturbed cells. In this case, when the number of cells is relatively balanced across data sets, for example, in pooled genetic screens with low MOI, this approach performs well, as presented in the ContrastiveVI paper. However, our experience is that when there are relatively few control cells, as is the case in a chemical screen (because of experimental design), the model may perform poorly because it may overfit the background data set in the case of few observations.

To combat this, a simple idea is to reweigh the contributions of each data set in the loss function, so that the equivalent objective function becomes:

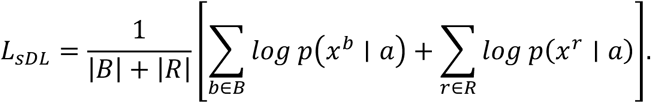

This loss function is referred to as sDL, because it is equivalent to using a single data-loader and sample random data points from each set of observations *X*^*b*^ and *X*^*r*^. This is a standard approach used for training deep generative models across several data sets (e.g., semi-supervised deep generative models). One possible caveat is that when the number of background observations is too large, the model may focus on modelling the control cells, and have lower performance at characterizing the effect of perturbation.

#### Sample rebalancing

As an orthogonal area of interest is the impact of the composition of the target data set on the method’s performance. Intuitively, if many perturbations have no effect, but are modeled with both background and salient latent variables, it biases the inference of salient variables to also encode variations in the control cells, negatively affecting the interpretability of the salient variables. This has already been documented in the first implementation of contrastive analysis approaches with deep generative models (Appendix G of^108^), where the performance of the model severely drops after background data points have been leaked into the target data set.

To investigate this in the context of perturbation screens, a simple method was used to identify perturbations with “non-discernable” effect, and split our data set into three data sets: (1) the set of control cells *X*^*c*^, (2) the set of perturbed but non-affected cells *X*^*neff*^ (here on a perturbation basis, but with different pre-processing this could be conducted at the cell level), and (3) the set of perturbed but likely affected cells *X*^*eff*^. Details on how to separate the perturbed cells into *X*^*neff*^ and *X*^*eff*^ using a conditional VAE appears in the above paragraph entitled “Environments Definition”. Evidently, practitioners are able to use their favorite methods for this purpose.

Then, this framework was applied in three different ways:

- The “vanilla” method, in which *X*^*b*^ = *X*^*c*^ and *X*^*r*^ = *X*^*neff*^ ∪ *X*^*eff*^;
- The “rebalanced” method, in which *X*^*b*^ = *X*^*c*^ ∪ *X*^*neff*^ and *X*^*r*^ = *X*^*eff*^;
- The “removal” method, in which *X*^*b*^ = *X*^*c*^ and *X*^*r*^ = *X*^*eff*^;

The removal method straightforwardly removes cells from any perturbation that has a less than discernible effect size, while the rebalanced method places those cells in the background data set. The optimal choice of the method might depend on whether recycling cells is needed. For example, if there are a large source of control cells, it might be safer to use the “removal” strategy. However, if there are only a few control cells, because of the potential overfitting issues mentioned above, the “rebalanced” version is expected to perform best.

#### Ablation studies

All possible combinations of those model enhancements were benchmarked on the icCITE-seq data set (human and mouse). There are in total 12 variants of ContrastiveVI, with 2 possible re-weighting schemes (dDL or sDL), 3 possible rebalancing schemes (vanilla, rebalanced or removal), and 2 possible priors (perturbation-specific, or global). The two best-performing variants both utilized rebalanced data and perturbation-specific priors, with the top variant (MoCAVI) using sDL reweighting and the second-best using dDL reweighting (fig. S3).

### Latent space benchmarking

MoCAVI’s performance was evaluated on real data sets according to the methodology outlined in the ContrastiveVI study^27^. Specifically, the grouping of perturbations according to their targeted pathway was assessed in the salient latent space. MoCAVI was compared to a broad set of methods, as well as other variants of the ContrastiveVI framework, and on two data sets.

#### Evaluation Metrics

To compare latent spaces across methods, available labels from the data sets that encode the biological annotation of the perturbation were used. In the case of the drug screening data, the MoA, as annotated by the reseller, is used. Importantly, those labels are not seen during training by any method. Data set specific details about perturbation labels appear in a later section.

Given a latent space and perturbation labels, two strategies were used to assess how informative the latent space is: (1) perturbed cells were clustered in the latent space using *k*-means clustering (where k is the number of distinct MoA in the groundtruth annotation), and clustering metrics were evaluated with respect to the labels using the average silhouette width, the adjusted Rand index, and the normalized mutual information; and (2) a logistic regression was trained to predict perturbation labels from embeddings of 75% of the cells, and the accuracy, as well as the F1 score, were evaluated. Details about each metric appear below.

#### Average silhouette width (AWS)

This metric assumes the availability of a Euclidean space where each data point *n* is associated with an embedding vector *t*_*n*_ ∈ *R*^*d*^ where *n* is a data point. Additionally, the AWS requires the definition of cluster assignments *y*_*n*_ ∈ {1,…, *K*}, where *K* is the total number of clusters. For each data point *n*, the silhouette score *SS*_*n*_ of sample *n* is defined as

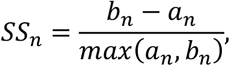

where *a*_*n*_ is the average distance between data point *n* and all other points with the same cluster label, and *b*_*n*_ is the average distance between *n* and all the points in the next nearest cluster. Then, for a data set with *N* samples, the AWS is defined as:

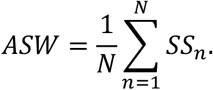

The value of ASW lies between −1 and 1, where a higher value indicates a better ability to distinguish the clusters in the embedding space.

#### Adjusted Rand index (ARI)

The adjusted Rand index (ARI) measures the similarity between two data clusterings, the true assignments *y*_*n*_ ∈ {1,…, *K*} and the predicted assignments *y*′_*n*_ ∈ {1,…, *K*′}, where *n* is a data point, *K* is the total number of true clusters, and *K*′ is the total number of predicted clusters. The ARI takes into account the number of pairings of data points that are in the same or different clusters for both the true and predicted assignments. Specifically, the ARI is defined as:

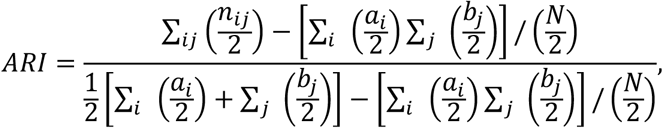

where *n*_*ij*_ is the number of data points that are in both cluster *i* of the true assignments and cluster *j* of the predicted assignments; *a*_*i*_ and *b*_*j*_ are the total number of data points in cluster *i* of the true assignments and cluster *j* of the predicted assignments, respectively; and *N* is the total number of data points. The value of ARI lies between −1 and 1, where a higher value indicates better clustering performance.

#### Normalized mutual information (NMI)

The normalized mutual information (NMI) gauges the similarity between two data partitions (here: cluster assignments): the true assignments *y*_*n*_ ∈ {1,…, *K*} and the predicted assignments *y*′_*n*_ ∈ {1,…, *K*′}, where *n* is a data point, *K* is the total number of true clusters, and *K*′ is the number of predicted clusters. The NMI is computed using the mutual information (MI) between the two assignments and their respective entropies:

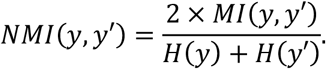

Mutual information between the true and predicted assignments is given by:

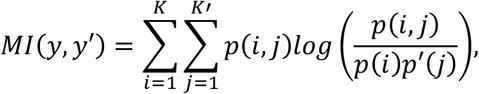

where *p*(*i, j*) is the joint probability of a data point belonging to cluster *i* in the true assignments and cluster *j* in the predicted assignments; and *p*(*i*) and *p*′(*j*) are the probabilities of a data point belonging to cluster *i* and *j* in the true and predicted assignments, respectively.

The entropies of the true and predicted assignments are:

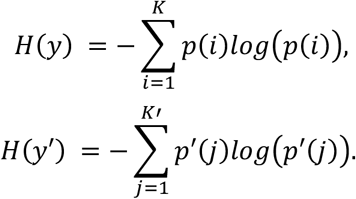

The value of NMI lies in the range [0, 1], where 1 indicates that the two sets of cluster assignments are identical, and 0 indicates no shared information between them.

#### F1 score

The F1 score evaluates the balance between precision and recall in classification tasks. Precision is the proportion of true positive predictions in all positive predictions, while recall (or sensitivity) is the proportion of true positive predictions in all actual positives. The F1 score is the harmonic mean of precision and recall, providing a single measure that balances both, and is defined as:

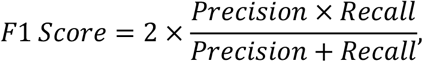

where Precision is 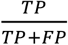, and Recall is 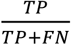. Here, *TP* is the number of true positives, *FP* is the number of false positives, and *FN* is the number of false negatives. The F1 score ranges from 0 to 1, where a higher value indicates a better balance of precision and recall, hence better model performance.

#### Accuracy

Accuracy evaluates the overall effectiveness of a classification model, by measuring the proportion of correct predictions (both true positives and true negatives) in relation to the total number of cases evaluated. It is defined as:

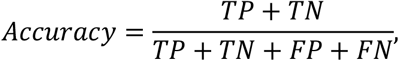

where *TP* and *TN* are the number of true positives and true negatives, respectively, and *FP* and *FN* are the number of false positives and false negatives, respectively. Accuracy ranges from 0 to 1, with higher values indicating a more accurate model.

##### Competing Methods

The following competing methods were considered in our benchmark. Our method was compared to the following methods (1) ContrastiveVI^27^; (2) TotalContrastiveVI^27^ (ContrastiveVI extension for CITE-seq data); (3) ContrastivePCA applied to library size-normalized, log-transformed count data (to compare with a non-DGM baseline)^30^; (4) mixscape (proposed for delineating the effect of perturbations from the control variation)^26^]; (5) scVI (a deep generative model for scRNA-seq data)^24^; (6) totalVI (a deep generative model for CITE-seq count data)^25^; and (7) a typical scRNA-seq analysis workflow in which PCA is applied to library size-normalized, log-transformed counts^101^. All methods were run using default parameters. However, because all those methods learn latent variables, the number of latent variables was set to be the same across all methods.

### MoCAVI application to icCITE-plex data

#### Environment specification

To retain treatments with significant effects, the identifiable version of TotalVI (iTotalVI) described in the section “*Environments Definition”* was applied (each species was processed separately), using the combination of drug and dose as label, the library and the plate as batch, and the selected surface proteins. The baseline environment was defined as the union of all labels such that the energy distance to the control cells was not significantly different from an empirical null of control cells (permutation test).

#### MoCAVI application

MoCAVI was run on gene expression and surface protein data (each species processed separately). The combination of library and plate identifiers was set as the batch variable. The model was trained with 15 salient variables, and 10 background variables.

#### Dosage-sensitivity quantification

To quantify how sensitive a given treatment is to changes in dosage, the shifts induced by the dose in MoCAVI’s salient latent space were used. The effect of each drug was assessed under three non-zero dosages (100 nM, 1 μM, 10 μM) in addition to the control condition (vehicle = 0 nM). The average shift between pairs of doses was assessed, using the Euclidean distance in latent space, if there were enough (> 30) cells for a given dosage, and the mean across all available pairs of dosages was taken.

#### Modality-specific regulatory map

Salient-specific LFCs for each modality were assessed with MoCAVI for all perturbations. Downstream analysis and visualization focused on the set of 49 perturbations (drug-molecule combinations) with highest energy distance with vehicle cells using MoCAVI’s salient latent space. Then, the 49 perturbations were clustered into drug modules with Leiden clustering of the salient latent space (resolution = 2), using the Euclidean metric, and features were clustered into gene programs with Leiden clustering of the change matrix (each observation is a perturbation), using the cosine similarity (resolution = 1.2).

### Integration of icCITE-plex with genetic knock-outs

#### MoCAVI details

A joint model of icCITE-plex RNA profiles and Perturb-seq RNA profiles was learned. All of the unperturbed cells from Perturb-seq were assigned to the baseline environment, and all of the non-stimulated cells (no TCR activation) were assigned to the environment they occupied in the icCITE-plex dataset. These two control conditions served as anchors across the two datasets. MoCAVI was then applied using the same architecture and hyperparameters as for the icCITE-plex dataset, adding the Perturb-seq data as a new batch identifier.

#### Regression-based differential expression

A linear regression model of the normalized, log-transformed expression of every gene was fitted as a function of the perturbation, batch identifier, and total number of UMIs. Significance was assessed with a t-test for the regression coefficients, using the statsmodels python package. FDR was calculated using the Benjamini-Hochberg procedure.

### Cross-species comparison

#### MoCAVI details

A joint model of the human and the mouse RNA profiles was learned. Mouse gene names were mapped to human orthologs using the Mouse Genome Informatics homology database^117^. Mouse and human cells were assigned to different batch identifiers. Control conditions were assigned to the same labels independently of species, but all other drug conditions were kept apart, to avoid overcorrection.

#### Conservation assessment

To assess conservation of drug effects across mouse and human, two similarity metrics were computed at the treatment level and averaged: (1) Cosine similarity between the mean salient embedding of each treatment; and (2) cosine similarity between the MoCAVI-computed log-fold changes of gene expression profiles, using the vehicle condition as a baseline.

#### Statistical significance for dosage matching

To support the statement that treatments in mouse cells consistently better matched to a treatment with a lower dosage in human cells, the conservation assessment metric was used to compute a statistic of how many mouse treatments, across all drugs and dosages, better matched to a human treatment with a lower dosage. Then, the rows of our embedding matrices used to compute similarities were permuted 5,000 times and a significant p-value (p < 0.005) was obtained.

### DOGMA-plex data processing

#### scRNA-seq processing pipeline

An initial library of 550,521 cells was reduced to 498,749 cells after quality control filtering based on mitochondrial read fraction (≤ 0.3), minimum expressed gene count (≥ 500), and minimum total UMI count (≥ 1,000). A total of 20,877 genes expressed in at least 1,000 cells were retained for downstream analyses. Hash and lentiviral barcodes were demultiplexed using the demuxEM algorithm^118^, resulting in 319,841 singlet cells with valid hash–barcode assignments. Counts were normalized to 1e4 UMIs per cell and log-transformed with a pseudocount of 1. A small population of TRGC2+ cells was identified and removed, leaving 314,329 cells for subsequent analyses.

#### Gene expression effect-size estimation

As part of the PERCISTRA pipeline, we assess gene expression fold changes across perturbations using linear regression on normalized expression levels computed separately for every gene. For each gene *g*, we fit the model:

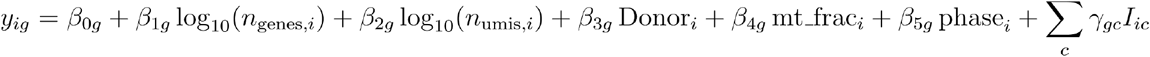

where *y*_*ig*_ is the log1p-normalized expression of gene *g* in cell *i*, and *I*_*ic*_ indicates the drug–dose condition *c*. Donor and cell cycle phase are treated as categorical covariates, while number of detected genes, number of total UMIs, and mitochondrial read percentage are treated as continuous covariates.

The reference level corresponds to the pooled set of stimulated vehicle (STIM) cells across all four dosages (10 nM–10 µM). Thus, *γ*_*gc*_ represents the covariate-adjusted expression change for condition *c* relative to this baseline. Models were fit independently for each gene using ordinary least squares, and coefficients and associated *P*-values were used as effect sizes and significance measures.

### ATAC-seq processing pipeline

#### Peak set generation

ATAC-seq data were processed with ArchR^119^. For each drug+dose combination, a minimum of five pseudo-bulk replicates was constructed, each composed of at least 80 cell profiles, to ensure sufficient coverage and reproducibility. A unified peak set was generated using the MACS2-based peak calling implementation within ArchR on the coverage tracks derived from all drug+dose combinations, with the following parameters: q-value threshold of 0.01, reproducibility filter requiring peaks to be detected in at least five distinct drug+dose combinations, and iterative overlap merging procedure to generate the union peak set. The resulting peak set consisted of 113,547 501bp fixed-width peaks.

#### Identification of differentially accessible peaks across drug-dosage conditions

Differentially accessible peaks were identified for each drug-dosage combinations *vs*. the stimulated control (vehicle) condition by a two-sided Wilcoxon test (ArchR implementation via presto^120^) while correcting for the TSS enrichment score and the number of unique nuclear fragments (log_10_(number of fragments)), as defined in ArchR^119^. Peaks with an absolute log_2_ fold-change > 0.5 and P-value < 0.05 were considered differentially accessible.

### MoCAVI application to the DOGMA-plex data set

#### Environment specification

To retain perturbations with significant effects, the identifiable version of TotalVI (iTotalVI) described in the paragraph *Environments Definition* was applied using the combination of drug and dose as label, the library and the plate as batch, and the selected surface proteins. The baseline environment was defined as the union of all labels such that the energy distance to the control cells was not significantly different from an empirical null of control cells (permutation test).

#### MoCAVI details

MoCAVI was applied to multimodal data (RNA expression, surface protein expression and chromatin accessibility). The combination of library and donor identifiers was set as the batch variable. The model was trained with 25 salient variables and 10 background variables.

#### Modality-specific regulatory map

Total LFCs for each modality were assessed with MoCAVI for all perturbations. Downstream analysis and visualization focused on a set of 220 strong-effect perturbations (drug-molecule combinations). Those perturbations were outliers on the distribution of energy distances of perturbed cells with vehicle cells using MoCAVI’s salient latent space, for each perturbation (fig S12B). Perturbations were clustered into drug modules with Leiden clustering of the salient latent space (resolution = 2), using the Euclidean metric, and features were clustered into gene programs (resolution = 1.2), peak programs (resolution = 1.2), and protein programs (resolution = 1.5) with Leiden clustering of the change matrix (each observation is a perturbation), using the cosine similarity.

#### Modality-specific weight analysis

To quantify the effect of treatments on each modality (here, chromatin accessibility, gene expression), the Euclidean norm of the total LFCs was computed for each treatment in each modality (peaks or expressed genes). To quantify whether a treatment would have been detected or not in one of the modalities, a reference was computed of the same scores for treatments that were not assessed as strong-effect perturbations in our initial processing (outliers in terms of perturbation effects as in fig S12B).

#### Assessment of dose-dependent chromatin-to-transcription effects

For each drug, differentially accessible chromatin peaks were identified at the lowest dose and their nearest genes were determined using pre-computed gene-peak associations from ArchR. Next the proportion of these peaks whose nearest genes became differentially expressed only at the higher dose than the dose of change in their chromatin accessibility (i.e. genes not differentially expressed at low dose but differentially expressed at higher dose) were determined. This analysis was performed separately for BET inhibitors and histone methyltransferase (HMT) inhibitors as a negative control. Statistical significance between the two drug classes was assessed using a two-sample Kolmogorov-Smirnov test (fig. S13C).

### TF motif sets identification

The *motifmatchr* package in the ArchR^119^ pipeline^121^ was used to annotate the 113,547 peaks with binding motif information for 401 transcription factors (TFs) from the HOCOMOCO database (57), yielding a sparse binary matrix of dimensions 113,547 × 401, where each entry indicates the presence or absence of a given TF motif in a specific peak. Next, the similarity between each pair of TF (motifs) was computed as the pairwise Jaccard similarity between their motif occurrence profiles. Specifically, for each pair of TF (motifs) *i* and *j*, the Jaccard index was J(i, j) = |P_i_ ∩ P_j_| / |P_i_ ∪ P_j_| where P_i_ and P_j_ denote the sets of peaks containing motifs for TF *i* and TF *j*, respectively. Next, a fully connected weighted graph was constructed where each TF (motif) is a node and Jaccard indices are edge weights, and TF motif clusters were defined by the *igraph* R package, based on the leading non-negative eigenvector of the graph’s modularity matrix^122^. This yielded 41 TF (motifs) clusters spanning 126 TF (motifs) and another 275 individual TF (motifs) as singletons, to a total of 316 TF sets (incl. singletons) used in downstream analyses.

### Identification of differential TF activity in each drug+dose combination solely based on the sequence and ATAC profiles

Let X ∈ {0,1}^*P ×M*^ be the binary motif matrix, where *X*_*pm*_= 1 if TF *m* has a motif in peak *p*, and 0 otherwise for *P* = 113,547 peaks and *M* = 316 TFs and TF (motif) sets (referenced as TFs from here on) and Y ∈ ℝ^*P ×C*^ is the differential chromatin accessibility matrix, where *Y*_*pc*_ is the log fold-change of peak *p* in drug+dose condition *c*. To estimate differential TF activity vs. stimulated vehicle control at each drug-dose combination *c*, feature selection is first performed via LASSO (least absolute shrinkage and selection operator) regression:

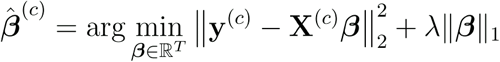

Where the regularization parameter λ is selected by 10 fold-cross validation for each drug+dose condition independently. The significance and effect size of the differential activity of each TF for condition *c* 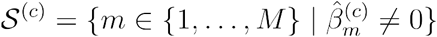, is inferred by ordinary least squares (OLS) models 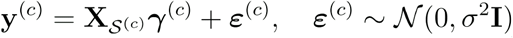, where 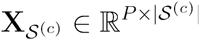 is the full motif matrix, and the coefficient vector 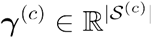 captures the motif-only TF activity scores of the TFs. Specifically, for each TF or TF set *t ∈ 𝒮*^*(c)*^, the scalar 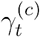 represents the estimated average change in chromatin accessibility at peaks containing its cognate motif in condition *c*, relative to peaks without the motif. For fully conditioned effect sizes and p-values the OLS models are fit on the full TF matrix, and the set of 77 TFs selected during the initial LASSO-based feature selection step are interpreted in the dowstream. Separate models are fit for each drug+dose combination with the statsmodels Python package^123^.

### Identification of differential TF activity in each drug+dose combination based on sequence, chromatin accessibility and TF mRNA levels

To quantify differential TF activity in response to drug treatments, pseudobulk peak-level differential accessibility was modeled using a linear regression framework that incorporates TF motif presence, pseudobulk TF gene expression levels, and their interaction. This modeling was performed separately for each drug *d* ∈{1, …274}, across multiple doses for drugs that had at least 3 doses.

Let *P* = 113,547 denote the number of peaks, *T* = 126 the number of individual TFs for which we had measured scRNA data, and *G* = 41 the number of TF groups, and, for each drug *d, D*_*d*_ ∈{3,4} is the number of doses assayed, leading to a total of N_*d*_ = *P* ×*D*_*d*_ peak-condition observations for drug *d*.

The response variable 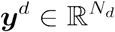 consists of differential chromatin accessibility values (log fold-change) for each peak and each dosage of drug *d*, concatenated by row. For each peak, a feature vector is constructed from the following components:

- TF motif presence: A binary matrix ‘s motif in peak M ∈ {0,1}^*P* ×*T*^, where *M*_*pt*_ =1 indicates the presence of TF
- TF motif set presence: A binary matrix M^group^ ∈ {0,1}^*P* ×*G*^, encoding motif availability for TF sets.
- TF expression levels: For each drug 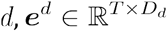 is the pseudobulk mean mRNA expression of the 126 TFs across the *D*_*d*_ drug-dose conditions.

To construct the design matrix for drug *d*, the feature matrices were expanded across dosages as follows:

- TF motif presence matrix, 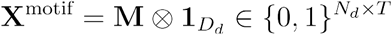 where ⊗ denotes the Kronecker product and 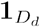 is a length-*D*_*d*_ vector of ones.
- TF expression matrix, 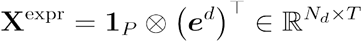
- Motif-expression interaction terms, 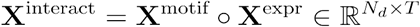 where ◯ denotes element-wise (Hadamard) product.
- TFmotif set matrix, 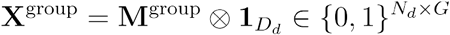

These components were concatenated to form the full design matrix:

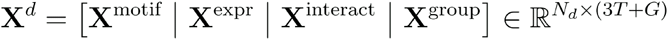

Finally, a standard linear regression model was fit of the form:

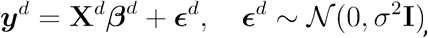

where β^*d*^ ∈ ℝ^3*T*+*G*^ are the regression coefficients for drug *d*, quantifying the contribution of TF motif presence, TF mRNA expression, their interaction, and TF motif sets, to the differential chromatin accessibility observed across all peaks and doses. The model was fit independently for each drug using ordinary least squares, and the resulting coefficients were used to assess the predictive value of individual TFs or TF set under each perturbation.

In this linear model framework, each TF is associated with three distinct regression coefficients: one for motif presence, one for TF expression, and one for their interaction. Specifically, for each TF *t* ∈ {1, …,*T*} and drug *d*, the model estimates:

- 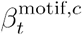: the effect of motif presence on chromatin accessibility in drug+dose combination *c*. Since motif presence is binary and fixed across dosage conditions, this coefficient is drug-specific in estimation and reflects a static feature of the peaks.
- 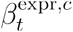: the effect of TF expression in drug+dose combination *c*, independent of motif presence.
- 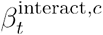: the interaction between motif presence and TF expression, capturing drug+dose-specific regulation.

To quantify the regulatory effect of TF *t* in a specific drug–dose combination *c*, its TF activity score is calculated, defined as its composite accessibility contribution at motif-containing peaks, as follows:

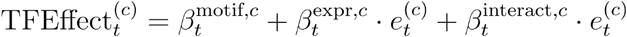

where 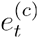 is the mean mRNA expression level of TF *t* in drug+dose combination *c*. This expression reflects how strongly TF *t*’s expression modulates accessibility at its binding sites, accounting for both global and motif-dependent effects.

### Gated TF–target gene regression models

First, all ATAC-seq peaks assigned to each target gene within the MoCAVI gene programs were identified based on nearest-gene annotations. Next, for each gene, a binary motif availability vector was defined indicating whether each TF or TF set had at least one binding site in the peaks assigned to the gene, which then served as a gene-specific gating variable. A LASSO-regularized linear regression model was fit for each gene across 1,085 drug–dose combinations, where the response variable was the gene’s RNA log-fold change (vs. stimulated (vehicle) control) and the predictors were TF activity scores gated by motif availability, such that TFs without an available motif in the regulatory regions of the gene had zero contribution, and only TFs with potential physical binding could influence the gene’s expression, as follows:

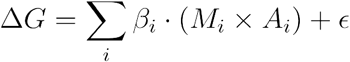

where ΔG is the gene expression log-fold change, A_i_ is the differential activity of TF i, M_i_ is the binary motif availability indicator for that TF at the gene, and *β*_i_ is the learned regulatory effect.

LASSO regularization was applied to control for multicollinearity among TFs and promote sparse, interpretable regulatory relationships, with the regularization parameter (λ) selected independently for each gene using 10-fold cross-validation, optimizing for minimum mean squared prediction error on held-out data. Model performance was quantified using the cross-validated coefficient of determination (R^2^). TF–target regulatory effects were summarized by the non-zero regression coefficients of the final model. Positive coefficients were interpreted as activating relationships, while negative coefficients were interpreted as repressive relationships.

