## Supplementary Information for "Massively multiplex multimodal chemical screens at single-cell resolution"

1. Laboratory of Experimental Immunology, Immunology Frontier Research Center, Osaka University, Osaka, Japan
2. Genentech Research and Early Development, Genentech, South San Francisco, CA, USA
3. Department of Genetics, Stanford University, Stanford, CA, USA
4. Department of Computer Science, Stanford University, Stanford, CA, USA
5. Department of Biology, Stanford University, Stanford, CA, USA
6. Department of Experimental Pathology, Institute for Life and Medical Sciences, Kyoto University, Kyoto, Japan
7. These authors contributed equally to this work.

†. Present Address: Arc Institute, CA, USA

### Supplementary Figures

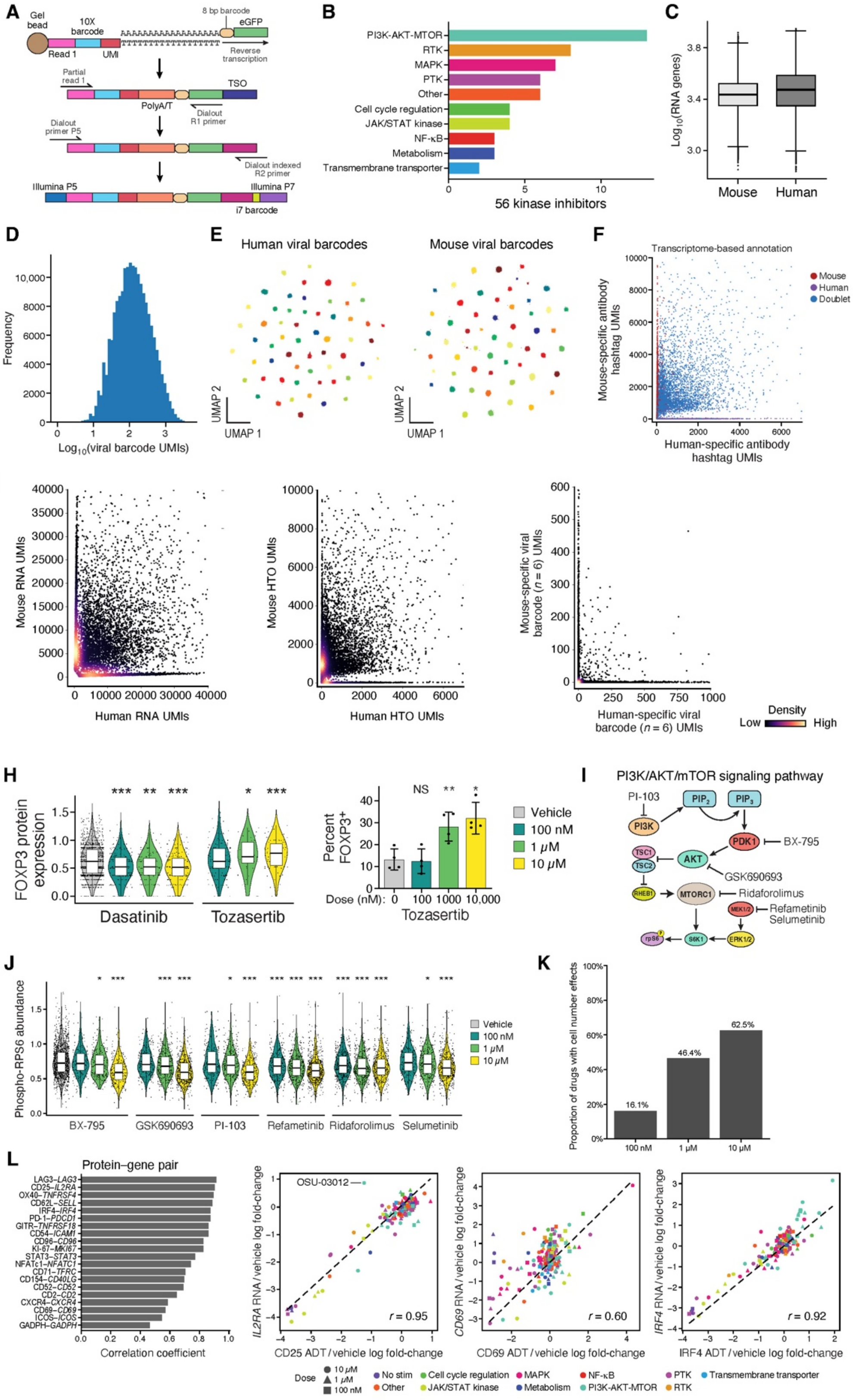

**Fig. S1. Characterization icCITE-plex and small-molecule perturbations, related to Fig. 1.**
(**A,B**) icCITE-plex species-mix experiment. (A) Workflow for capturing mRNA-encoded viral barcodes. (B) Pathway distribution of the 56 kinase inhibitors in the icCITE-plex experiment. (**C-G**) icCITE-plex QC metrics. Distributions of number of genes detected per cell (C, y axis) for mouse and human cells (x axis), and number of viral barcode UMIs recovered per cell (D, x axis). E, UMAP embedding based on icCITE-plex viral barcode expression (dots) in human (left) and mouse (right) cells, colored by viral barcode assignment. (F) Number of human (x axis) and mouse (y axis) antibody hashtag UMIs associated with each cell barcode (dots), colored by transcriptome assignment as human (purple), mouse (red), or both (doublet) (purple). (G) Number of human (x axis) and mouse (y axis) RNA UMIs (left), antibody hashtag UMIs (middle) or species-specific viral barcode UMIs ($n$ = 6 for each species, right) associated with each cell barcode (dot), colored by point density (color bar). (**H**-**J**) Select regulatory effects of SM treatment on protein expression. (H) Left: Distribution of FOXP3 protein levels (y axis) across cells following dasatinib or tozasertib treatment (x axis) (left) (*$P < 5\times10^{-2}$, **$P < 10^{-2}$, ***$P < 10^{-3}$, two-tailed Mann-Whitney U-test). Right: Percent (y axis, mean ± s.e.m across $n$ = 4 independent donors) of vehicle or tozasertib-treated (x axis) $CD4^+$ T cells expressing FOXP3 protein as assessed by flow cytometry *$P < 5\times10^{-2}$, **$P < 10^{-2}$, NS, not significant (repeated-measures one-way ANOVA with Dunnett's multiple-comparisons test). (I) Schematic of PI3K/AKT/mTOR signaling and downstream pathways. Small-molecule inhibitors targeting specific pathway components are annotated. Drawing is adapted from refs. [92,93]. (J) Distribution of phospho-RPS6 protein levels (y axis) across cells under different PI3K/AKT/mTOR inhibitor treatments (x axis), colored by dose (color bar). *$P < 5\times10^{-2}$, ***$P < 10^{-3}$ (two-tailed Mann-Whitney U-test). (**K**) Prevalence of cell number effects across SM treatments. Proportion of drugs (y axis) with significant (**Methods**) cell number effects per dose (x axis). (**L**) Protein-RNA correspondence. Left: Pearson's correlation coefficients (x axis) between the levels of protein and cognate RNA for selected protein-RNA pairs (y axis). Right: Log fold-changes in protein (x axis) and RNA (y axis) expression relative to vehicle for CD25/*IL2RA* (left), CD69/*CD69* (middle) and IRF4/*IRF4* (right) in each drug-dose combination (dots), labeled by drug target (color) and dose (shape).

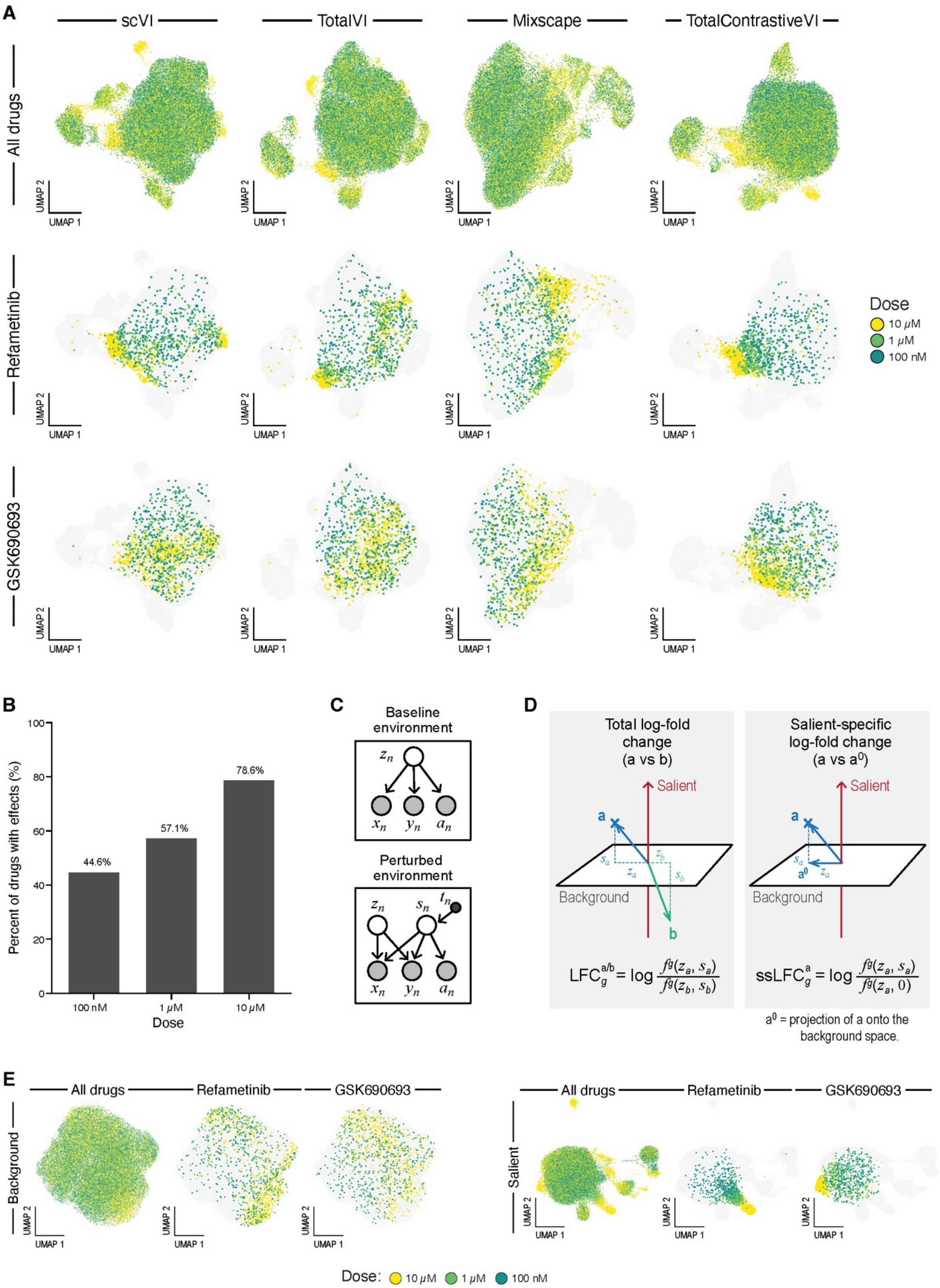

A
scVI
TotalVI
Mixscape
TotalContrastiveVI
All drugs
Refametinib
GSK690693
UMAP 1
UMAP 2
Dose
10 μM
1 μM
100 nM
B
Percent of drugs with effects (%)
44.6%
57.1%
78.6%
100 nM
1 μM
10 μM
Dose
C
Baseline environment
Perturbed environment
$z_n$ $x_n$ $y_n$ $a_n$
$z_n$ $s_n$ $t_n$ $x_n$ $y_n$ $a_n$
D
Total log-fold change (a vs b)
Salient-specific log-fold change (a vs $a^0$)
Salient
Background
$LFC_g^{a/b} = \log \frac{f^g(z_a, s_a)}{f^g(z_b, s_b)}$
$ssLFC_g^{a} = \log \frac{f^g(z_a, s_a)}{f^g(z_a, 0)}$
$a^0$ = projection of a onto the background space.
E
All drugs
Refametinib
GSK690693
Background
Salient
Dose: 10 μM 1 μM 100 nM

**Fig. S2. Additional description of MoCAVI's approach to single-cell perturbation data modeling, related to Fig. 2.**
(**A**) Sensitivity of existing single-cell perturbation analysis methods to dosage. UMAP representation of cell profiles (dots) in the latent space obtained via different embedding techniques (labels on top), colored by dosage across all drugs (top), refametinib (middle) or GSK690693 (bottom). (**B**) Drugs with significant effects. Proportion of drugs (y axis) with significant effect on icCITE-plex profiles at each dose (x axis). (**C**) MoCAVI's graphical model. Observed (shaded vertices) and latent (white vertices) random variables for the baseline (top) and any perturbed (bottom) environment (bottom). Edges: conditional dependence. (**D**) Definition of total and salient-specific log-fold changes. Left: Total log-fold changes are calculated across two cell populations by averaging the log-fold changes estimated from MoCAVI's decoder outputs over many pairs of cells. Right: Salient-specific log-fold changes are estimated for one population of perturbed cells by averaging the log-fold change between the decoded gene expression and a counterfactual gene expression level that represent what the gene expression of that same cell would have been if it were not perturbed. (**E**) Sensitivity of MoCAVI's latent representation to dosage. UMAP representations of cell profiles (dot) from MoCAVI's background (left) and salient (right) space, colored by dose for all drugs, refametinib or GSK690693.

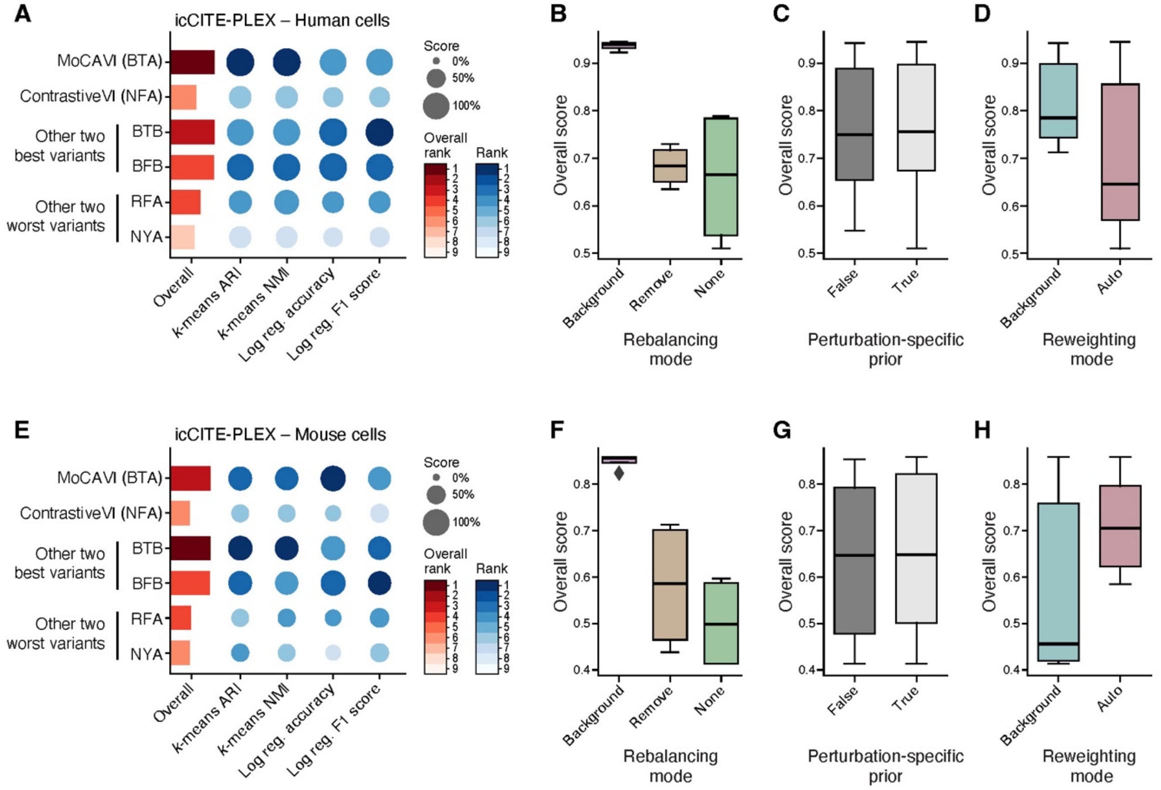


**Fig. S3. MoCAVI ablation studies on icCITE-plex, RNA-only model**, **related to Fig. 2.**
Ablation studies assessing three important MoCAVI parameters (**Methods**). The rebalancing mode handles cell profiles where the treatment had no discernible effect by either placing them as control cells in the baseline environment (Background; B), discarding them (Remove, R), or leaving them in their own non-baseline environment (None, N). The environment-specific prior parameter encodes whether the model has a perturbation-specific prior on salient latent variables (True, [T]) or not (False, [F]). Third, the reweighting mode specifies whether the loss for the background cells is weighted by the proportion of background cells of the entire dataset (Background, B) or not (Auto, A). Variant names are assembled in the order the modifications are introduced. For example, MoCAVI corresponds to the variant BTA. (**A,E).** Score (circle size or bar length) and rank (color intensity) of MoCAVI, ContrastiveVI and two other best and worst performing variants (y axis) by individual metrics (x axis, blue) and that quantify the separation of cells in latent space by drugs' mechanism of action and an overall score averaging all metrics (bars, red) in models learned on the human (A) or mouse (H) data. Higher values: better performance. (**B-H**) Distribution of the overall evaluation score (y axis) of all variants (x axis) grouped by parameter choice for the rebalancing mode (B,F), perturbation-specific prior (C,G) or reweighting mode (D, H). In all Box plots, Centers: median overall score values, upper (lower) box bounds: third (first) quartile; upper (lower) whiskers: third quartile + 1.5 × interquartile range (first quartile − 1.5 × interquartile range).

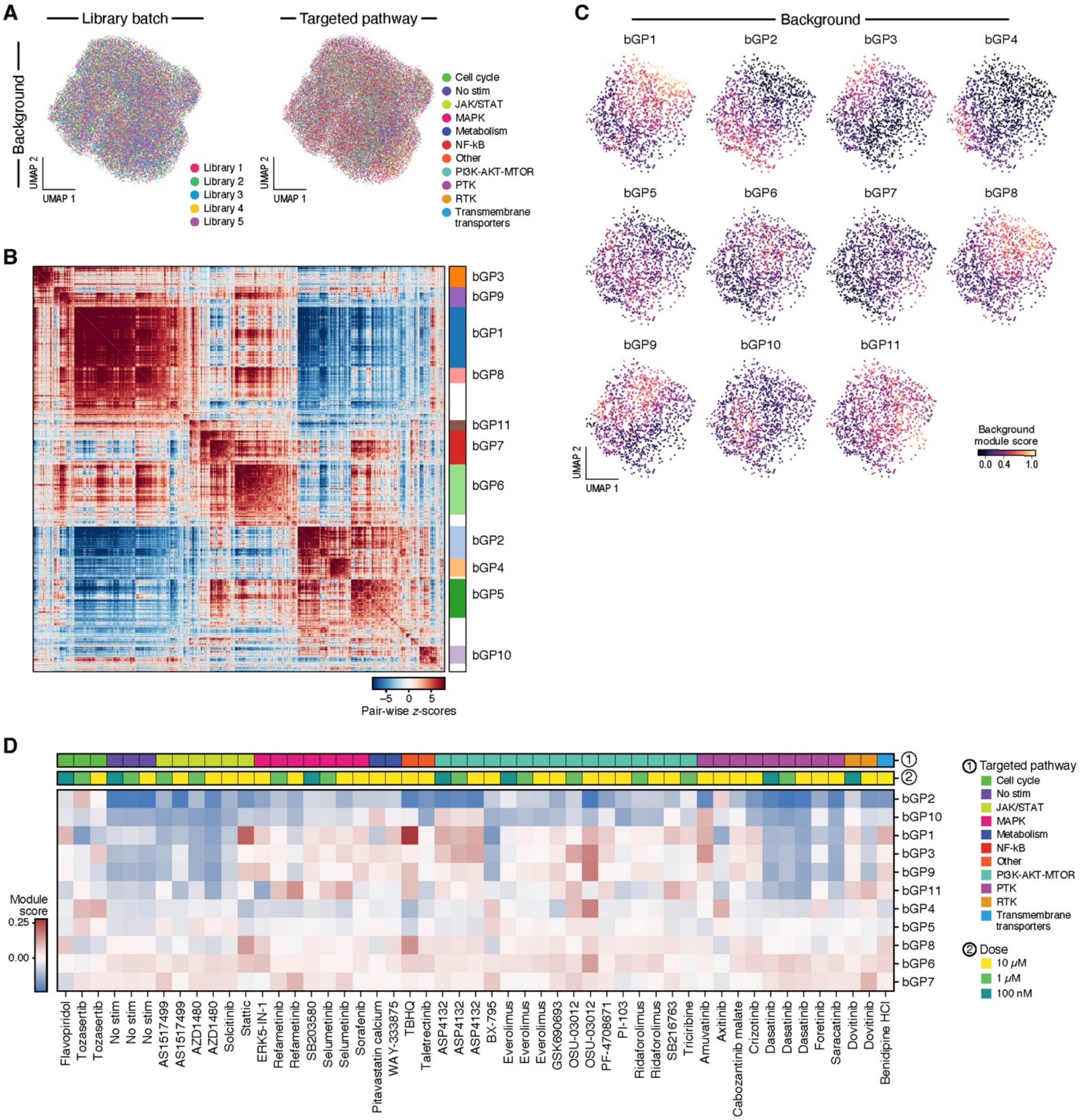


**Fig S4. Analyses of background gene programs, related to Fig. 3.**
(**A**) MoCAVI's background space. UMAP representation of icCITE-plex cell profiles (dots) in MoCAVI's background space, colored by library batch (left) or compound target pathway (right). (**B**-**D**) Background gene programs (bGPs). (B) Auto-correlation statistics computed using Hotspot (pairwise *z*-scores; color scale) between expression profiles weighted by distances in MoCAVI's background space for each genes (rows, columns), ordered by hierarchical clustering (average linkage) and annotated by background gene program membership (right, color legend). (C) UMAP representation as in (A) colored by score (color bar) of each bGP. (D) Scores (intensity color bar) for each bGP (rows) in pseudobulk profiles (columns) for the SM treatments with the strongest impact in the background space, related to **Fig. 3C**, with treatments ordered by drug-targeted pathway (bar 1) and annotated by dose (bar 2).

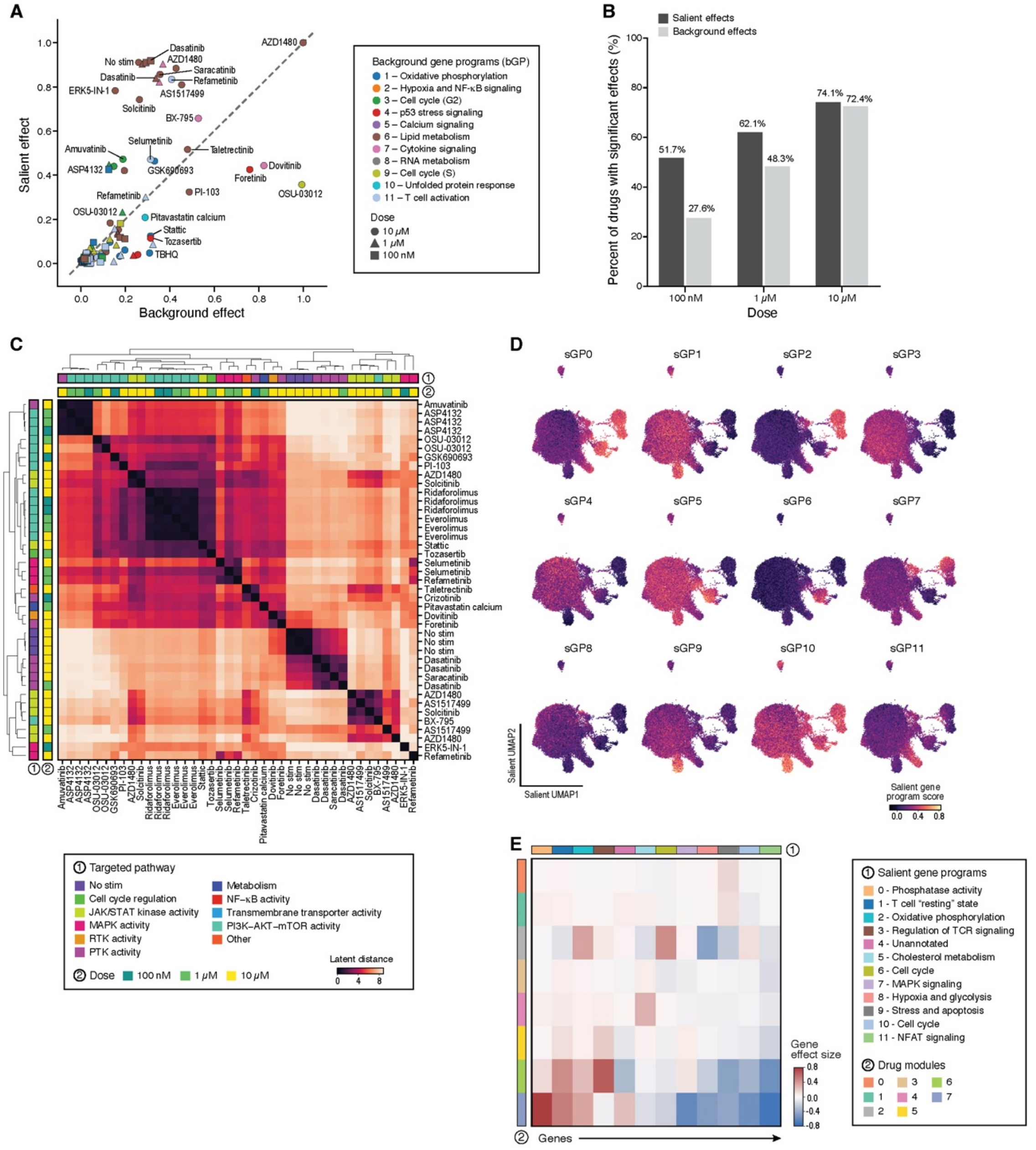

**Fig. S5. Analyses of salient gene programs, related to Fig. 4.**
(**A,B**) Salient effects are stronger and more prevalent than background effects for most SM perturbations. (A) MoCAVI background (x axis) and salient (y axis) effect scores for each SM treatment (points, as in **Fig. 4A**), marked by dose (shape) and highest-scoring background gene expression program (color legend)). (B) Prevalence of significant perturbation effects across the salient and background latent spaces. Proportion of drugs (y axis) in each dose (x axis) with significant effects in the salient (black) or background (grey) latent space. (**C**) Similarity of perturbations in the salient space. Euclidean distance (color bar) between SM treatments (rows, columns, ordered by hierarchical clustering and annotated by the targeted pathway (bar 1) and dose (bar 2)), based on mean aggregation of MoCAVI's salient latent space at the perturbation level. (**D**,**E**) Salient gene programs (sGPs). (D) UMAP representation of cell profiles (dots) in MoCAVI's salient space, colored by score (color bar) of each sGP. (E) Mean-aggregated log-fold changes (color bar) inferred by MoCAVI in the salient space for each sGP (columns, bar 1) in each drug module (rows, bar 2).

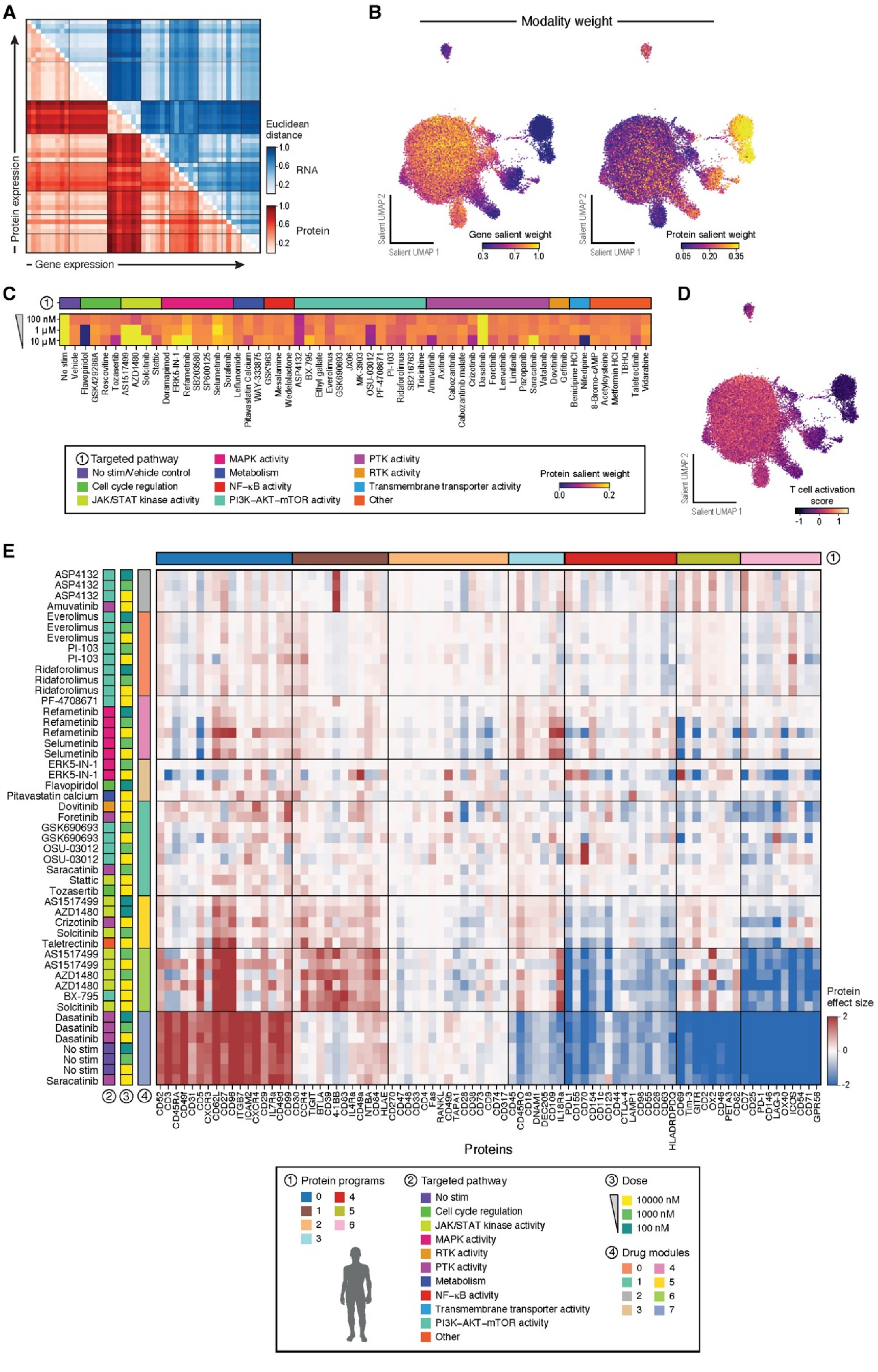
A
Protein expression
Gene expression
Euclidean distance
1.0
0.6
0.2
RNA
Protein
B
Modality weight
Salient UMAP 2
Salient UMAP 1
Gene salient weight
0.3 0.7 1.0
Protein salient weight
0.05 0.20 0.35
C
100 nM
1 µM
10 µM
No stim
Vehicle
Flavopiridol
GSK429286A
Roscovitine
Tozasertib
AS1517499
AZD1480
Solcitinib
Stattic
Doramapimod
ERK5-IN-1
Refametinib
SB203580
SP600125
Selumetinib
Sorafenib
Leflunomide
Pitavastatin Calcium
WAY-333875
GSK983
Mesalamine
Wedelolactone
ASP4132
BX-795
Ethyl gallate
Everolimus
GSK690693
JX06
MK-3903
OSU-03012
PF-4708671
PI-103
Ridaforolimus
SB216763
Triciribine
Amuvatinib
Axitinib
Cabozantinib
Cabozantinib malate
Crizotinib
Dasatinib
Foretinib
Lenvatinib
Linifanib
Pazopanib
Saracatinib
Vatalanib
Dovitinib
Gefitinib
Benidipine HCl
Nifedipine
8-Bromo-cAMP
Acetylcysteine
Metformin HCl
TBHQ
Taletrectinib
Vidarabine
① Targeted pathway
No stim/Vehicle control
Cell cycle regulation
JAK/STAT kinase activity
MAPK activity
Metabolism
NF-κB activity
PI3K–AKT–mTOR activity
PTK activity
RTK activity
Transmembrane transporter activity
Other
Protein salient weight
0.0 0.2
D
Salient UMAP 2
Salient UMAP 1
T cell activation score
-1 0 1
E
ASP4132
ASP4132
ASP4132
Amuvatinib
Everolimus
Everolimus
Everolimus
PI-103
PI-103
Ridaforolimus
Ridaforolimus
Ridaforolimus
PF-4708671
Refametinib
Refametinib
Refametinib
Selumetinib
Selumetinib
ERK5-IN-1
ERK5-IN-1
Flavopiridol
Pitavastatin calcium
Dovitinib
Foretinib
GSK690693
GSK690693
OSU-03012
OSU-03012
Saracatinib
Stattic
Tozasertib
AS1517499
AZD1480
Crizotinib
Solcitinib
Taletrectinib
AS1517499
AS1517499
AZD1480
AZD1480
BX-795
Solcitinib
Dasatinib
Dasatinib
Dasatinib
No stim
No stim
No stim
Saracatinib
② ③ ④
CD52
CD3
CD45RA
CD49f
CD31
CD5
CXCR3
CD62L
CD27
CD96
ITGB7
ICAM2
CXCR4
CD29
IL7Ra
CD49d
CD99
CD30
CCR4
TIGIT
BTLA
CD39
4-1BB
CD83
IL4Ra
CD49a
NTBA
CD84
HLAE
CD270
CD47
CD48
CD33
CD4
Fas
RANKL
CD49b
TAPA1
CD28
CD38
CD9
CD74
CD317
CD45
CD45RO
CD18
DNAMI
DEC205
CD109
IL18Ra
PDL1
CD155
CD70
CD154
CD11c
CD123
CD44
CTLA-4
LAMP1
CD98
CD55
CD26
CD63
HLADRDPDQ
CD69
Tim-3
GITR
CX2
CD46
PETA3
CD82
CD7
CD25
PD-1
CD146
LAG-3
OX40
ICOS
CD54
CD71
GPR56
Proteins
Protein effect size
2
0
-2
① Protein programs
0 1 2 3 4 5 6
② Targeted pathway
No stim
Cell cycle regulation
JAK/STAT kinase activity
MAPK activity
RTK activity
PTK activity
Metabolism
NF-κB activity
Transmembrane transporter activity
PI3K–AKT–mTOR activity
Other
③ Dose
10000 nM
1000 nM
100 nM
④ Drug modules
0 1 2 3 4 5 6 7

**Fig. S6. Characterization of protein features in MoCAVI's salient space, related to Fig. 4.**
(**A**) Agreement in perturbation similarity based on RNA and protein changes. Euclidean distances between SM treatments (rows, columns) based on salient-specific log-fold change profiles in either RNA(summed across genes, upper triangle, blue/white color bar) or protein (summed across proteins, lower triangle, red/white) measurements. Treatments are ordered by single-linkage hierarchical clustering in MoCAVI's salient latent space aggregated at the perturbation level and integrating both modalities. (**B-D**) MoCAVI protein salient weights are enriched in 'naive-like' cells. (B) UMAP representation of cell profiles (dots) in MoCAVI's salient space, colored by MoCAVI's RNA (left) and protein (right) salient weights (color bar). (C) MoCAVI's protein salient weights (color bar) for pseudobulk profiles of SM treatments (columns, annotated by drug target pathway) in increasing doses (rows, from top). (D) UMAP representation of cell profiles (dots) in MoCAVI's salient latent space, colored by gene signature scores of T cell activation (color bar). (**E**) Co-regulated protein programs. Log-fold changes of select protein markers (columns, **Table S4,** clustered into protein programs, bar 1) across SM treatments (rows, ordered by drug modules (as defined in **Fig. 4E**, bar 4) and annotated by drug target pathway (bar 2), dose (bar 3) and protein programs (bar 1).

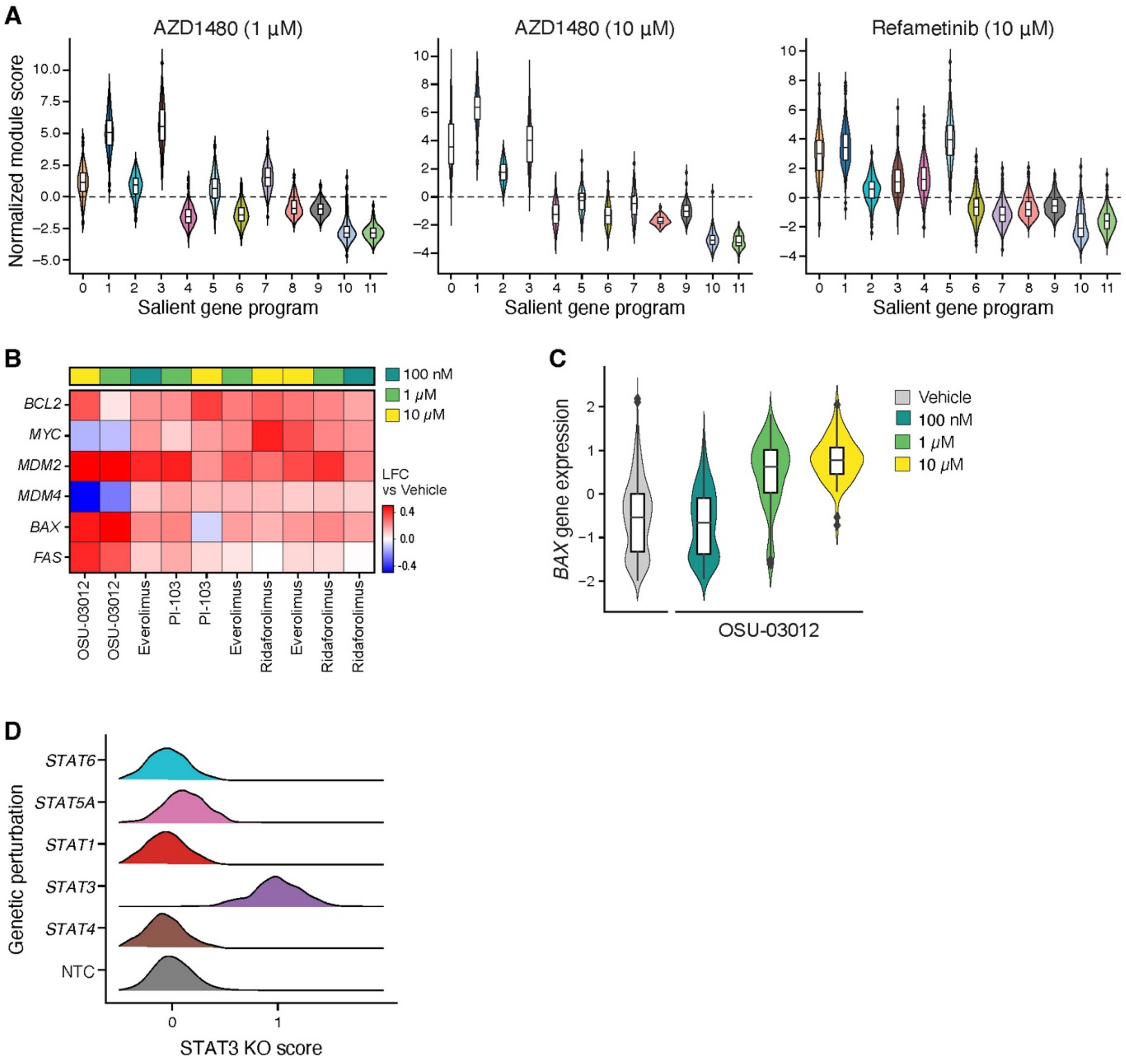


**Fig. S7. SM treatment-induced phenotypes, related to Fig. 4.**

(**A**) Salient gene program activity for select SM treatments. Distribution of normalized module scores (y axis) for each salient gene program (x axis) in cells treated by AZD1480 (left, middle) or refametinib (right). (**B**,**C**) OSU-03012 induces a pro-apoptotic RNA expression program. (B) Log fold-change (color) in expression of select apoptosis-related genes (rows) relative to vehicle across SM treatments (columns) and doses (top bar). (C) Distribution of *BAX* RNA expression levels (y axis) in individual cells following treatment with OSU-03012 (x axis), colored by dose (color bar). (**D**) STAT3 KO module scores across select STAT family knockouts. Distributions of STAT3 KO module scores (x axis) in individual cells in different STAT family genetic perturbations (y axis).

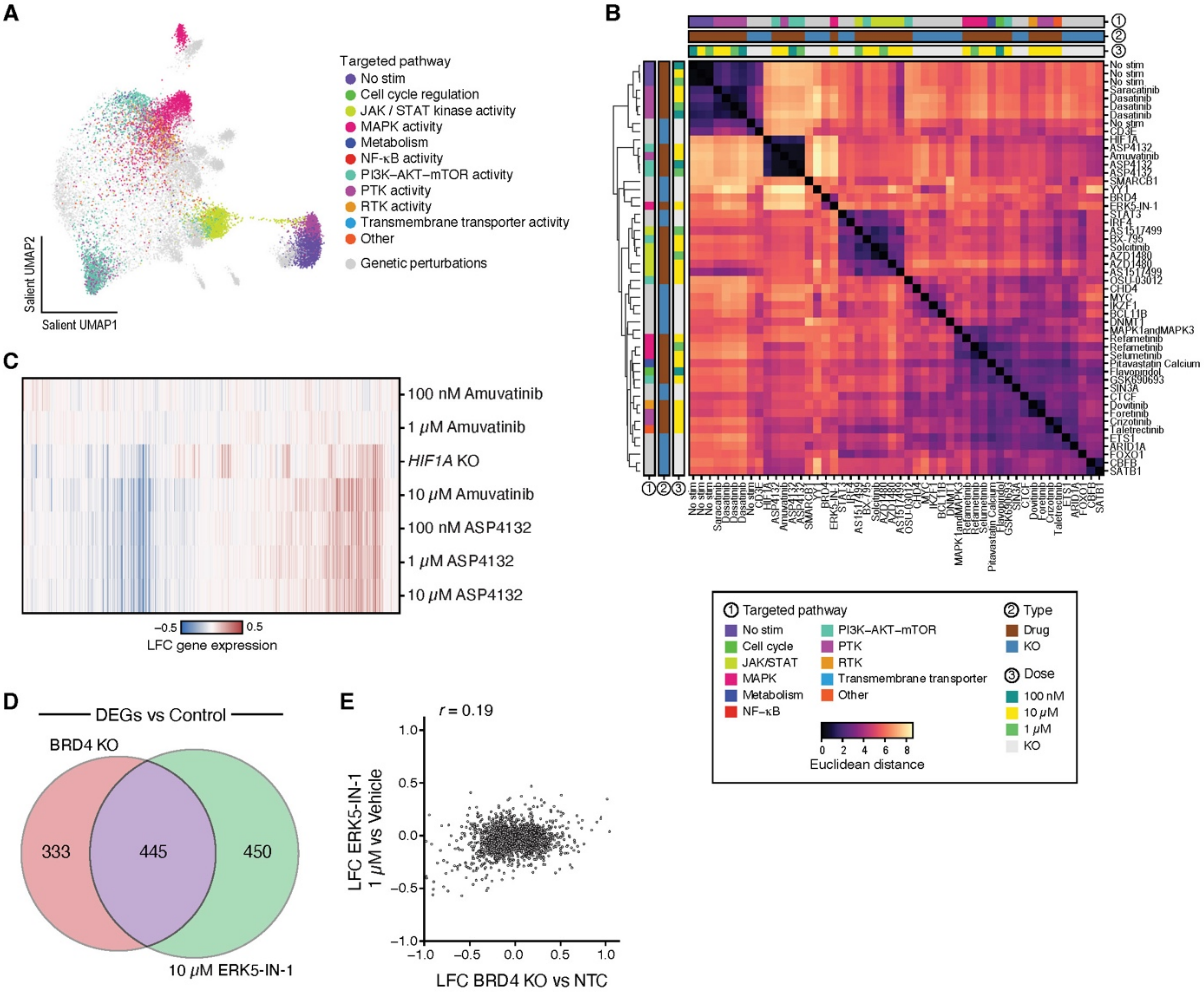


**Fig. S8. Chemical and genetic perturbation effects, related to Fig. 5.**
(**A,B**) Joint analysis of chemical and genetic perturbations. (A) UMAP representation of single cell profiles (dots) in MoCAVI's joint salient space (as in **Fig. 5B**), colored by drug-targeted pathway. (B) Euclidean distance between SM treatments and genetic knockout mean aggregated profiles (rows, columns) in MoCAVI's salient latent space, ordered by hierarchical clustering and annotated by perturbation type (bar 2), drug targeted pathway (bar 1) and dose (bar 3). (**C**-**E**) Phenotypic similarities between responses to gene knockouts and SM treatments. (C) Log fold-change (LFC, color) in expression of each gene (columns) under each perturbation (rows) relative to control (Vehicle or NTC). (D) Overlap of differentially expressed genes between BRD4 KO and ERK5-in-1 treatment relative to control (NTC or Vehicle). (E) Log fold change (y and x axes) in expression of each gene (dot) in BRD4 knockout vs. NTC (x axis) and 1 μM ERK5-IN-1 vs Vehicle (y axis). No significant correlation was observed at this dose.

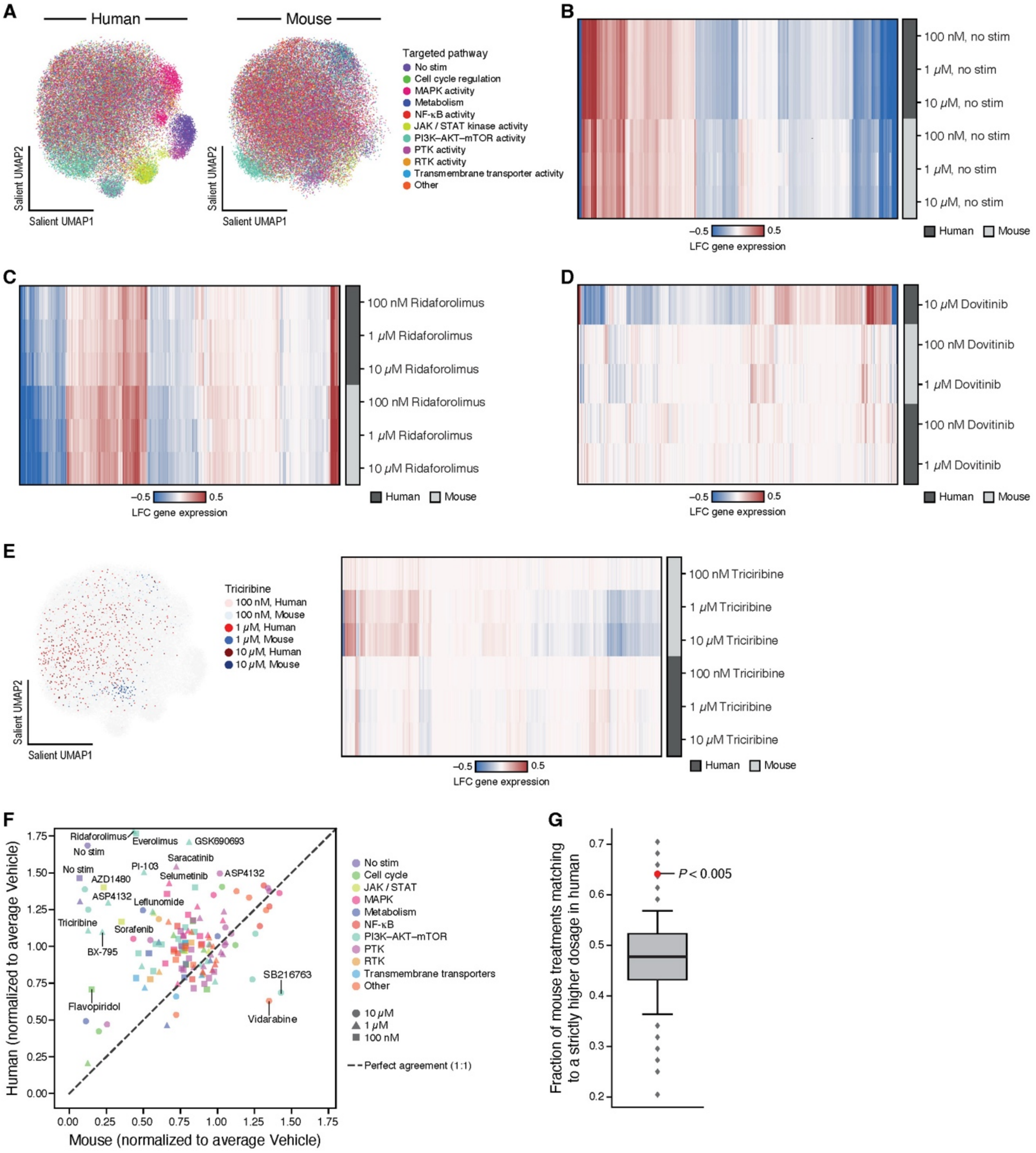


**Fig. S9. icCITE-plex cross-species analyses, related to Fig. 5.**
(**A**) Cross species salient latent space. UMAP representation of icCITE-plex multimodal profiles (dots) of human (left) or mouse (right) cells in the cross-species salient latent space (as in **Fig. 5H**), colored by the drug-targeted pathway. (**B**-**E**) Similarity and distinction of species responses to SM treatment. Log fold-change (LFC, color) in the expression of each gene (columns) for human (dark grey) and mouse (light grey) cells under non-stimulated conditions (B, rows) or following treatment with indicated doses of ridaforolimus (C, rows), dovitinib (D, rows) or triciribine (E, right) relative to vehicle control. (E, left) UMAP representation as in A, highlighting cells treated by triciribine in each species and dose (color legend). (**F**) Reduced cell viability in mouse cells for some SM treatments. Normalized recovered cell numbers for each SM treatment (points, annotated by targeted drug pathway (color) and dose (shape)) in mouse (x axis) and human (y axis) samples. (**G**) Assessment of dosage-consistent cross-species similarity. Fraction of drug perturbations in mouse (y axis) that are more closely related in the joint MoCAVI salient latent space to the same drug at higher dosage in human than to lower dosage (y axis). Red point: observed statistic; gray distribution: empirical null generated by shuffling dose labels (**Methods**).

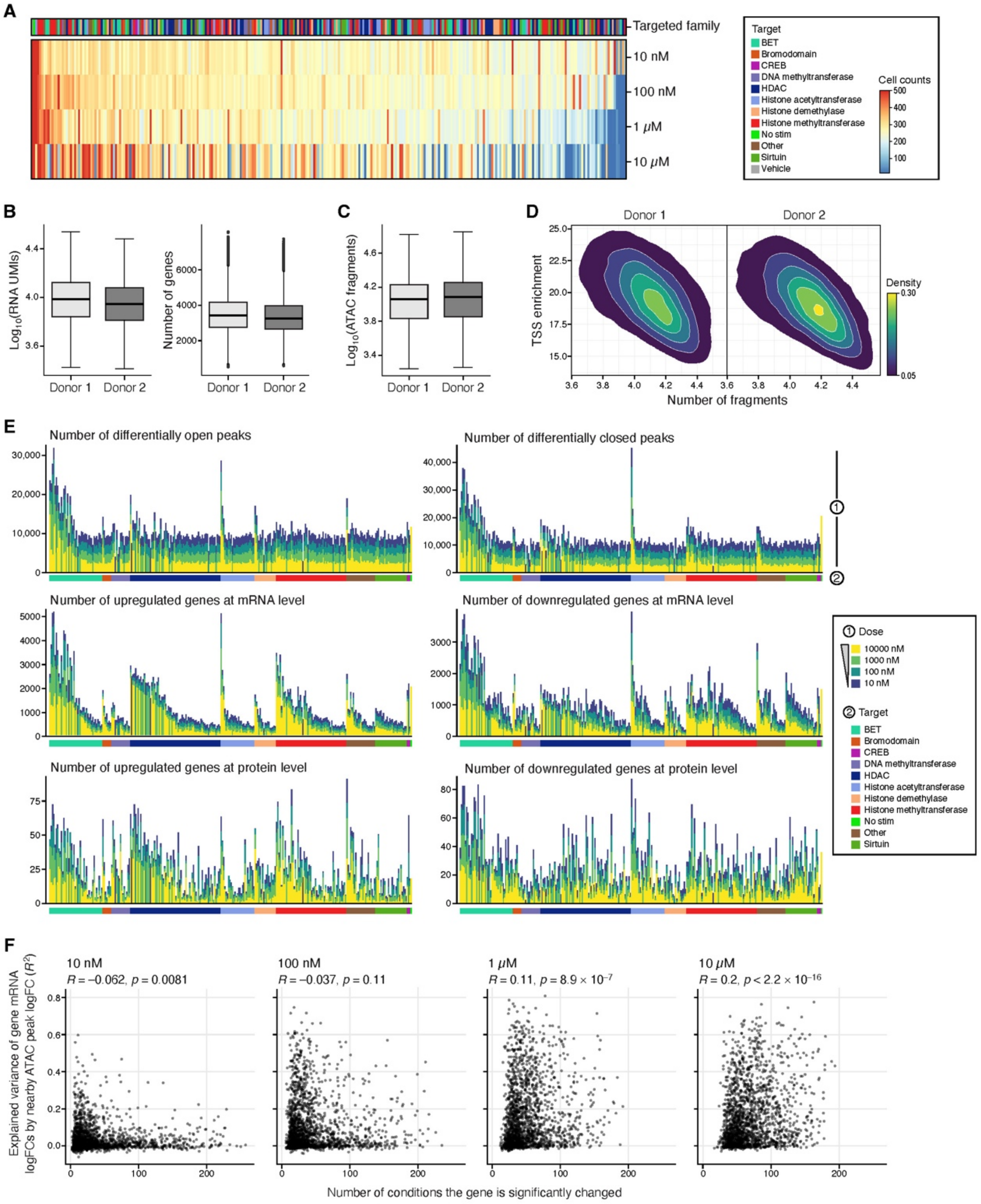


**Fig. S10. Quality metric analyses for DOGMA-plex, related to Fig. 6.**
(**A-D**) DOGMA-plex experiment quality control metrics. (A) Number of cells recovered (color bar) from each drug (columns, annotated by drug target, top bar) in each dose (rows, increasing from top). (B,C) Number of mRNA UMIs recovered per cell (B, left, y axis), number of genes detected per cell (B, right, y axis), number of ATAC fragments recovered per cell (C, y axis) for each donor (x axis). Boxplots denote three quartiles, distribution with whiskers, and outliers as dots. (D) Distribution of unique ATAC-seq nuclear fragments (x axis) and transcription start site (TSS) enrichment (y axis) across single cells, colored by local point density (color bar). (**E**) Multimodal responses across drug treatments. Number of differentially accessible peaks (top) and differentially expressed genes (middle) and proteins (bottom) in each drug drug dose (stack color) of each SM (x axis, sorted by drug target (bar 2)). (**F**) Relationship between mRNA changes and chromatin accessibility. Number of conditions with significant mRNA changes (FDR < 0.1, x axis, **Methods**) and the variance in mRNA explained by nearby ATAC peak accessibility (y axis) for each drug treatment (dot) in each dose (panels).

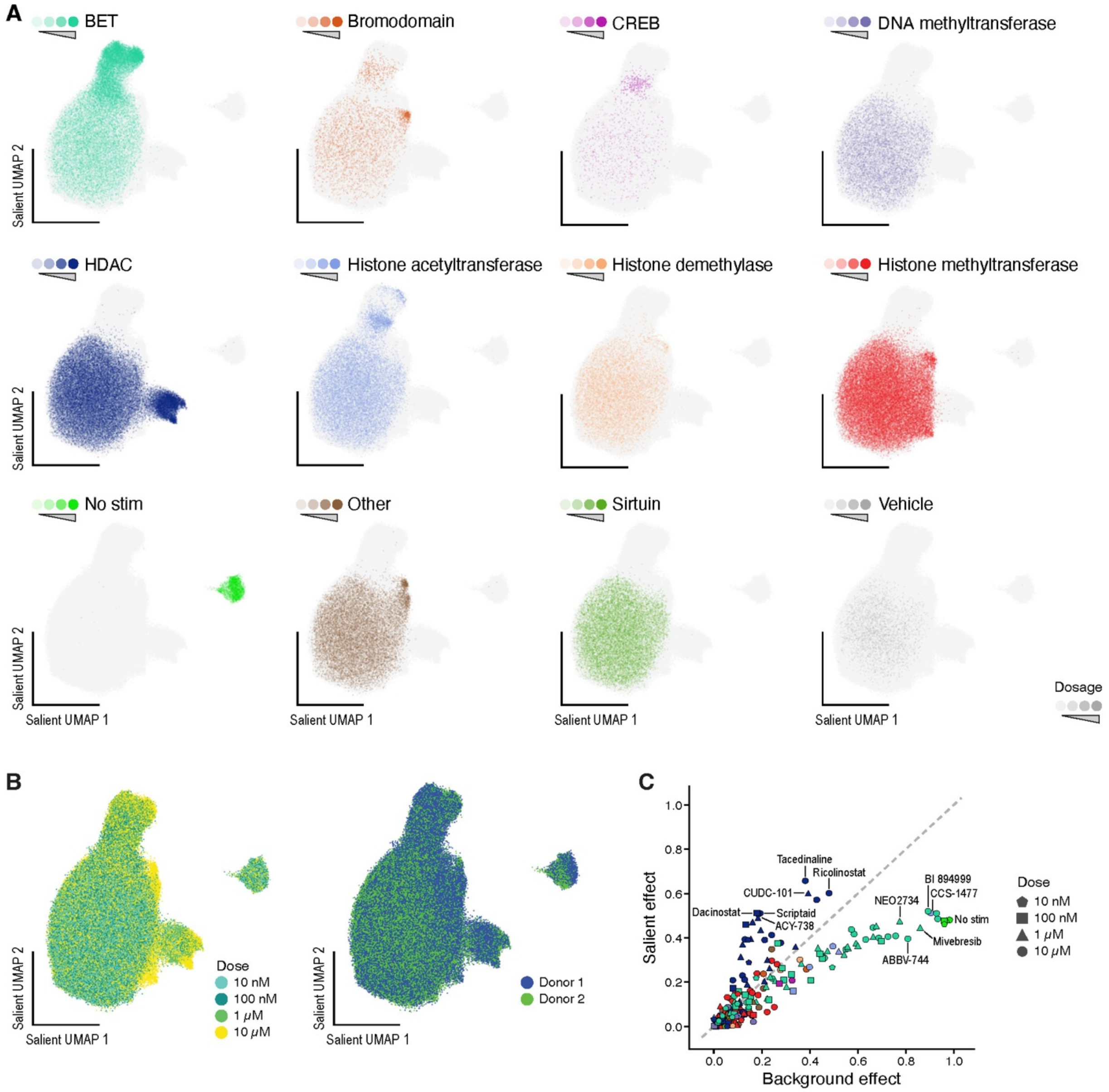


**Fig. S11. MoCAVI's salient space in DOGMA-plex, related to Fig. 6.**
(**A,B**) MoCAVI representation of the DOGMA-plex salient latent space. UMAP representation of cell profiles (dots) in the MoCAVI salient latent space, colored by drug target and dose combination (A), dose (across all drugs, B, left) or donor (B, right). (**C**) Salient and background effects. MoCAVI background (x axis) and salient (y axis) effect scores for each SM treatment, annotated by dose (shape) and drug target (color, code as in **Fig. 6B**).

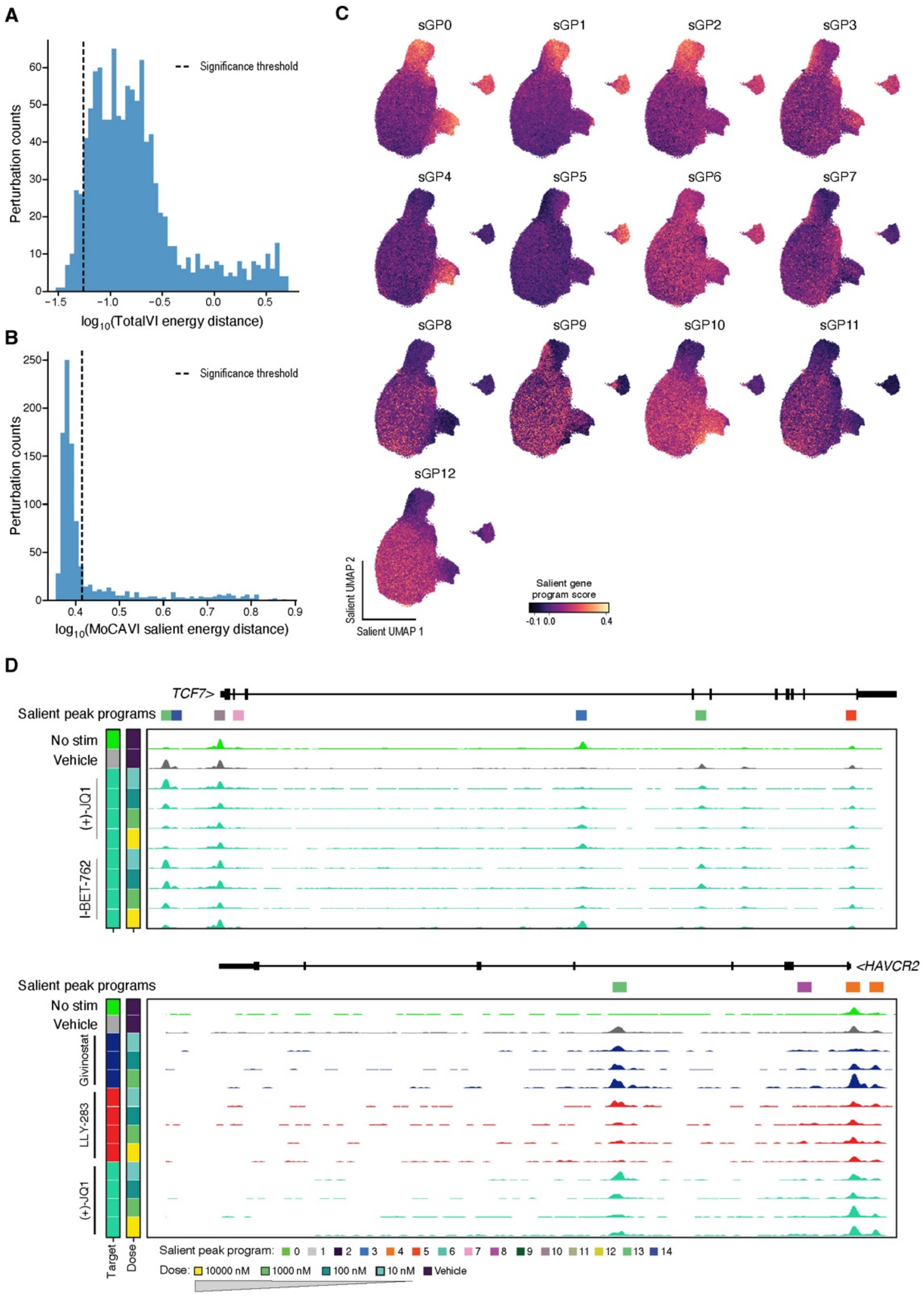


**Fig. S12. DOGMA-plex salient gene and peak programs, related to Fig. 6.**
(**A,B**) Impactful perturbations. Distribution of energy distances (x axis) between control and drug-treated cell profiles in the iTotalVI (A) and MoCAVI (B) latent spaces. Dashed line in A: significance threshold (P<0.05, empirical p-value, **Methods**). In B, significant perturbations were further classified by their effect magnitude in MoCAVI's salient space, with upper-tail defined as strong-effect perturbations and the remaining significant perturbations considered weak-effect. Dashed line in B: strong-effect threshold. (**C**) Salient gene programs. UMAP representation of cell profiles (dots) in MoCAVI's salient space, colored by the score of each sGP. (**D**) Chromatin regions differentially accessible between drug-treated and vehicle cells at key T cell activation-related loci. Pseudo-bulk ATAC reads at the *TCF7* (top) or *HAVCR2* (bottom) locus for select treatments (rows, annotated by drug target (as in **Fig. 6B**) and dose) annotated by salient peak programs (color bar).

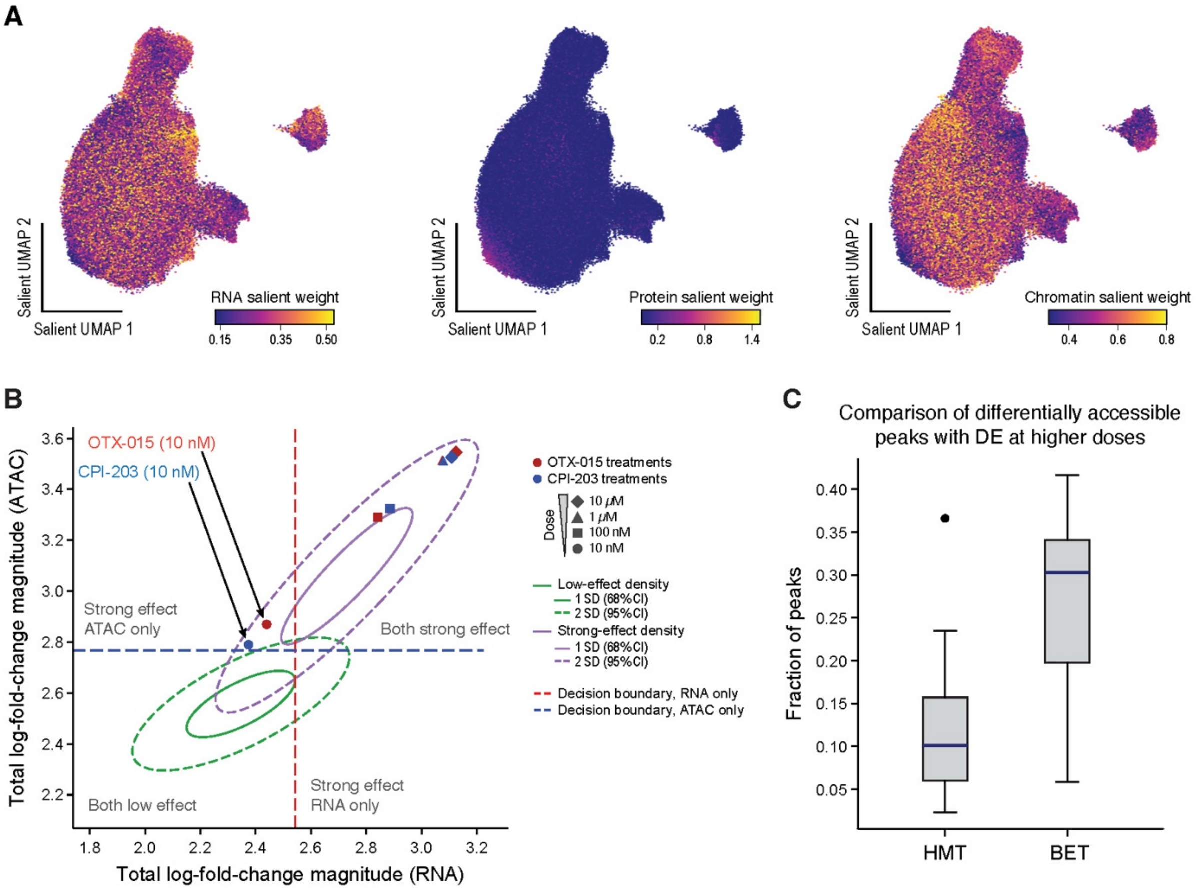


**Fig. S13. MoCAVI's DOGMA-plex salient space, related to Fig. 6.**
(**A**) Salient weights for different modalities. UMAP representation of cell profiles (dots) in MoCAVI's salient space, colored by MoCAVI's RNA (left), protein (middle), or chromatin (right) salient weights (color bar). (**B,C**) At lower doses, chromatin accessibility captures the effects of some BET inhibitors better than RNA expression. (B) Total log-fold-change in RNA (x axis) and chromatin accessibility (y axis) for drug-dose combination (dots) of OTX-015 (red) and CPI-203 (blue) (annotated by dose (shape)). Ovals: 68% (solid oval) and 95% (dashed oval) probability contours for the density of perturbation classes (weak-effect, green; strong-effect, purple; assessed globally across modalities in **fig. S12B**). Dotted vertical (blue) and horizontal (red) lines indicate the decision boundaries (50% probability contour) of linear discriminant analysis models trained separately on RNA and chromatin accessibility effect sizes using these global labels. This analysis projects the global perturbation classes into RNA and chromatin accessibility space. (C) Fraction of differentially accessible peaks (y axis) that are linked to genes that are differential expressed at higher drug doses for HMT and BET inhibitors (x axis).

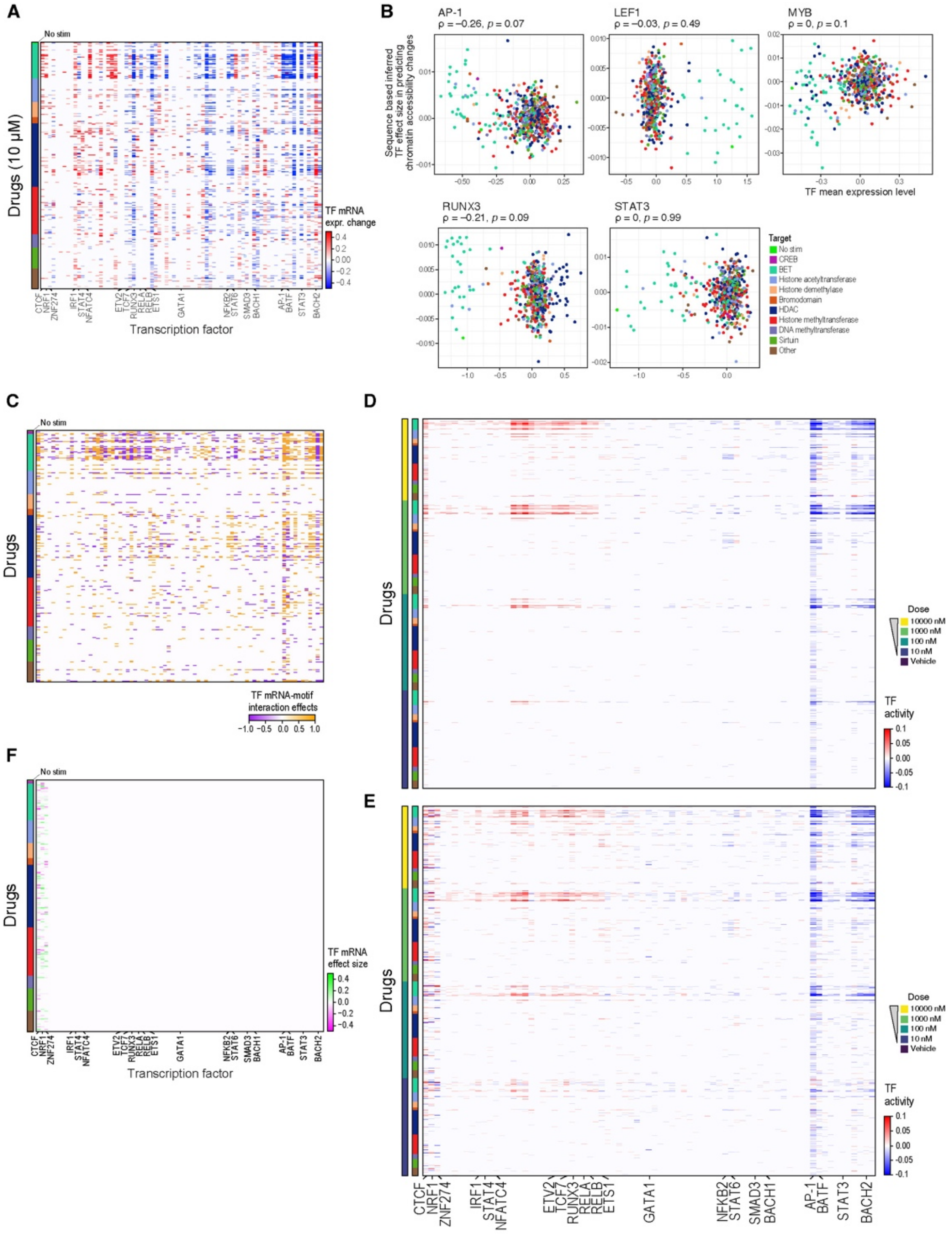


**Fig. S14. Impact of incorporating TF mRNA levels in regulatory predictions, related to Fig. 7.**
(**A**) TF mRNA expression changes. RNA expression changes (FDR < 0.1, **Methods**) for each of 77 TF genes (columns) in each 10 µM drug treatment (rows, annotated by drug target, left bar, color code as in **B**). (**B**) Lack of association between TF activity and permuted motif accessibility. Mean TF RNA level (x axis) and TF activity inferred from motif accessibility in permuted data (y axis) in each drug–dose combination (points, colored by drug target). (**C**) Significant interactions between TF mRNA levels and motif accessibility across drug doses. TF mRNA–motif interaction effects on chromatin accessibility (color bars) for each TF (columns, labeled in **F**) in each drug treatments (rows, sorted by drug target, color code in **B**). White: FDR ≥ 0.1. (**D,E**) Integrating TF motifs, chromatin accessibility and TF mRNA increases TF activity prediction. Estimated activity scores (color bars) for each TF (columns) in each drug treatment (rows, ordered by dose, drug target, colored as in **B**) at all doses based on motif-accessibility relationship only (D) or when also incorporating TF mRNA expression (E). (**F**) CTCF, NRF1 and ZNF274 mRNA levels across drug doses are predictive of chromatin accessibility changes independent of their motif sites. Effect of TF mRNA levels on chromatin accessibility (color bar) for each TF (columns) across doses of each drug (rows, sorted by drug-target pathway, color-coded as in **B**).

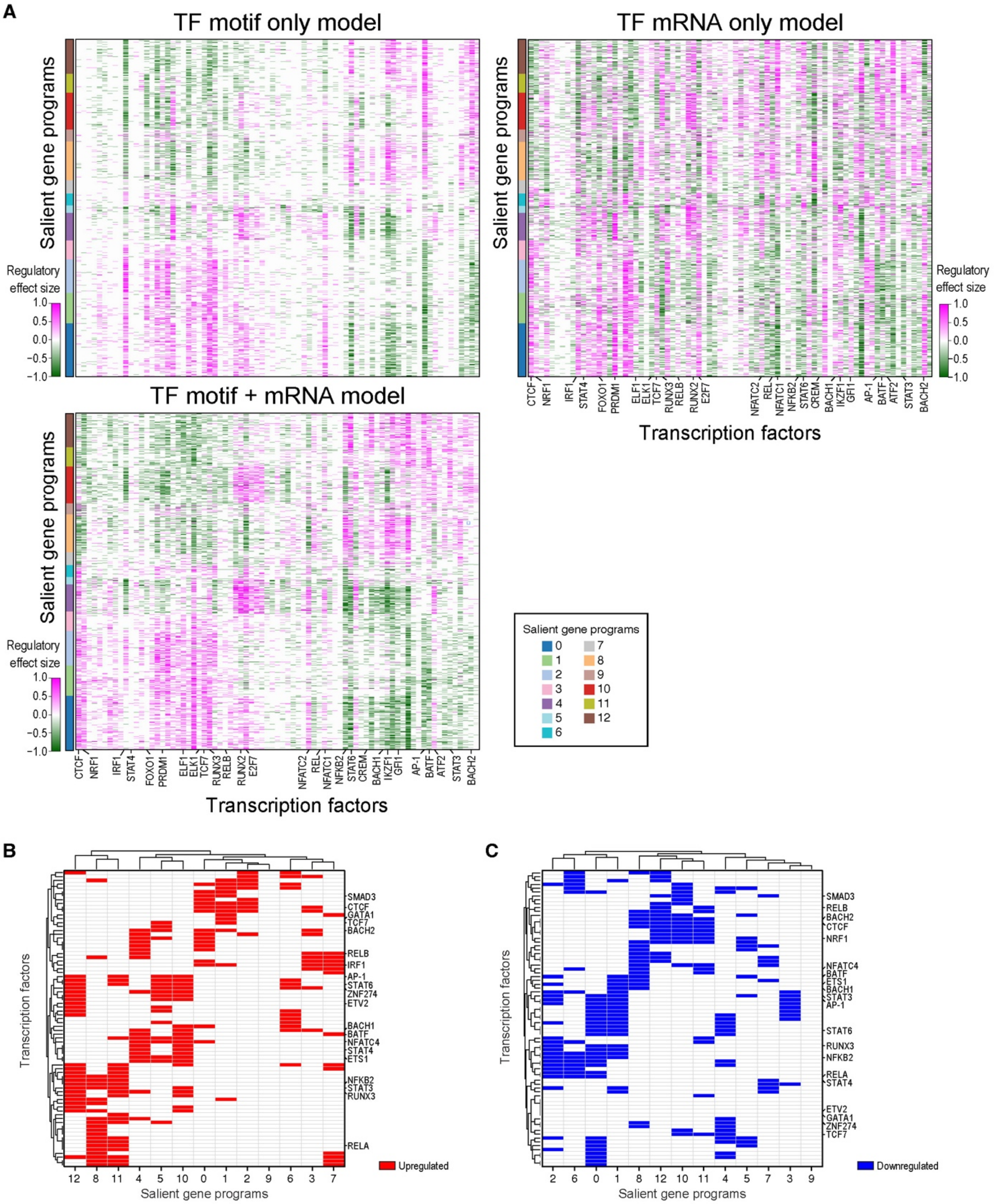


**Fig. S15. Motif-gated regression links TF activity to gene programs, related to Fig. 7.**
(**A**) TF motif activity and mRNA yield a richer regulatory model. Regulatory effect size (activator (pink)/repressor (green) color bar; regularized regression; **Methods)** of each TF (columns) on each gene (rows), ordered by MoCAVI sGPs (left bar), using TF motif activity scores only (top left), TF expression levels only (top right) or both (bottom, as in Fig. 7F top left), conditioned on the presence of corresponding TF motifs in regulatory regions. (**B,C**) Linking TF activity to sGPs. Enrichment (FDR < 0.1, Fisher's exact test) in inferred upregulated (B, red) or downregulated (C, blue) target genes from each MoCAVI sGPs in inferred target genes of each TF (rows).

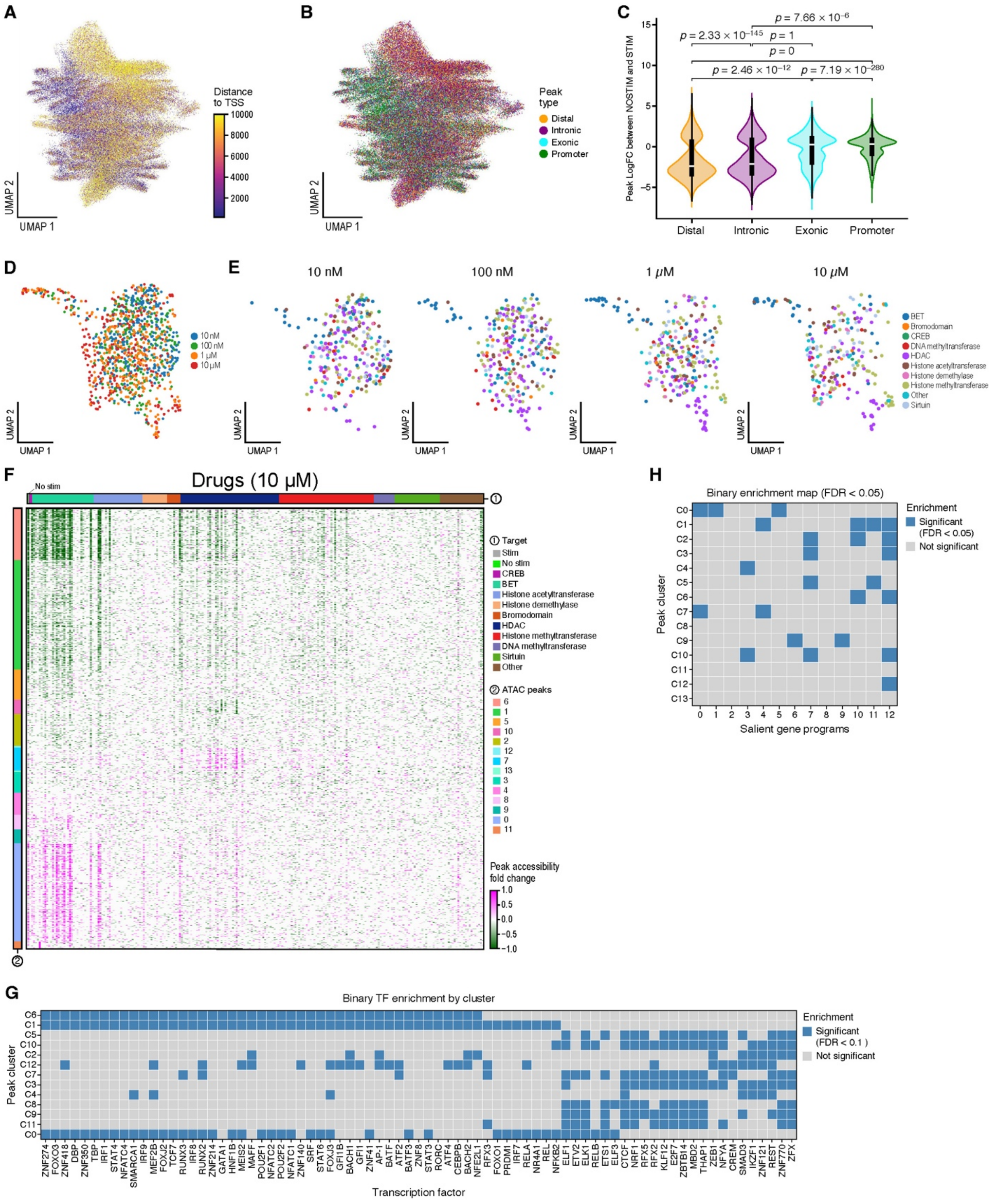

A
Distance to TSS
10000
8000
6000
4000
2000
UMAP 2
UMAP 1
B
Peak type
Distal
Intronic
Exonic
Promoter
UMAP 2
UMAP 1
C
Peak LogFC between NOSTIM and STIM
p = 7.66 × 10^−6
p = 2.33 × 10^−145
p = 1
p = 0
p = 2.46 × 10^−12
p = 7.19 × 10^−280
Distal
Intronic
Exonic
Promoter
D
10 nM
100 nM
1 μM
10 μM
UMAP 2
UMAP 1
E
10 nM
100 nM
1 μM
10 μM
BET
Bromodomain
CREB
DNA methyltransferase
HDAC
Histone acetyltransferase
Histone demethylase
Histone methyltransferase
Other
Sirtuin
F
Drugs (10 μM)
No stim
① Target
Stim
No stim
CREB
BET
Histone acetyltransferase
Histone demethylase
Bromodomain
HDAC
Histone methyltransferase
DNA methyltransferase
Sirtuin
Other
② ATAC peaks
6
1
5
10
2
12
7
13
3
4
8
9
0
11
Peak accessibility fold change
1.0
0.5
0.0
−0.5
−1.0
H
Binary enrichment map (FDR < 0.05)
Enrichment
Significant (FDR < 0.05)
Not significant
Peak cluster
C0
C1
C2
C3
C4
C5
C6
C7
C8
C9
C10
C11
C12
C13
0 1 2 3 4 5 6 7 8 9 10 11 12
Salient gene programs
G
Binary TF enrichment by cluster
Enrichment
Significant (FDR < 0.1)
Not significant
Peak cluster
C6
C1
C5
C10
C2
C12
C7
C3
C4
C8
C9
C11
C0
ZNF274 FOXO3 ZNF418 DBP ZNF350 TBP IRF1 STAT4 NFATC4 SMARCA1 IRF9 MEF2B FOXJ2 TCF7 RUNX3 IRF8 RUNX2 ZNF214 GATA1 HNF1B MEIS2 MAFF POU2F1 NFATC2 POU2F2 NFATC1 ZNF140 SRF STAT6 FOXJ3 GFI1B BACH1 GFI1 ZNF41 AP-1 BATF ATF2 BATF3 ZNF8 STAT3 RORC ATF4 CEBPB BACH2 NFE2L1 RFX3 FOXO1 PRDM1 IRF7 RELA NR4A1 REL NFKB2 ELF1 ETV2 ELK1 RELB ETS1 ELF3 CTCF NRF1 RFX5 RFX2 KLF12 E2F7 ZBTB14 MBD2 THAP1 ZEB1 NFYA CREM SMAD3 IKZF1 ZNF121 REST ZNF770 ZFX
Transcription factor

**Fig. S16. Annotation and clustering of perturbation-induced ATAC-seq peaks, related to Fig. 8.**
(**A, B**) Distal and intronic regions show the greatest differential accessibility between vehicle and drug-dose conditions. UMAP representation of accessibility changes relative to the vehicle control in each ATAC-seq peak region (dots, as in **Fig. 8A-G**), colored by distance to the closest gene's transcription start site (TSS) (A) or its genomic feature (B). (**C**) Distal and intronic regions show the greatest differential accessibility under stimulation conditions. Distribution of accessibility log fold-changes in stimulated vs. unstimulated (vehicle) conditions (y axis) of peaks in different genomic regions (x axis) that are differentially accessible between stimulated and unstimulated conditions. P-values: One-sided Wilcoxon test. (**D,E**) Drug-dose effects on chromatin accessibility that are independent of T cell activation are shared across members of the same drug family. UMAP representation of embeddings of drug-dose treatments (dots) based on chromatin accessibility changes at peak regions, colored by dose (D) or drug target (E). (**F**) Chromatin accessibility peak clusters. Differential accessibility (color bar) of peaks (rows, ordered by peak clusters (bar 2)) that are significantly differentially accessible in at least 10 treatments across each 10 μM drug treatment (columns, annotated by drug targets (bar 1)). (**G,H**) Linking functional loci to gene programs. Enrichment (blue, FDR < 0.1) of peak clusters (rows) in nearest genes assignments from each sGP (columns, G) or from inferred TF targets (columns, H) (**Methods**).

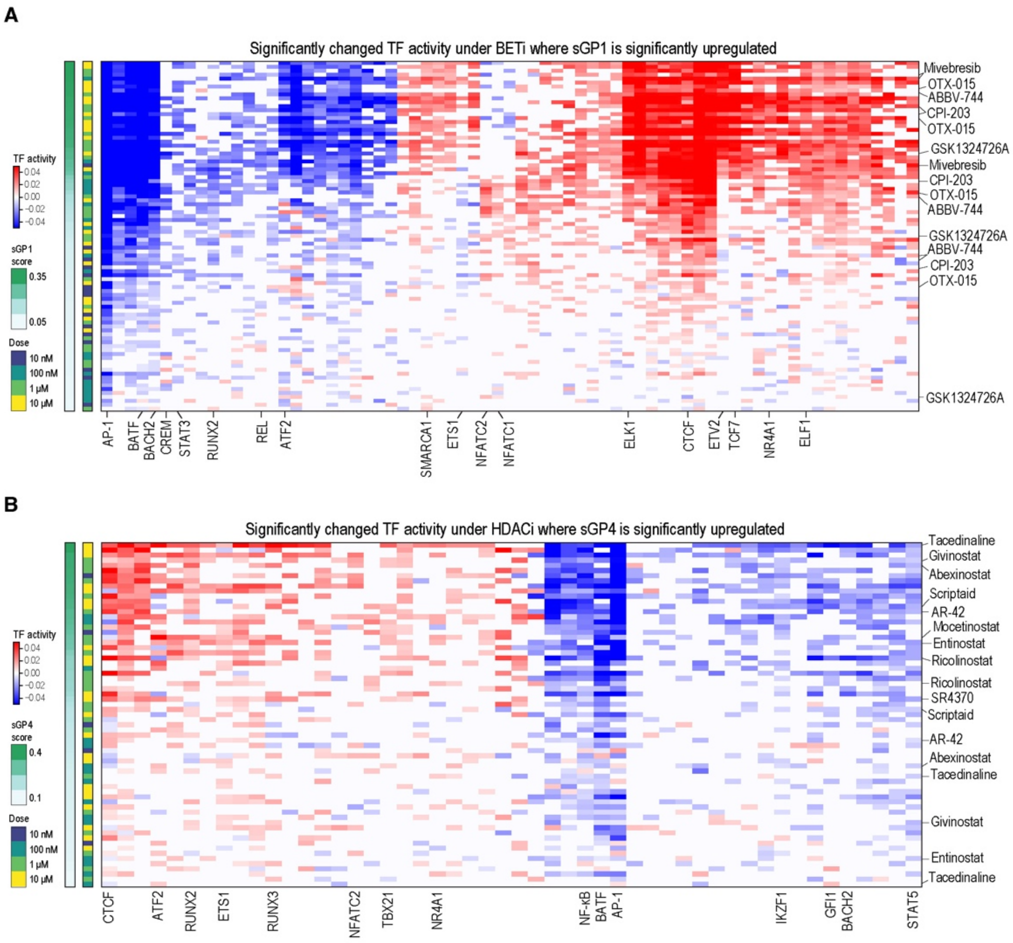


**Fig. S17. BETi and HDACi TF activity, related to Fig. 8.**
Activity scores (red/blue color bar) for each TF (columns) in pseudobulk profiles of select BETi (A) or HDACi (B) treatments (rows), ordered by decreasing sGP1 (A) or sGP4 (B) module scores (left most bar) and annotated by dose (second left bar).

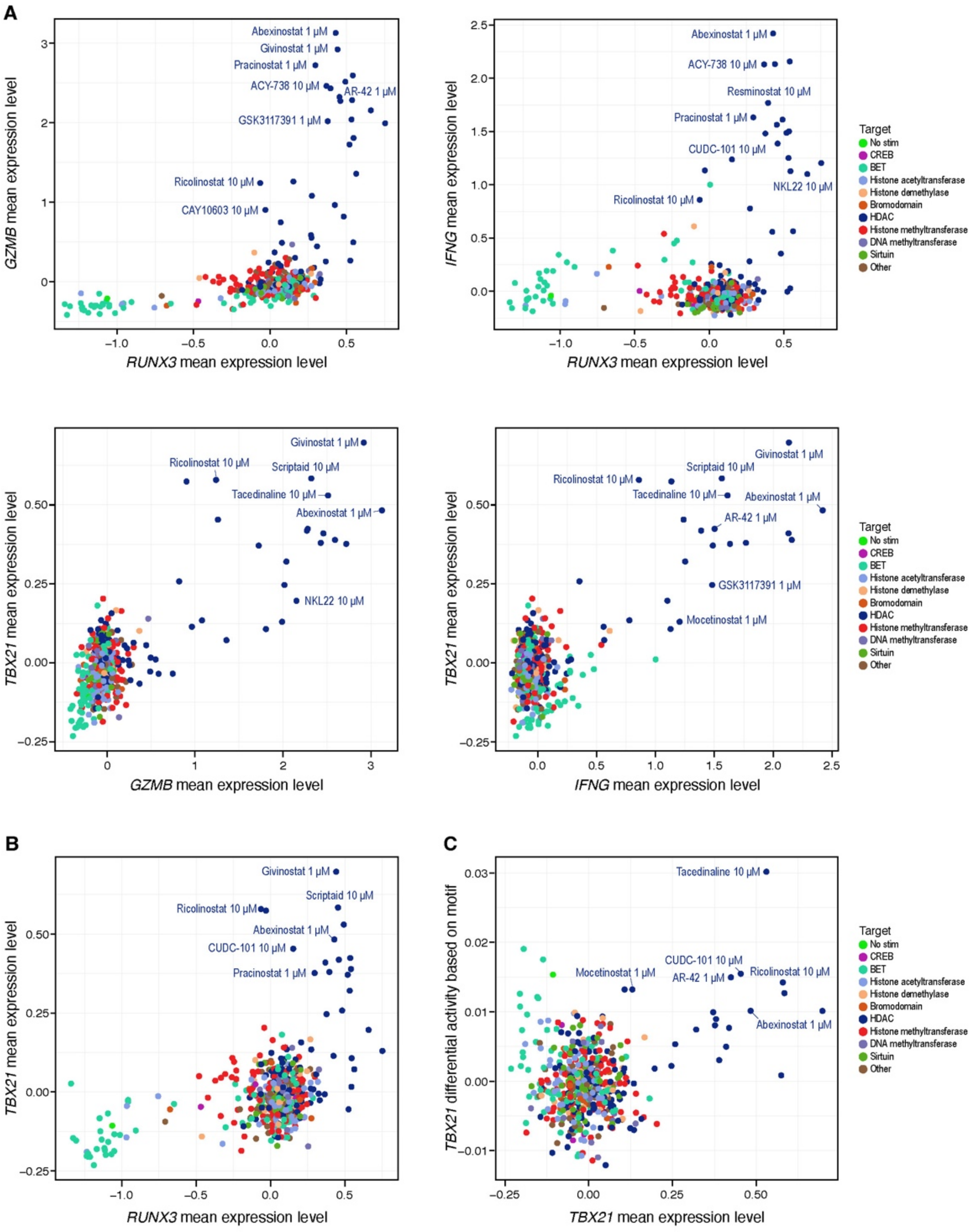


**Fig. S18. RUNX3 and TBX21 regulate proinflammatory cytokine expression, related to Fig. 8.**
**(A,B)** Mean mRNA expression (x and y axes) for indicated genes in each drug-dose combination (dots), colored by drug target (color bar). (**C**) Differential TF activity (vs. vehicle) (y axis) and mean mRNA expression (x axis) of TBX21 in each drug-dose combination (dots), colored by drug target (color bar).

**List of Supplementary Tables**

Table S1. icCITE-plex kinase species-mix experiment meta-data, related to Figures 1–5
Table S2. MoCAVI background gene programs, related to Figures 2–3
Table S3. MoCAVI salient gene programs, related to Figures 4–5
Table S4. icCITE-plex salient protein effect sizes, related to Figure 4
Table S5. Epigenetic inhibitor DOGMA-plex experiment meta-data, related to Figures 6–8
Table S6. DOGMA-plex salient protein effect sizes, related to Figure 6
Table S7. Motif-based TF groups, related to Figures 7–8
Table S8. TF motif–TF mRNA correlations, related to Figure 7.
Table S9. Permuted TF motif–TF mRNA correlations, related to Figure 7.
Table S10. Predicted TF–downstream gene regulation, related to Figure 7.
Table S11. TF–salient gene program associations, related to Figure 7.
Table S12. A list of antibodies, sgRNAs and primer sequences used in the study, related to Figures 1, 6 and 8.

### Supplementary Note 1: Comparison of salient-specific LFC estimation versus differential expression

We present the difference between differential gene expression, and our procedure for salient space-specific LFC (sssLFC) estimation. We also draw connections with previous work, justifying its utility for interpreting high-throughput screening data.

#### Building intuition about how sssLFC and DE differ

Differential expression analysis in scRNA-seq aims at assessing whether a given gene has significantly different expression between two sets of cells (*e.g.*, A and B). For generative models, such as scVI, this can be performed by repeatedly decoding the gene expression for different values of the latent variables, and calculating the log fold change (**Methods**). In the context of ContrastiveVI and MoCAVI, this is implemented by decoding the concatenation of the latent variables $(z, s)$, where $z$ is the background latent variable, and $s$ is the salient latent variable (**fig. S2D**, left).

The sssLFC, however, aims at assessing which genes are most affected by the salient variable $s$ for a unique set of cells A. We decode the gene expression for both $(z_a, s_a)$ and $(z_a, 0)$, and then compare how gene expression changes. Because we decode gene expression after altering the value of the latent variable, this estimate is a counterfactual estimate ("What if"). However, this presents the advantage of assessing contribution of the salient latent space, while controlling for variations within the background space (**Fig. S2D**, right).

#### Biological motivation and relationship to prior art

Calculation of the sssLFC is motivated by controlling for variation that occurs in the background data, and may not be the main focus for the screen. A common example is cell cycle or pre-existing cell subsets, which may be corrected for when analyzing Perturb-seq data [29].

Instead of correcting for these unwanted variations in a supervised way (for example, by identifying cell cycle as a signal to control for, and adding it as a covariate to the model;[29]), our model allows for unsupervised identification of background latent variables, and correction via the sssLFC estimate.

### Supplementary Note 2: Robustness of sssLFC estimation

We assess the robustness of our sssLFC estimation procedure. For ensuring lower runtimes, we performed these analyses on the smaller icCITE-plex kinase inhibitors dataset.

**Reproducibility of the estimates across technical replicates**

The kinase human data set includes several technical replicates for the non-stimulated condition. By design, the non-stimulated condition appears in three different plates, usually attributed to different concentrations of the chemical (here DMSO). We used this as a set of positive controls for which we expect perfect reproducibility between the sssLFCs. We therefore calculated sssLFCs for each of the three available DMSO concentrations (100 nM, 1 µM, and 10 µM) in unstimulated cells vs. stimulated vehicle, and verified that the results are highly reproducible between technical replicates (**Fig. 2.1**).

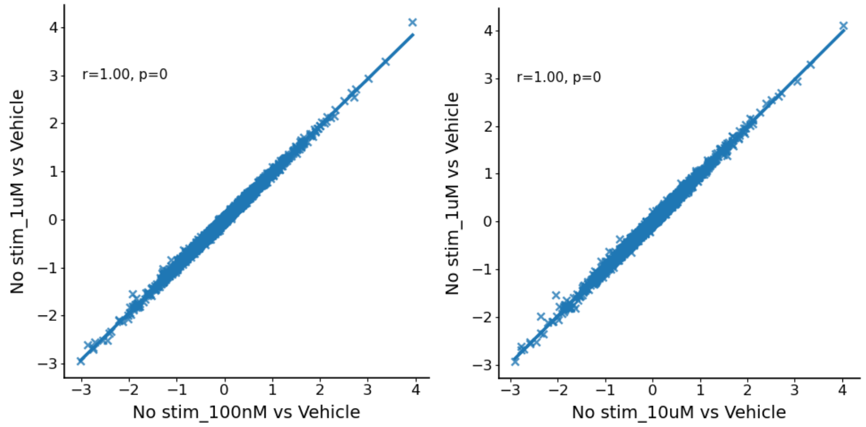


*Figure 2.1: Reproducibility of sssLFCs across two pairs of technical replicates on the kinase human data set (n=5,366 genes)*

**Assessment of sssLFCs and their significance with negative control samples**

Because we do not have ground-truth available for the value of sssLFCs, or their significance, we validated the method using negative control samples. In particular, we applied the procedure for estimating sssLFCs to samples from the baseline (stimulated) environment (Vehicle 100 nM). Because these cells are from the background, we should decode them with $s_n = 0$, and the sssLFCs should be exactly zero. However, in this assessment we decode them with $s_n$ from the variational distribution, and we would like to check that the sssLFCs are close to zero, and non-significant. We therefore first compared the values of the sssLFCs for Vehicle 100 nM with the ones from the non-stimulated condition (**Fig. 2.2**). As expected, the sssLFCs were small for the vehicle condition compared to the non-stimulated condition.

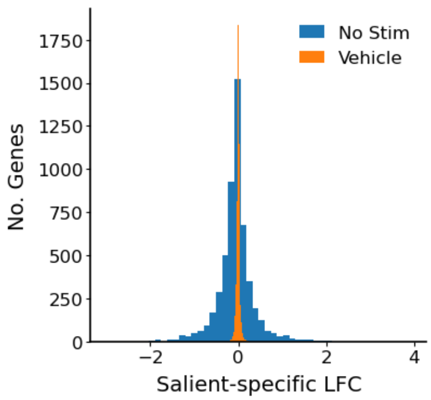


*Figure 2: Negative control assessment of ssLFCs on the kinase-human icCITE-plex data.*

Next, we assessed our criterion for statistical significance, by calculating the probability of sssLFC > 0.25 with the lvm-DE method, for each of the non-stimulated and vehicle control conditions. For the stimulated vehicle control condition, o genes were significantly changed under the 0.05 posterior expected FDR threshold, while for the non-stimulated condition, 256 genes were significant under the same threshold.

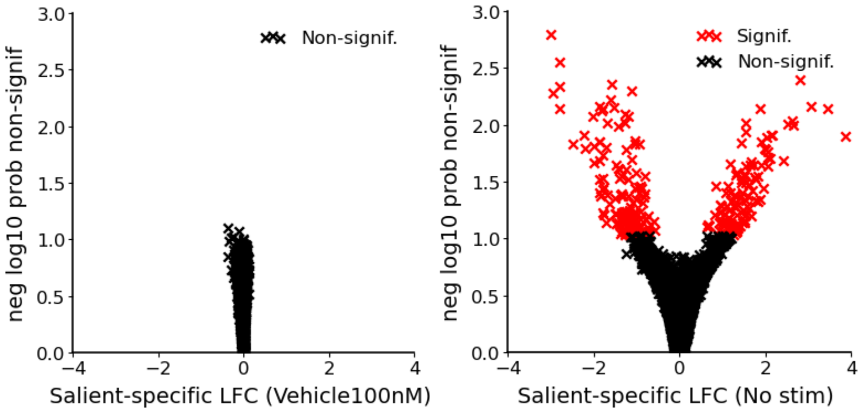


*Figure 3: Negative control assessment of significance for ssLFCs on the kinase human data set.*

**Robustness with respect to random initialization** The loss function of deep generative models is not convex with respect to the parameters, and the optimization procedure is not deterministic. Therefore, running the same training procedure several times may yield different sets of parameters. Because in the context of MoCAVI, the latent space could be changed, it is important to ensure that any counterfactual estimates are themselves unchanged across random seeds. We therefore assessed the correlation between the sssLFCs across two random initializations of the parameters, finding that the estimates are indeed reproducible across random seeds.

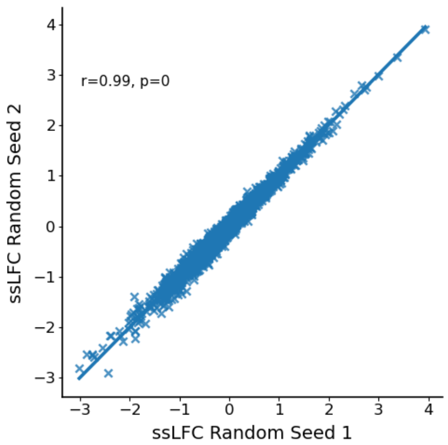


*Figure 4: Reproducibility of ssLFCs across two random seeds on the kinase human data set (n=5,366 genes)*
